# LUMI-lab: a Foundation Model-Driven Autonomous Platform Enabling Discovery of New Ionizable Lipid Designs for mRNA Delivery

**DOI:** 10.1101/2025.02.14.638383

**Authors:** Haotian Cui, Yue Xu, Kuan Pang, Gen Li, Fanglin Gong, Bo Wang, Bowen Li

## Abstract

The complexity of molecular discovery requires autonomous systems that efficiently explore vast and uncharted chemical spaces. While integrating artificial intelligence (AI) with robotic automation has accelerated discovery, its application remains constrained in fields with scarce historical data. One such challenge is the design of lipid nanoparticles (LNPs) for mRNA delivery, which has relied on expert-driven design and is hindered by limited datasets. Here, we introduce LUMI-lab, a self-driving lab (SDL) system that enables efficient learning with minimal wet-lab data by integrating a molecular foundation model with an automated active-learning experimental workflow. Through ten iterative cycles, LUMI-lab synthesized and evaluated over 1,700 LNPs, identifying ionizable lipids with superior mRNA transfection potency in human bronchial cells compared to clinically approved benchmarks. Unexpectedly, it autonomously uncovered brominated lipid tails as a novel feature enhancing mRNA delivery. *In vivo* validation further confirmed that inhalation of LNPs containing the top-performing lipid, LUMI-6, achieved 20.3% gene editing efficacy in lung epithelial cells in murine models, surpassing the highest efficiency reported for inhaled LNP-mediated CRISPR-Cas9 delivery in mice to our knowledge. These findings demonstrate LUMI-lab as a powerful, data-efficient platform for advancing mRNA delivery, highlighting the potential of AI-driven autonomous systems to accelerate innovation in material science and therapeutic discovery.

## 1 Introduction

The increasing complexity of molecular and material discovery necessitates the development of autonomous systems powered by artificial intelligence (AI) to navigate vast, uncharted molecular spaces with precision and efficiency. Self-Driving Laboratories (SDLs), which combine advanced robotic automation with data-driven experimental workflows, have emerged as powerful platforms for accelerating discovery [1, 2, 3]. By executing iterative design-make-test-analyze (DMTA) cycles autonomously, SDLs have demonstrated remarkable potential in areas with well-established datasets, such as solid-state materials [4, 5] and small molecules [6, 7]. In these studies, experimental design is typically guided by either existing language models (such as GPT-4 [8]), which have already learned sufficient domain knowledge, or newly developed machine learning (ML) models trained on extensive historical data of materials and chemical reactions.

However, the development of SDLs remains particularly challenging in emerging fields due to the scarcity of annotated historical data, which makes it difficult for models to learn and adapt to unexplored chemical spaces. The reliance on large-scale, annotated historical datasets and extensive computational tuning presents a significant obstacle in such fields. A prominent example of data scarcity is the development of delivery vehicles for nucleic acid drugs, such as mRNA. Following the success of Pfizer and Moderna COVID-19 vaccines [9, 10], mRNA therapeutics have rapidly emerged as one of the most promising and fastest-growing drug modalities, with lipid nanoparticles (LNPs) serving as a clinically-validated and safe delivery system [11, 12]. However, the development of new LNPs remains hindered by the limited availability of historical data as only three LNPs have received FDA approval to date [13]. In particular, the design of ionizable lipids in LNPs, crucial for mRNA encapsulation and endosomal escape, has been heavily relying on prior expert knowledge [14] due to the absence of comprehensive research data and systematic development platforms [15, 16].

To address the challenges inherent in emerging fields with limited historical data, we propose a two-step solution: First, a foundation model [17] can be pretrained with unsupervised learning to develop a generic understanding of molecular structures. This approach leverages the fact that, even in the absence of annotated datasets, the chemical space of interest can be enumerated in its raw form and enables the training of large-scale unsupervised ML models. Second, unlike conventional supervised-leaning models, the pretrained foundation model can perform few-shot learning, allowing it to adapt quickly with minimal ongoing wet-lab data. When integrated into an active learning framework, the model can be continuously optimized in a closed-loop experimental workflow, further enhancing its predictive accuracy and efficiency.

Here, we introduce LUMI-lab (**L**arge-scale **U**nsupervised **M**odeling followed by **I**terative experiments), an SDL system based on these principles. LUMI-lab incorporates large-scale molecular pretraining, active learning, and robotic automation to enable efficient exploration of new chemical space. It consists of two primary components: first, LUMI-model, a molecular foundation model pretrained on over 28 million molecular structures, which facilitates few-shot learning of new molecules with minimal experimental data; second, an automated, closed-loop experimental system, driven by LUMI-model predictions, that generates wet-lab data to iteratively optimize LNPs through ionizable lipid synthesis, LNP formulation, and high-throughput *in vitro* screening.

Through ten iterative cycles of active learning, LUMI-lab synthesized and evaluated more than 1,700 new LNPs, identifying numerous new ionizable lipids with superior mRNA transfection potency (mTP) in human bronchial epithelial cells compared to clinically approved benchmarks. Surprisingly, it autonomously uncovers **brominated lipid tails** as a previously unrecognized structural feature that enhances mRNA delivery efficiency, an insight not reported in prior literature to our knowledge. To validate these findings, we conducted *in vivo* experiments using LNPs formulated with the top six identified lipids. Five out of these candidates demonstrated mTP comparable to or exceeding that of SM-102, the commercial ionizable lipid used in Moderna’s COVID-19 vaccine. Furthermore, pulmonary delivery of CRISPR-Cas9 genome editing system using the best-performing LNP, formulated with our LUMI-6 lipid, achieved **20.3% epithelial cell editing efficacy in the lung**, surpassing the highest editing efficiency previously reported for LNP-mediated CRISPR-Cas9 delivery via inhalation [18].

These results highlight the capacity of LUMI-lab to autonomously identify novel molecular features, offering a powerful, data-efficient platform for advancing nucleic acid delivery. Beyond its application in LNP design, LUMI-lab exemplifies a transformative approach to autonomous molecular discovery, seamlessly integrating the adaptability of foundation models with the precision and high throughput of automated experimentation. By bridging computational and experimental innovation, it provides a versatile framework for addressing critical challenges in data-sparse fields. Its success in uncovering new ionizable lipid designs for mRNA delivery underscores its potential to accelerate innovation in material science and therapeutic discovery.

## 2 Main

### 2.1 Closed-loop optimization and autonomous lab for ionizable lipid engineering

LUMI-lab is a fully autonomous platform for molecular discovery, integrating molecular ML models, experimental automation, and active learning to efficiently explore vast, uncharted chemical spaces. Here, we demonstrate its capability in advancing the development of novel LNPs for mRNA delivery. LNPs typically consist of four key components: ionizable lipids, cholesterol, helper lipids, and PEGylated lipids. Among the key compositions of LNPs, ionizable lipids play the most essential in enabling efficient RNA encapsulation and facilitating endosomal escape, a crucial step in cytoplasmic mRNA release. These lipids dynamically transition between a neutral state at physiological pH, minimizing toxicity, and a protonated state in the acidic endosome, where they disrupt the membrane and release mRNA into the cytoplasm [19, 20, 21, 22]. Given their essential role in LNPs, this study focuses on using LUMI-lab to facilitate the discovery and optimization of ionizable lipids for enhanced mRNA delivery.

At the core of the system is the LUMI-model, a pretrained foundation model that serves as the computational “brain” of LUMI-lab (Figure 1a). In each experimental iteration, LUMI-model proposes candidate molecules for synthesis and testing, gradually prioritizing and identifying those with high mRNA transfection potency (**mTP**). The experimental process is supported by a suite of software modules, which manage laboratory communication, experimental orchestration, and hardware control (Figure 1b, Supplementary Figure S1). These modules integrate robotic operations with real-time data acquisition, providing a centralized, automated platform for workflow management. The hardware infrastructure includes two liquid handlers, one for ionizable lipid synthesis and the other for LNP formulation, along with a robotic arm, cell incubator, plate reader for assay measurements, and automated loading systems for pipette tips and 96-well plates (Figure 1c,d).

**Figure 1.**
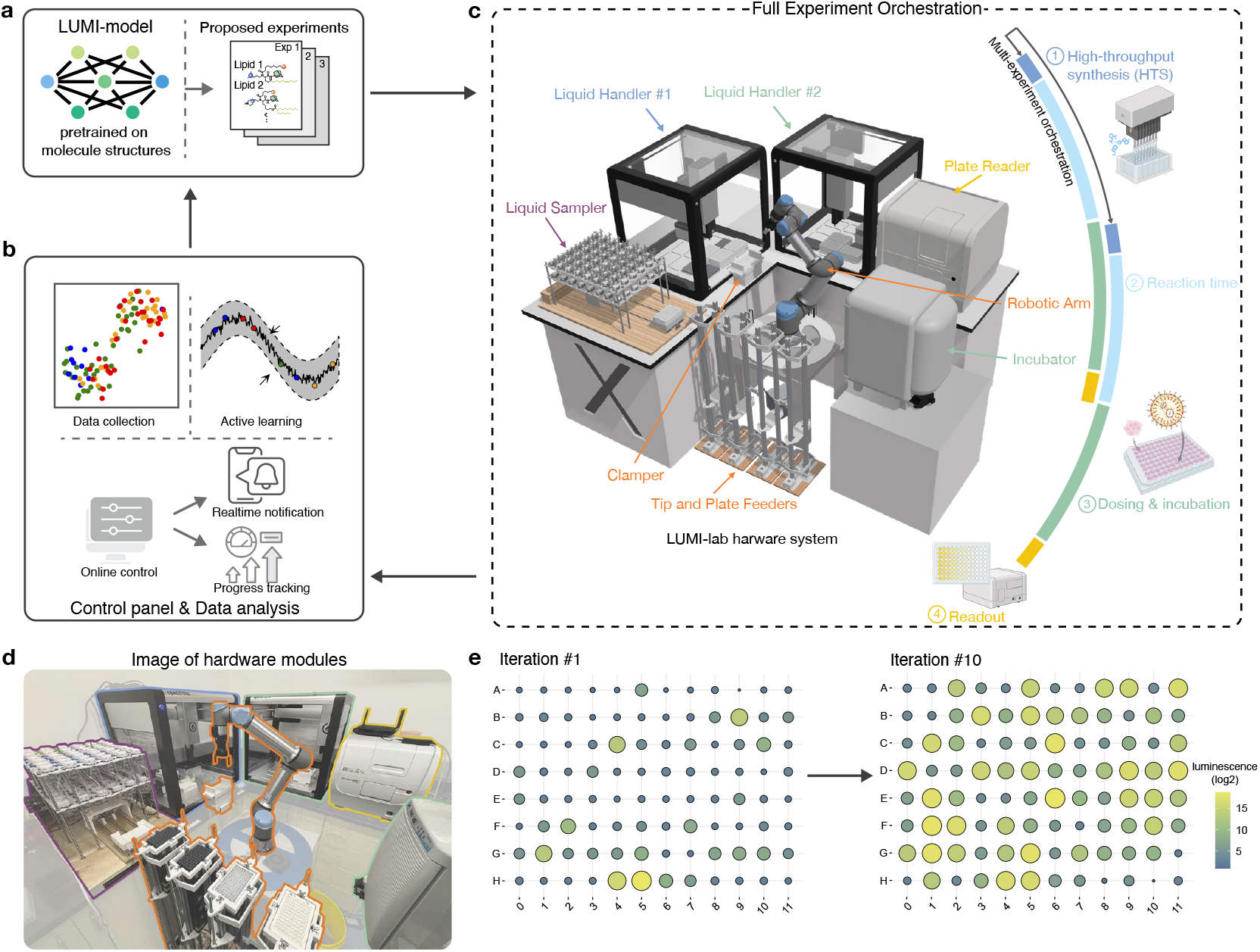
Overview of LUMI-lab, including **(a)** the molecular foundation model, LUMI-model, **(b)** software module of the control panel and online data analysis, and **(c)** hardware modules and the automatic experiments controlled by the orchestration module. Two experiments conducted simultaneously in an orchestrated manner are visualized by the colored curves. **(d)** The image of hardware modules shown in the physical layout. Each hardware equipment is outlined by the same color as in (c). **(e)** The relative log2 luminescence readout after dosing the Fluc mRNA-LNPs and 18-h cell incubation, for the proposed and tested LNPs in iterations 1 and 10, respectively. The increased luminescence intensity in iteration 10 indicates the higher luciferase expression in cells.

Each experimental iteration begins with LUMI-model proposing a set of chemically diverse ionizable lipid structures. These lipid candidates are then synthesized in a high-throughput manner by the automated liquid handler (Opentrons OT-2), which is programmed to precisely mix samples and perform controlled shaking. Following an 18-hour reaction, the resulting ionizable lipids are transferred to the second liquid handler and formulated into LNPs with helper lipids, cholesterol, PEG-lipids, and firefly luciferase (Fluc) mRNA, adhering to a classic LNP formulation ratio [23] (Methods). The 96-well plates containing cultured cells are then treated with mRNA-LNP samples using the second liquid handler, followed by an 18-hour cell incubation. After the incubation, the robotic arm retrieves the treated cell culture plates and transfers them back to the liquid handler, where a luciferase substrate reagent is added. The robotic arm then moves cell cultures to the plate reader, which measures the bioluminescence signal intensity to directly reflect Fluc mRNA transfection efficiency. This enables fast and quantitative mTP evaluation of LNPs with varying ionizable lipids. The data from each experimental cycle are processed automatically by the LUMI-lab’s software modules, which handle error correction, quality control, data normalization, and iterative updates to LUMI-model using active learning algorithms.

LUMI-lab’s iterative design enables continuous enhancement of predictive accuracy. The mRNA transfection data generated in each experimental cycle are used to fine-tune LUMI-model, improving its ability to evaluate and identify high-efficacy candidates. This closed-loop approach systemically integrates experimental insights for optimizing LUMI-model’s performance, thereby resulting in significant improvements in mTP across iterations (Figure 1e). Each experiment spans approximately 39 hours, primarily due to the reaction and cell incubation time. To maximize the experiment throughput, LUMI-lab incorporates an experiment orchestration module that manages concurrent tasks. For example, while one liquid handler doses cells during an ongoing experiment, the other initiates lipid synthesis for the next experiment (colored curves in Figure 1c, Supplementary Figure S2). This parallelized workflow substantially reduces the turnaround time, achieving a throughput far exceeding traditional manual methods.

### 2.2 A chemical foundation model designed for data-efficient active learning

LUMI-model, functioning as the computational engine of the system, is a foundation model trained on an extensive dataset of small molecule structures and further optimized for discovering ionizable lipids. LUMI-model is a transformer [25] based model designed to process two key inputs: atomic types and 3D atomic coordinates that define molecular conformations. By incorporating 3D structural information, the model develops a comprehensive understanding of physicochemical properties and atomic interactions, which are critical for tasks involving molecular structure analysis and optimization. To overcome the limitations in data-sparse fields, it employs a three-stage training pipeline: unsupervised molecular pretraining, continual pretraining, and supervised fine-tuning within an active learning framework. This progressive learning strategy equips the model with robust molecular representations, enabling rapid adaptation to nucleic acid delivery and broader molecular discovery applications.

The first stage of LUMI-Model employs self-supervised pretraining on a massive dataset of over 13 million molecular structures and their corresponding 3D conformations (Figure 2a, Methods). This stage uses a masked token prediction objective, where the model learns to reconstruct original atoms and denoise the atom 3D coordinates during training. Additionally, a contrastive learning approach is also integrated into the pretraining to improve the model’s ability to distinguish similar molecule structures (Methods). This combined approach enables the model to capture general chemical and spatial features that are broadly applicable across molecular design tasks.

**Figure 2.**
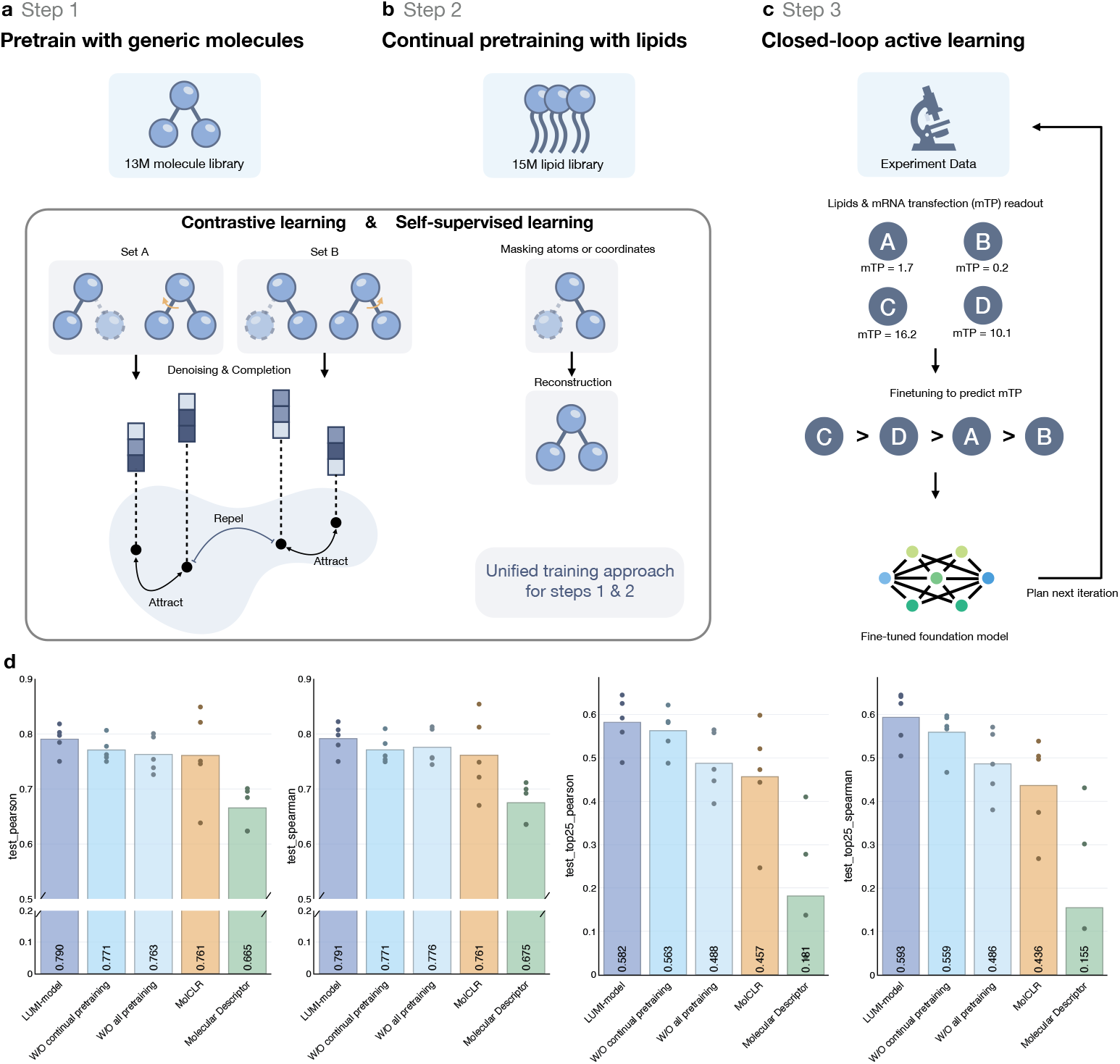
The LUMI-model training steps include **(a)** step 1. pretraining on a library of the 3D conformation of 13M generic small molecules, **(b)** step 2. continual pretraining on 15M lipid-like molecules, and **(c)** step 3. the closed-loop optimization via active learning during the self-driving lab experiment iterations. **(d)** Performance comparison of different model training on a pilot dataset of 1920 lipids, including the full LUMI-model, the LUMI-model without continual pretraining (step 2), the LUMI-model without all pretraining (stps 1 and 2), a commonly used pretrained GNN molecular model of MolCLR [24], and an MLP model with molecular descriptor input. Four metrics are shown, including Spearman and Pearson correlation coefficients on all data points and the coefficients on the top 25% lipids with the best experiment mTP results.

The second stage, continual pretraining, refines the model’s embeddings by focusing on a domain-specific dataset of lipid molecules (Figure 2b). This step adapts the general chemical representations learned in the first stage to the chemical space relevant to ionizable lipid design. Importantly, continual pretraining ensures that LUMI-model obtains generic molecular knowledge while prioritizing features most critical for LNP engineering.

The third stage involves supervised fine-tuning within an active learning framework (Figure 2c). In this stage, the model is incrementally trained on labeled experimental data from previous iterations of LUMI-lab to predict the mTP. The actual lipids proposed for the next experiment iteration will be selected based on the predicted mTP in an active learning manner (Methods). Later after the coming iteration, the predicted value will be matched with the actual on-board *in vitro* luminescence readout and fed into the next round of fine-tuning. This fine-tuning allows the model to incorporate experimental feedback and improve its ability to propose high-performing candidates for subsequent rounds of synthesis and testing. This three-stage process culminates in a versatile model capable of both generalizing across chemical spaces and excelling in targeted molecular design tasks.

To evaluate the effectiveness of LUMI-model’s pretraining and 3D inputs, we evaluated the fine-tuning of various models on a previously collected dataset containing the experimentally measured mTP of 1,920 LNPs with different lipids (Methods). In this test, LUMI-model is compared against other baseline models including (1) a variant of LUMI-model without the step 2 continual pretraining. (2) the LUMI-model without all pretraining of steps 1 and 2, (3) a commonly used graph neural network (GNN) model, MolCLR [24], and (4) a multi-layer perceptron (MLP) model taking only input of molecular descriptors [26]. We reported the Spearman and Pearson correlation between these models’ predictions and the actual experiment data on all lipids in the test set (Methods) and on the subset of lipids with top 25% mTP (Figure 2d). We first observed that all deep learning models with rich input of molecular structure information surpass the performance of the MLP model with only molecular descriptors. Particularly, the MLP model struggled on the predictions of the top 25% subset, with correlation coefficients less than 0.2. In comparison, LUMI-model and all its variants achieved higher scores than MolCLR, indicating the advantage of the 3D conformation input features. Within the comparison of LUMI-model variants, the full LUMI-model outperformed all alternatives, verifying the effectiveness of the first and second stage pretraining.

### 2.3 LUMI-lab finds new effective LNPs for bronchial epithelial cell transfection in ten iterations

We utilized LUMI-lab to conduct consecutive DMTA iterations to optimize LNPs for mRNA delivery to human bronchial epithelial (HBE) cells. Each iteration comprised two parallel experiments, synthesizing 184 lipid candidates across two 96-well plates (92 × 2), followed by LNP formulation and evaluation. Each plate included two wells treated by industry-standard LNPs containing DLin-MC3-DMA (MC3) and two non-treated wells as negative controls (Figure 3a).

**Figure 3.**
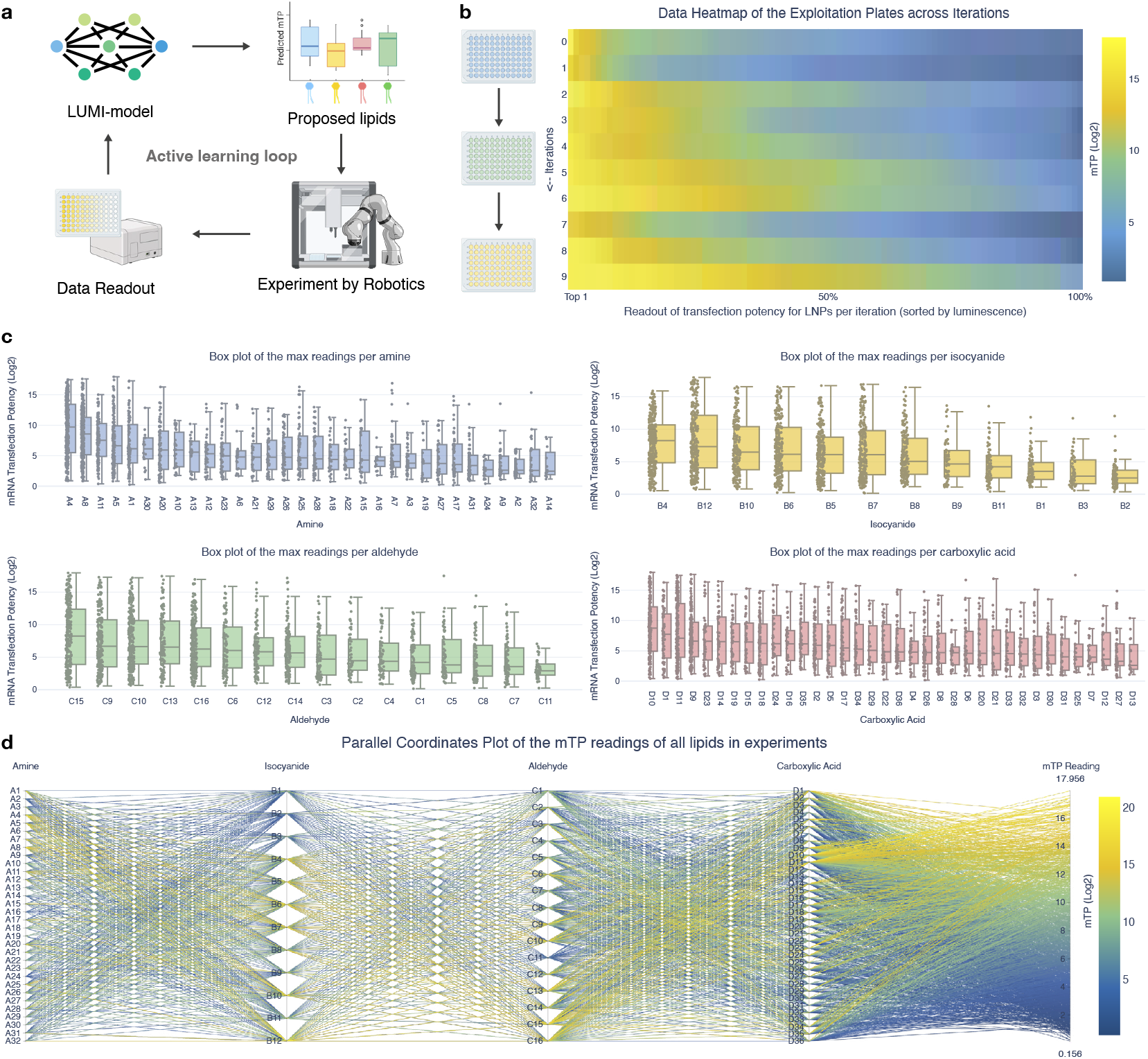
**(a)** Illustration of the iterations of experiments in LUMI-lab. **(b)** mTP readouts through ten experiment iterations. **(c)** Performance of four components in the combinatorial chemical library for the ionizable lipid synthesis, i.e. the amines, isocyanides, aldehydes, and carboxylic acids, respectively. **(d)** Component distribution and mTP readouts of all LNPs through experiments.

Unlike conventional active learning workflows, which primarily focus on improving predictive accuracy, LUMI-lab is required to simultaneously refine predictions and identify high-performing lipids within a finite number of iterations. To achieve this, each iteration follows a dual-plate strategy, optimizing top-performing candidates while maximizing information gain (Methods): (1) **Exploitation plate:** This first plate contains 92 ionizable lipid candidates predicted to exhibit high mTP, prioritized based on the latest LUMI-model predictions. (2) **Exploration plate:** The second plate contains 92 ionizable lipid candidates selected based on high ensemble model uncertainty. Lipids with the greatest variance across ensemble predictions were prioritized, ensuring systematic exploration of underrepresented regions in the chemical space. This exploitation-exploration framework enabled LUMI-lab to efficiently probe a broad ionizable lipid chemical space, ensuring both the discovery of superior lipid candidates and the continuous expansion of structure-activity relationship (SAR) knowledge. By iteratively improving predictions and expanding the diversity of evaluated lipid structures, LUMI-lab steadily advanced toward optimizing LNPs for mRNA delivery.

Throughout ten iterations, LUMI evaluated a total of 1,781 distinct lipid candidates, achieving substantial improvement in mRNA transfection efficiency. Early iterations prioritized refining the predictive accuracy of LUMI-model, as the experimental data revealed more SARs between molecular features and transfection performance. By the fourth iteration, the system identified several promising lipids that outperformed positive controls (Figure 3b). Notably, performance gains accelerated in later iterations, driven by the combined effects of refined exploitation and informed exploration.

The final iteration further demonstrated LUMI-model’s ability to consistently propose high-efficacy lipids. In the tenth iteration, over 50% of candidates exhibited superior transfection efficiencies (RLU *>* 10), highlighting the effectiveness of LUMI-lab’s optimization strategy (Figure 3b). Analysis of the ten iterations revealed key trends indicative of LUMI-model’s learning process. The exploitation plates consistently yielded high-performing candidates, while exploration plates expanded the diversity of chemical features captured by the model. By later iterations, LUMI had converged on specific headgroups and linker structures (Supplementary Figure S3) associated with high transfection potency (Figure 3c,d), particularly: N-(2-Aminoethyl)piperazine (A4), Tris(2-aminoethyl)amine (A8), and 1-Isocyanoadamantane (B12). These components were likely key to enhancing lipid packing and facilitating endosomal escape, critical factors for effective mRNA transfection [27, 28, 29]. These findings underscore the importance of integrating exploration into active learning workflows, particularly in sparse and underexplored chemical spaces, where expanding structural diversity is essential for identifying novel, high-performance molecules.

### 2.4 A new family of effective ionizable lipids with brominated tails discovered by LUMI-lab

Throughout the active learning and fine-tuning process, LUMI-model has progressively learned the relationship between molecular features of ionizable lipids and their mRNA transfection performance. This relationship can be visualized through UMAP projections of lipid embeddings generated by the optimized LUMI-model after the tenth iteration. In the UMAP of the entire 221k synthesizable lipid chemical space (Figure 4a), lipids with the highest and lowest predicted mTPs form distinct local clusters, reflecting well-captured SARs. Notably, one of the most prominent clusters corresponds to lipids with brominated tails (Figure 4b). Further analysis reveals that this brominated lipid cluster is enriched in lipids with globally high predicted mTP and low prediction variance (Figure 4b, zoom-in panels), suggesting a strong correlation between bromine modification and mRNA transfection performance.

**Figure 4.**
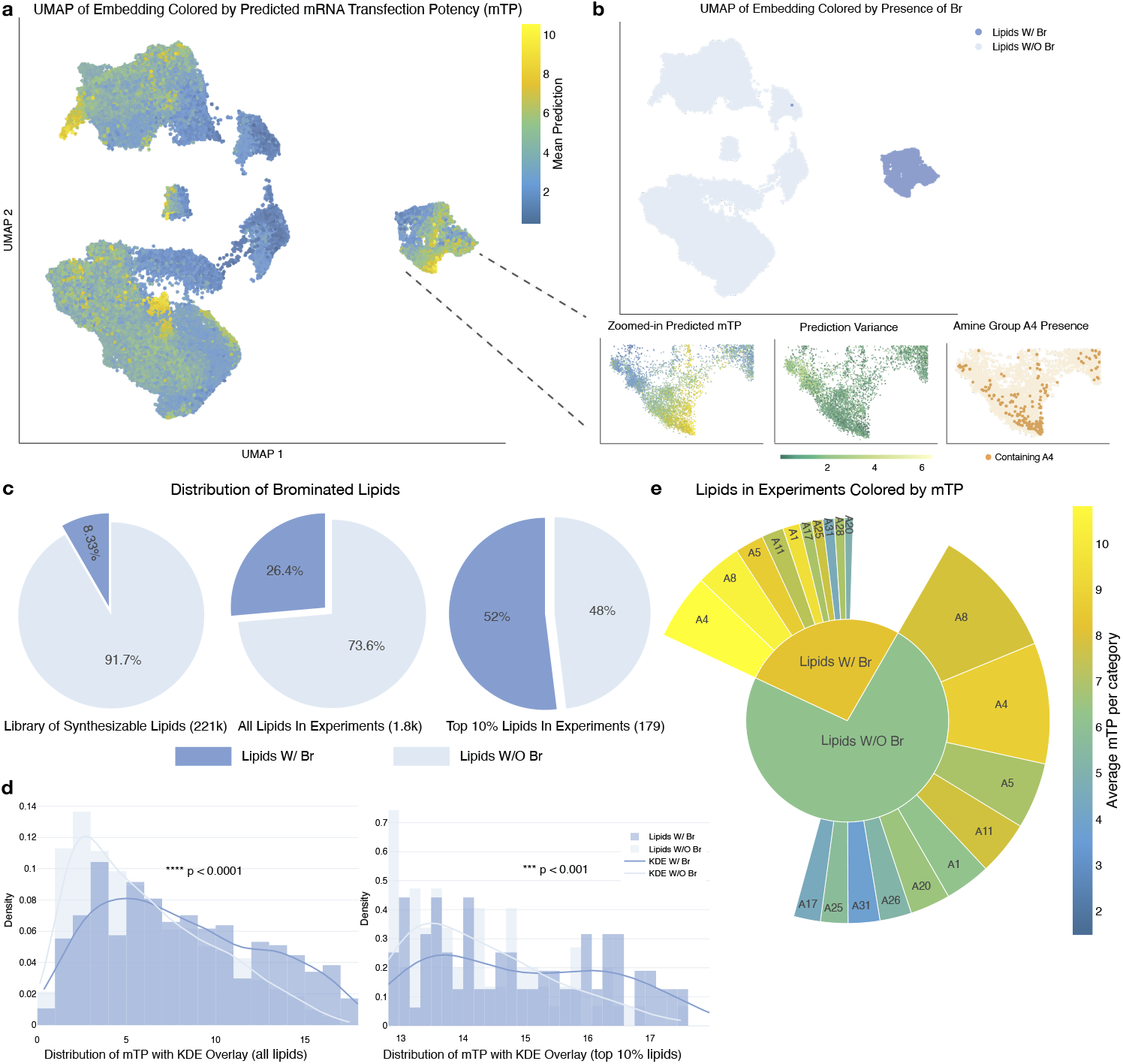
**(a)** UMAP of all synthesizable lipids (221k) composed of the designed Ugi-4CR library. Embeddings extracted from the fine-tuned model at the final iteration. Colored by the average mTP predicted by the ensemble of five models. **(b)** (Top) UMAP of all synthesizable lipids (221k) colored by whether lipids contain bromine atoms. (Bottom) Zoom-in panels showing the predicted mTP, the variance of predictions in the model ensemble, and the presence of the A4 component in the subcluster of brominated lipids. **(c)** Distribution of brominated lipids in the synthesizable chemical space, in actual experiments, and in the top 10% lipids in experiments. **(d)** mTP distributions of all lipids in experiments (left) and top 10% lipids in experiments (right). The overlay curves show the kernel density estimation (KDE) **(e)** Pie chart of lipids in experiments, categorized by whether the lipid contains Bromine (inner) and by the headgroup in the lipid (outer). The ten most common headgroups are shown in the diagram for brominated lipids and non-brominated lipids, respectively.

Beyond identifying brominated lipids as a distinct structural feature in the embedding space, LUMI-model actively prioritized these lipids in experimental iterations. Despite accounting for only 8.33% of the 221k synthesizable lipids, bromine-tail lipids constituted 26.4% of all candidates proposed by LUMI-model for synthesis and testing (Figure 4c). More strikingly, 52% of all top-performing lipids (top 10% mTP) across experiments were brominated, confirming their impactful contribution to enhanced mRNA transfection efficiency.

The superior performance of brominated lipids was further validated through a comparative analysis with non-brominated lipids. Brominated lipids exhibited significantly higher expected mTP values (*p <* 0.0001) and a greater proportion of high-mTP lipids (Figure 4d, left). This advantage remained consistent even among the top-performing 10% of lipids (*p <* 0.001) (Figure 4d, right). However, while bromination contributes substantially to mTP, it is not the sole determinant; other molecular features, particularly headgroups and linkers also play important roles. Interestingly, brominated and non-brominated lipids across the LUMI-lab experiments share similar headgroup compositions (Figure 4e) with high-performing headgroups such as A4 and A8 present in both categories. Therefore, the superior mTP of brominated lipids, even among those with similar headgroups, highlights bromination as a distinct structural feature that enhances mRNA delivery efficacy.

### 2.5 Enhanced *in vivo* mRNA delivery and gene editing potential of brominated ionizable lipids

While LUMI-lab successfully identified high-mTP lipids and uncovered brominated tails as a key feature enhancing LNP performance in *in vitro* studies, we sought to validate the functional consistency of these findings in complex physiological environments. To this end, we conducted an *in vivo* study of pulmonary mRNA delivery, evaluating the top-performing lipids from LUMI-lab’s final iteration. Specifically, the six highest-mTP lipids were selected, all of which interestingly contained brominated tails (highlighted in red) (Figure 5a). LNPs encapsulating Fluc mRNA were formulated with each ionizable lipids candidate, respectively (Figure 5b). The characterization of the nanoparticle size and polydispersity index (PDI) confirmed that all six new LNPs formed stable complexes with known helper lipid systems and successfully encapsulated Fluc mRNA (Figure 5c,d) [18]. LNPs were administrated into mouse lungs via intratracheal (I.T.) injection, and the IVIS images of the lungs were captured six hours post-injection to assess *in vivo* mRNA transfection efficiency. Notably, five candidates exhibited mRNA delivery efficiency comparable to or exceeding SM-102, the industry-standard lipid used in Moderna’s COVID-19 mRNA vaccine, while the least effective candidate still matched the performance of MC3, another FDA-approved ionizable lipid. (Figure 5e,f). To assess whether the enhanced mRNA transfection efficiency of LUMI-6 was attributable to bromine incorporation, we synthesized LUMI-6D, its debrominated derivative, which differs only by a single bromine atom (Supplementary Figure S4a). LNPs were formulated under identical conditions to ensure experimental consistency. Bioluminescence assays in HBE cells revealed that LUMI-6 exhibited 1.8-fold higher mRNA transfection efficiency than LUMI-6D, directly confirming that bromine incorporation plays a critical role in enhancing mRNA delivery (Supplementary Figure S4b). No significant difference in cytotoxicity was observed between LNPs formulated with brominated and non-brominated ionizable lipids (Supplementary Figure S5). These results manifest brominated tails as a key structural feature that improves LNP-mediated mRNA transfection, aligning closely with LUMI-model predictions.

**Figure 5.**
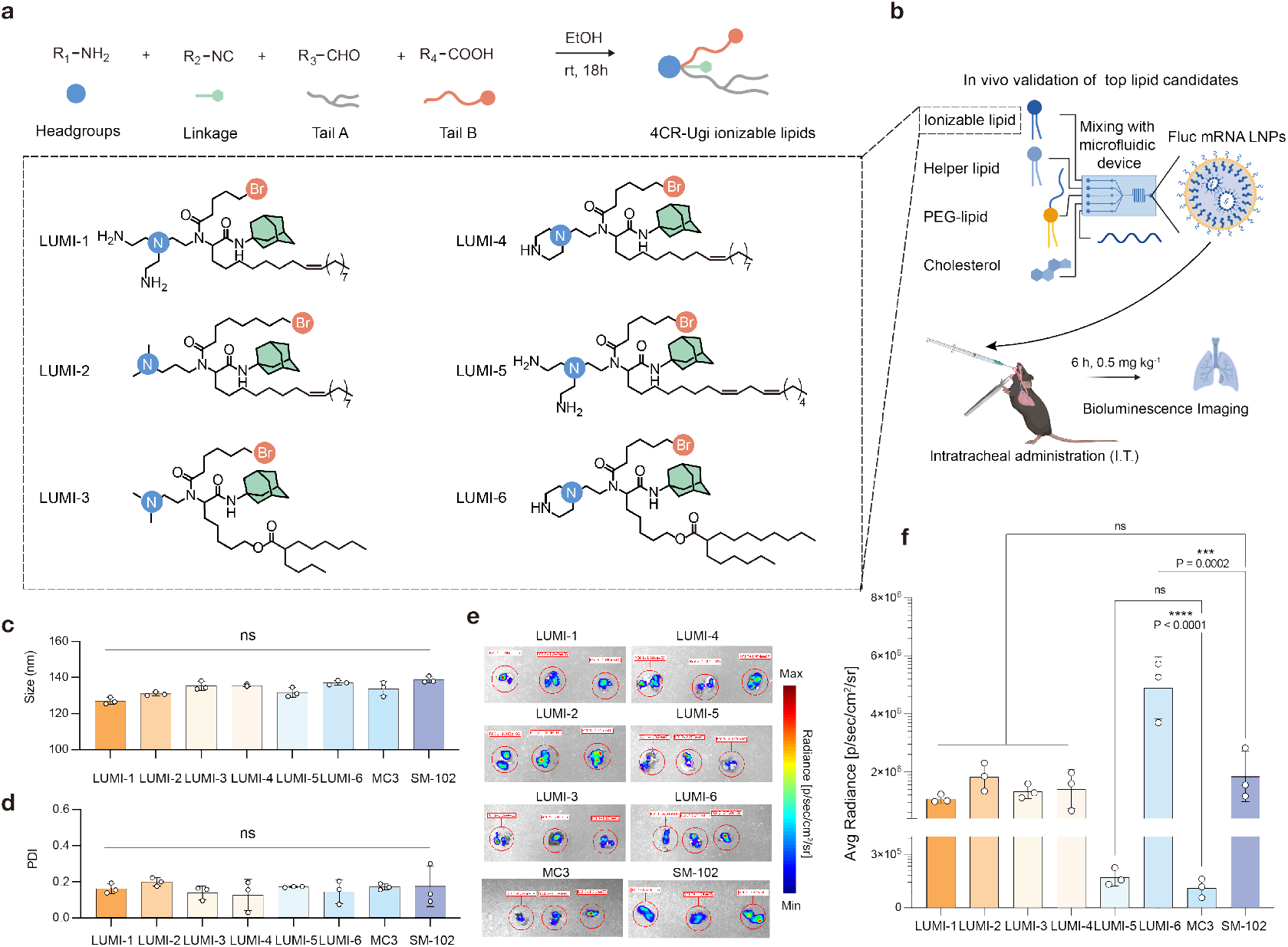
**(a)** Top lipid candidates selected by LUMI, accompanied by a diagram illustrating the four key components of ionizable lipids synthesized via Ugi-4CR. **(b)** Formulation of LNPs by combining the top lipid candidates with other lipids, followed by I.T. injection for functional validation. **(c**,**d)** Physical properties of the LNPs, including size (nm) and Polymer Dispersity Index (PDI). **(e**,**f)** *In vivo* imaging system (IVIS) results showing luciferase expression in mouse lungs mediated by LNPs formulated with different lipid candidates. Data are shown as mean ± s.e.m. (*n* = 3 biologically independent animals). Statistical significance evaluated using two-way ANOVA (\**P <* 0.05; \*\**P <* 0.01; \*\*\**P <* 0.001; \*\*\*\**P <* 0.0001).

Building on these findings, we further investigated LUMI-6, the best-performing lipid from the above pulmonary Fluc mRNA delivery study, to assess its potential for CRISPR-Cas9 gene editing in the lung. LUMI-6 LNPs encapsulating Cas9 mRNA/gRNA complexes were formulated using a previously optimized ratio [18] and administered via I.T. instillation into Ai9 reporter mice (Figure 6a) [15, 30]. *In vivo* IVIS imaging revealed that LUMI-6 LNP significantly out-performed SM-102 LNP, demonstrating superior editing efficiency in the lung tissue (Figure 6b). Flow cytometry analysis of tdTomato-positive cells further showed that LUMI-6 LNP achieved a gene editing efficiency of 20.3% in lung epithelial cells, a substantial improvement over SM-102 LNP (Figure 6c). To assess the distribution of CRISPR-Cas9 editing mediated by LUMI-6 LNPs across cell subtypes, we performed immunofluorescence staining of ciliated cells (a-tubulin^+^) and club cells (CCSP^+^), two important airway epithelial populations implicated in lung diseases such as cystic fibrosis. The results confirmed efficient gene editing in both cell types, thus reinforcing LUMI-6’s potential for gene editing in the airway epithelium (Figure 6d).

**Figure 6.**
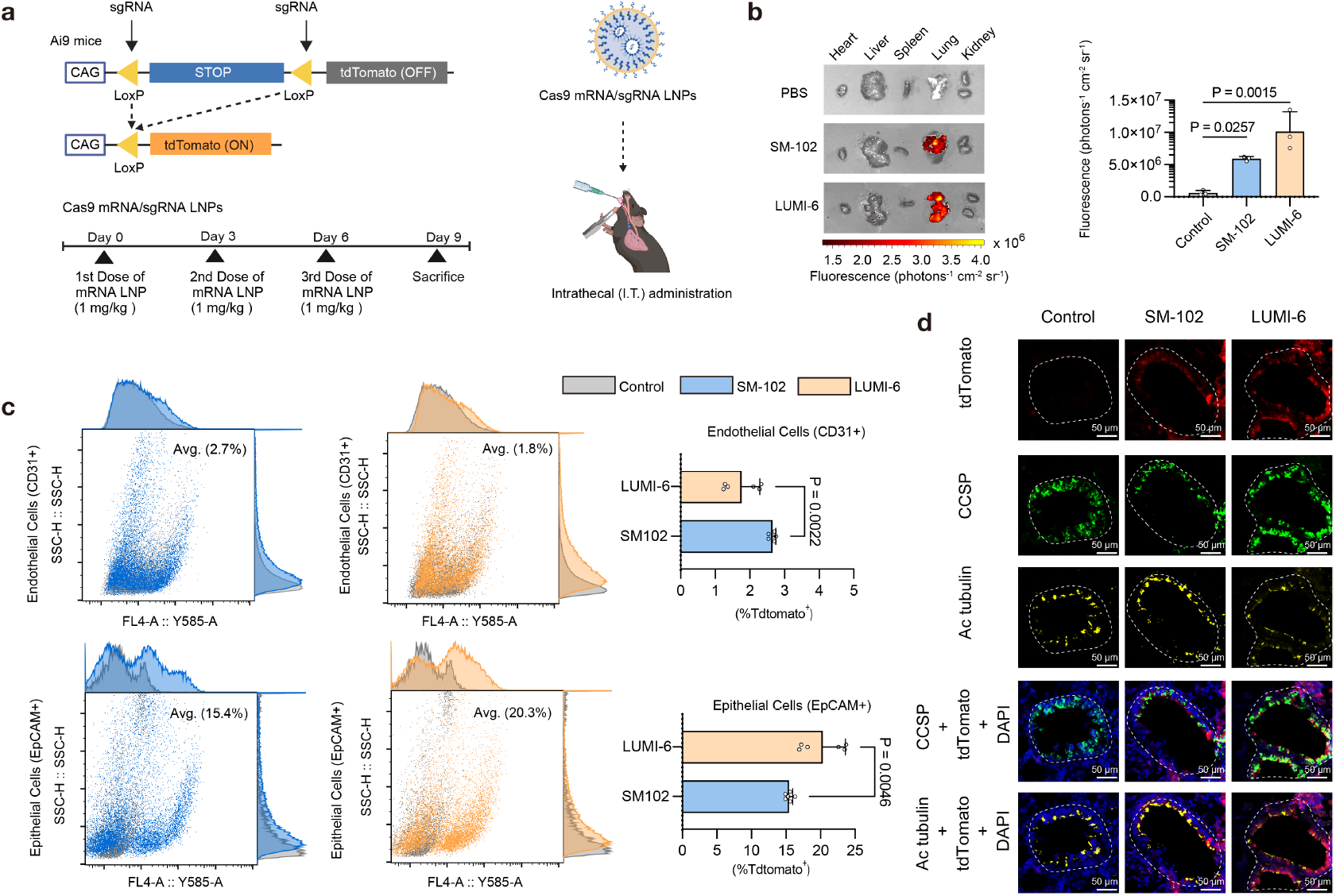
**(a)** Ai9 mice were used to evaluate LNP-mediated CRISPR/Cas9 gene editing *in vivo*. A CRISPR Cas9 gene editing strategy is introduced to trigger the disruption of stop signals and turn on of tdTomato expression in the Ai9 reporter mice. Three days after the last dose, the lungs were collected and analyzed by flow cytometry (*n* = 3 mice for each group). **(b)** Representative IVIS images of mouse lungs at 6 h following I.T. administration of SM-102 or LUMI-6 mLuc-LNPs (1 mg/kg mLuc per mouse). **(c)** Flow cytometry analysis of Tdtomato^+^ cells in lung endothelial and epithelial cells from Ai9 mice treated with Cas9 mRNA and sgRNA-formulated LNPs (dose: 1 mg/kg, *n* = 3 biologically independent animals and *n* = 2 samples for each mice). **(d)** Representative native immunofluorescence images of lung sections, including club (CCSP^+^) and ciliated cells (Ac tubulin^+^). Statistical significance evaluated using a two-tailed unpaired student’s *t*-test (\**P <* 0.05; \*\**P <* 0.01; \*\*\**P <* 0.001; \*\*\*\**P <* 0.0001).

## 3 Discussion

A fundamental challenge in molecular and material discovery is the lack of systematic strategies to efficiently explore vast, uncharted chemical spaces, particularly in fields with limited historical data. Traditional experimental approaches rely heavily on expert-driven design and extensive trial-and-error methods, which inherently slow the pace of innovation. This work introduced LUMI-lab, an AI-driven autonomous laboratory, to overcome these barriers by integrating foundation model pretraining, high-throughput robotic experimentation, and closed-loop active learning. Through LUMI-lab, we conducted ten iterative experimental cycles, synthesizing and evaluating over 1,700 ionizable lipids. This led to the discovery of LUMI-6, an ionizable lipid with superior mRNA transfection efficiency in lung epithelial cells, showcasing the power of AI-guided molecular discovery for RNA delivery.

A particular notable finding of this study is LUMI-lab’s autonomous identification of brominated lipid tails as a novel structure feature that enhances mRNA delivery, an insight previously unrecognized in expert-driven LNP design. This discovery underscores the power of AI-guided autonomous platforms to reveal new fundamental design principles beyond human intuition. Bromine atoms are widely used in medicinal chemistry to modulate key physicochemical properties, such as lipophilicity, membrane permeability, and metabolic stability. For instance, bromination has been shown to enhance the blood-brain barrier penetration of Pheniramine, leading to the development of Brompheniramine [31], while the incorporation of bromine improved the corneal penetration of Bromfenac[32]. Despite its well-established role in small-molecule drug development, bromine’s potential in lipid engineering for nucleic acid delivery has remained largely unexplored [33, 34]. Our findings indicate that brominated lipid tails consistently enhanced LNP-mediated mRNA transfection efficiency, as validated both *in vitro* and *in vivo*. This suggests that bromine substitution optimizes key molecular interactions, potentially influencing lipid packing, membrane fusion, and endosomal escape efficiency. The fact that LUMI-lab autonomously prioritized brominated lipids over iterations highlights its ability to recognize and exploit new design rules, refining its chemical intuition over time. Further studies are needed to elucidate the precise molecular mechanisms underlying bromine’s effects and to determine whether alternative halogen modifications such as fluorination, chlorination, and iodination could yield similar or superior enhancements in mRNA delivery efficiency.

In addition to remarkable mTP, LUMI-6 LNP demonstrated exceptional efficiency in pulmonary delivery of CRISPR-Cas9 gene editing tools, achieving 20.3% gene editing efficiency in lung epithelial cells, the highest reported editing efficiency for inhaled LNP-mediated CRISPR-Cas9 delivery in mice to our knowledge. This result is intriguing given the limited number of FDA-approved LNPs and the scarcity of autonomous platforms for developing nucleic acid delivery systems. Interestingly, LUMI-6 exhibited a preference for transfecting lung epithelial cells over endothelial cells compared to SM-102. This selectivity is relevant for gene therapy targeting many lung congenital diseases such as cystic fibrosis, alpha-1 antitrypsin deficiency, and surfactant protein disorders, where precise gene correction in airway epithelial cells is critical. Further optimization efforts can be conducted by LUMI-lab to refine LNP composition, enhancing cellular tropism while maintaining high mRNA transfection efficiency and minimizing off-target effects.

Beyond its immediate impact on ionizable lipid discovery, LUMI-lab represents a scalable and generalizable framework for molecular engineering. At its core, the foundation model, LUMI-model, which leverages unsupervised pretraining on 28 million molecular structures, enables few-shot learning and rapid adaptation to previously unexplored chemical spaces. The incorporation of 3D molecular representations and coordinate-based contrastive learning further enhanced the model’s prediction accuracy, as demonstrated in our comparative analyses. These improvements highlight the importance of holistic structure representations in molecule modeling, allowing LUMI-model to efficiently assimilate data from ongoing autonomous experiments. This continuous learning capability establishes LUMI-lab as a powerful tool for the co-evolution of large-scale molecular models and SDLs.

While this study demonstrates the potential of LUMI-lab, several areas warrant further exploration. Incorporating human organoid screening into the workflow might bridge the gap between *in vitro* and *in vivo* testing, providing a more physiologically relevant platform for evaluating LNP performance and predicting *in vivo* efficacy in human patients. Expanding the range of assay readouts, such as stability, biodegradability, and immunogenicity, would also provide a more comprehensive evaluation of LNP candidates. From a modeling perspective, LUMI-model could be extended beyond pre-defined chemical libraries to actively generate new molecular designs, enabling iterative refinement of lipid structures based on experimentally validated backbones. By integrating structure-based optimization with active learning loops, LUMI-lab could continuously propose new lipid architectures tailored for specific targets, further advancing rational lipid engineering and precision medicine.

In conclusion, LUMI-lab establishes a scalable and adaptive framework for tackling data-sparse challenges in emerging fields by integrating advanced machine learning with autonomous experimentation. Its success in uncovering novel structural features of ionizable lipids for mRNA and gene editor delivery demonstrates its promising potential to uncover new structure-activity relationships and drive the development of next-generation nucleic acid delivery systems. Looking ahead, the principles and platform technologies demonstrated in this work could extend beyond LNPs to inform discoveries in material science, biomolecular engineering, and therapeutic development, paving the way for AI-driven breakthroughs across diverse disciplines.

## 4 Methods

### 4.1 LUMI-model architecture

LUMI-model is a large-scale molecular representation model that builds on advances in 3D molecular pretraining frameworks. LUMI-model leverages a 3D transformer architecture that has been proposed for modeling small molecules [35, 36], and expands on it with training procedures and objectives tailored to ionizable lipid engineering. The backbone of the model employs a 15-layer transformer architecture [25] augmented to handle 3D spatial data effectively. The input to the model contains atom types and atom coordinates in 3D space, typically representing a molecule conformation.

#### 4.1.1 Input embeddings

For an input molecule containing *N* atoms, the 3D structural information is composed into two parts: (1) The atom types are denoted as a list of integer identifiers {*x*_1_, *x*_2_, …, *x*_*N*_}, *x*_*i*_ ∈ ℕ. Each integer denotes an atom element, such as Carbon and Nitrogen. (2) The pairwise distance matrix between all atoms, ***D***_*x*_ ∈ ℝ ^*N ×N*^.

We applied pytorch embedding layers to encode the atom type *x*_*i*_ into a learnable vector of 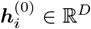, where *D* is the dimension of the embedding features, and the superscript (0) denotes it as the embedding before the first transformer layer. The combined embedding matrix, 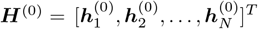, will be input to the stacked transformer layers in LUMI-model.

To encode the distance matrix, an atom type aware Gaussian Kernel method [37] was used to process each distance record *d*_*i,j*_ between atoms *i* and *j* from ***D***_*x*_ into a pairwise interaction matrix ***S***^(0)^ ∈ ℝ^*N ×N*^ between atoms *i* and *j*. Notably, these encodings are only dependent on the pairwise distances, and thus ensure the model is invariant to global rotations and translations, making it particularly suitable for handling the 3D conformation of molecules.

#### 4.1.2 Self-attention for molecular representation learning

For the self-attention between atom embedding vectors, the Query, Key, and Value matrices at *l*-th layer are first computed as in standard multi-head self-attention [25]. These matrices, denoted as 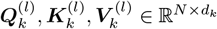, are learnable transformations from the input embeddings ***H***^(*l*−1)^. *d*_*k*_ is the embedding dimension for the *k*-th attention head, and 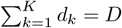.

The attention layer is enhanced to incorporate the atom coordinates information encoded in the pairwise distance representation, which was originally introduced by Uni-Mol [35]. Specifically, in the *l*-th (*l* ∈ [1, 15]) transformer layer, the self-attention per head *k* is computed as

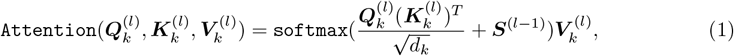

where a standard softmax operator [38] is used, and the interaction matrix per layer ***S***^(*l*)^ is computed as

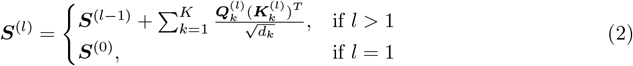

The output embedding of the *l*-th layer, ***H***^(*l*)^ is then generated by consecutive computations of the concatenation of multi-head attentions, a feed-forward neural network, and layer normalization operation [39].

### 4.2 Model Training

LUMI-model employs a three-stage training workflow to maximize its adaptability and performance:

#### 4.2.1 Step 1. Unsupervised pretraining on generic molecules

The initial unsupervised pretraining phase is conducted on a comprehensive molecular dataset of 13,369,320 distinct molecules with 147,062,520 conformations. This extensive dataset consists of commercially available molecules, originally released by Uni-Mol [35]. To enhance data quality and reduce redundancy, we apply deduplication, ensuring a structurally diverse and high-quality molecular space that optimizes LUMI-model’s representation learning.

LUMI-model is trained using three complementary learning objectives: (1) masked atom prediction, (2) 3D positional recovery, and contrastive representation learning. In the first task, a subset of atomic identities in the input molecule is randomly masked, and the model learns to reconstruct the missing atomic properties. In the second task, Gaussian noise is added to the atomic coordinates, and the model is trained to predict the original 3D positions of perturbed atoms. These first two objectives follow the settings in Uni-Mol [35].

In addition to these objectives, we introduce (3) contrastive learning to improve conformation-aware molecular embeddings. For each mini-batch of training examples, we generate two augmented versions of the same molecule by applying independent atom masking and coordinate perturbations. These augmented versions serve as positive pairs, while embeddings of different molecules within the mini-batch are treated as negative examples. The model is trained using the normalized temperature-scaled cross-entropy loss (NT-Xent), following the SimCLR framework [40]. The NT-Xent loss is defined as:

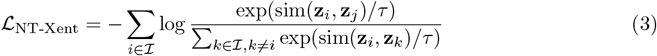

where **z**_*i*_ and **z**_*j*_ represent the embeddings of two augmented versions of the same molecule, sim(**z**_*i*_, **z**_*j*_) denotes the cosine similarity between the two embeddings, *τ* is a temperature scaling parameter, and ℐ is the set of all samples in the minibatch. This loss encourages the model to bring representations of different conformations of the same molecule closer while pushing apart those of different molecules.

#### 4.2.2 Step 2. Continual pretraining on lipid-like molecules

Following initial unsupervised pretraining, the model further undergoes continual pretraining on a domain-specific dataset of lipid-like molecules, employing the same learning objective as unsupervised pretraining. This specialized dataset includes 15,491,072 unique lipid-like molecules with 170,401,792 distinct conformations. The details of constructing this lipid-focused corpus are provided in Section 4.2.4.

To avoid catastrophic forgetting [41] during the continual pretraining stage, a phenomenon where the model loses its learning from the previous stage, we also sample data from the prior stage and utilize a lower learning rate during training (Supplementary Note S.1). This step adapts the general chemical knowledge gained in the first stage to the specific chemical space of ionizable lipids, ensuring the model prioritizes features most relevant to nucleic acid delivery applications.

#### 4.2.3 Step 3. Fine-tuning with closed-loop active learning

The final training stage involves fine-tuning in a supervised manner using experimental data generated by LUMI-lab. The model predicts the potential mTP of the lipids, optimized to match the onboard mTP readouts. Fine-tuning is performed iteratively, with the model updated after each experimental cycle to incorporate new data and refine its predictions. For each round of the experiment, we finetune an ensemble of 5 models on 5-fold cross-validation. The ensemble improves prediction robustness and enables uncertainty quantification with prediction variance in the active learning framework.

LUMI-model operates within a closed-loop active learning framework, enabling it to improve predictions and optimize lipid proposals iteratively. At each iteration, the model generates a candidate pool of lipids for LUMI-lab. To balance exploitation and exploration, two sets of candidates are selected for synthesis and testing: (1) lipids predicted to have the highest mTP and (2) lipids with the highest uncertainty in ensemble predictions, maximizing the information gain in subsequent iterations.

To effectively explore the vast pool of 221K candidates, we design a scalable sampling policy tailored to the foundation model. Initially, we quantify prediction uncertainty using ensemble prediction variance. Most uncertain lipid candidates often share similar chemical properties, such as having the same ionizable headgroups. Sending all such molecules for synthesis would inefficiently utilize experimental resources and fail to diversify the training data, as many samples would be overly redundant. Thus, we propose a strategy of diversity-aware sampling during the exploration stage.

Inspired by Citovsky et al. [42], we implement a diversity-aware sampling policy. First, we identify the *N* = 10, 000 (∼5% of the pool) most uncertain lipids based on ensemble variance. These lipids are then clustered using molecular embeddings generated by the foundation model, which captures the structural and chemical diversity of the molecules. To ensure a balanced representation across clusters, we employ a round-robin sampling strategy, selecting candidates from each cluster in a cyclical manner. This approach ensures both high uncertainty and molecular diversity in the exploration set. Additionally, we apply a limit for the number of repeated chemical reagents (component in the Ugi-4CR reaction) used in each experiment. This intends to first encourage the diversity of molecules again, and avoid stock management challenges by overusing of few specific reagents. The maximum number of usage per reagent is set to 35 for each experiment.

#### 4.2.4 Datasets for training stages

##### Construction of in-domain lipid-like molecule dataset

To further align LUMI-model’s capability for lipid engineering, we also construct an expanded lipid-like molecule dataset for continual pretraining. Following the Ugi-4CR combinatorial chemistry used for synthesizing lipids in SDL, we generate the dataset by systematically enumerating various options for each chemical component. Based on the coverage of chemical properties and our empirical experience, we select 64 amines, 62 isocyanides, 64 alkyl aldehydes, and 61 alkyl carboxylic acids for enumeration (see Supplementary Note S.2 for complete list), constructing 15,491,072 distinct lipid-like molecules in total.

To generate the 3D conformations for each molecule, we follow a similar conformation generation pipeline used in constructing a dataset for unsupervised pretraining 4.2.1. We use ETKGDv3 [43, 44] with RDKit [45] to propose conformation and apply Merck molecular force field (MMFF) to optimize the generated conformation [46, 47]. For each of the molecules, 10 3D conformations are generated along with an additional molecular graph, resulting in a total of 11 conformers per molecule.

##### Combinatorial chemical library for the closed-loop experiments

Our virtual library for screening ionizable lipids consists of 221,184 lipids, covering 32 Amines, 12 Isocyanides, 36 Lipid Aldehydes, and 16 Lipid Carboxylic, the detailed chemical library can be found in Supplementary Figure S3. The construction of this virtual library is similar to the method we discussed in 4.2.4.

### 4.3 Design of automatic experiment

Each experimental iteration begins with LUMI-model proposing a diverse set of ionizable lipid candidates, balancing predicted mTP and chemical diversity to ensure both optimization of high-performing lipids and exploration of new chemical space. The selected lipids are synthesized using an automated liquid handling system (Opentrons OT-2), which precisely dispenses reagents, mixes components, and performs controlled shaking to initiate the reaction. The synthesis process continues for 18 hours, allowing the ionizable lipids to form under standardized conditions. Once the reaction is complete, the synthesized lipids are transferred to a second liquid handler, where they are formulated into lipid nanoparticles (LNPs) by combining them with helper lipids, cholesterol, PEG-lipids, and firefly luciferase (Fluc) mRNA. The formulation follows an optimized molar ratio to ensure efficient encapsulation and LNP stability [23].

To evaluate mRNA delivery efficiency, LNPs are dosed into two replicate 96-well plates containing HBE cells. This redundancy helps mitigate technical variations that may arise during cell plating, incubation, or liquid handling. The second liquid handler ensures precise and uniform LNP dosing, minimizing well-to-well variability. The plates are then incubated for 18 hours, allowing sufficient time for cellular uptake and mRNA translation. After incubation, a robotic arm retrieves the treated plates and transfers them back to the liquid handler, where a luciferase substrate reagent is automatically added to initiate the luminescence reaction.

To further enhance data reliability, each 96-well plate is read twice in the bioluminescence plate reader, resulting in *four replicate readings* per lipid (two independent cell culture plates × two reads per plate). This approach reduces technical fluctuations, ensuring that potential inconsistencies in cell incubation or signal detection are minimized and yielding highly reproducible transfection potency (mTP) measurements. The luminescence intensity, which directly quantifies mRNA transfection efficiency, is automatically recorded and processed for analysis.

All experimental data undergo real-time processing within LUMI-lab’s integrated software modules, which perform error detection, data normalization, and quality control. The system automatically corrects inconsistencies, removes outliers, and normalizes luminescence readings using control wells. The processed data are then fed back into LUMI-model, iteratively refining its predictive framework with each experimental cycle. By continuously learning from experimental results, LUMI-model improves its ability to propose increasingly effective lipid candidates in subsequent iterations. The custom hardware and software designs are detailed in Supplementary Notes S.3 and S.4.

To balance optimization and exploration, each iteration follows a two-pronged experiment strategy. The first experiment plate consists of 96 lipid candidates prioritized for high predicted mTP, ensuring that the most promising candidates are experimentally validated. The second plate includes 96 lipids selected based on high model uncertainty, capturing molecules with the greatest variability across ensemble model predictions. This selection approach allows the system to refine its understanding of lipid structure-function relationships while identifying new high-performing lipid architectures.

#### 4.3.1 Online quality control and data processing

To ensure the reliability and consistency of luminescence readings in high-throughput screening, we implemented an automated online quality control (QC) and data processing pipeline. This pipeline integrates log transformation, normalization, and quality assessment of experimental data obtained from 96-well plate assays, enabling real-time validation for downstream analysis.

Raw luminescence readings from each well were first log-transformed using base-2 logarithm to stabilize variance and improve comparability across experiments. The first two wells (A1, B1), which contained no ionizable lipids, were designated as empty control wells, while subsequent wells without lipid components but containing benchmark lipids (e.g., MC3) served as positive control wells. The remaining wells represented experimental lipid candidates. Normalization was performed by adjusting each reading relative to the control wells:

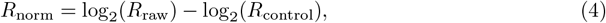

where *R*_raw_ represents the raw luminescence reading, and *R*_control_ is the mean control well reading. Any negative normalized value was clipped to zero to ensure robust normalization while maintaining interpretability.

Each well in the 96-well plate typically contained up to four replicate readings (Setion 4.3), with the first two replicates derived from one well and the latter two from another. The QC pipeline assessed the consistency of replicate readings using predefined thresholds. Replicates were deemed reliable if the absolute difference between paired readings did not exceed a predefined threshold of 3 in log scale. If the difference exceeded this threshold but the absolute values were high (*R*_norm_ *>* 7), the maximum value was retained; otherwise, the data point was flagged as unreliable. For multiple replicates deemed trustworthy, the final well reading was computed as the max reading of the replicates.

This automated QC and data processing pipeline enables real-time validation of experimental data, ensuring that only high-confidence readings are used in subsequent analyses. The framework is designed to scale efficiently across experimental iterations, providing a robust foundation for adaptive learning and active optimization within LUMI-lab.

### 4.4 Pre-experimental benchmark dataset

To validate our model design, we utilized a dataset generated using the established high-throughput synthesis and screening method [48], which included 1,920 unique ionizable lipids synthesized through the 4CR-Ugi reaction [49]. This combinatorial lipid library was constructed with 20 diverse amines, 4 alkyl aldehydes, 4 alkyl carboxylic acids, and 6 isocyanides. The 1,920 LNPs were tested *in vitro*, providing labeled transfection data for the pre-experimental test.

### 4.5 Pre-experimental evaluation and comparison

To assess the predictive performance of LUMI-model, we conducted a systematic benchmark analysis against several baseline models. The evaluation aimed to determine the effectiveness of pretraining strategies and 3D molecular representations in improving mTP predictions. We compared LUMI-model against the following alternatives: a variant of LUMI-model trained without any pretraining, a version that excludes continual pretraining while retaining the initial unsupervised pretraining step, a graph-based neural network without 3D inputs following MolCLR [24], and a multi-layer perceptron (MLP) model that utilizes predefined molecular descriptors from Mordred [26] instead of learned representations.

The models were evaluated on a dataset of 1,920 lipid molecules with experimentally validated transfection efficacies (Section 4.4). To ensure robust performance estimation, we employed a five-fold rolling validation scheme, where each model was trained five times, with 20% of the data held out as a test set in each iteration. The final reported performance for each model was obtained by averaging results across the five test sets.

Model accuracy was assessed using Spearman and Pearson correlation coefficients, which measure the rank-based and linear associations between predicted and experimental mTP values, respectively. These metrics were computed on two evaluation subsets: the full test set containing all lipids and the subset comprising the top 25% of lipids based on mTP, which specifically evaluates model performance on high-performing lipid candidates. This comprehensive benchmarking allowed us to quantify the contribution of pretraining, continual pretraining, and 3D molecular representations in enhancing model accuracy.

### 4.6 Wetlab materials

All materials were prepared and processed under nuclease-free conditions throughout synthesis and formulation. CleanCap Firefly Luciferase mRNA and CleanCap M6 CRISPR-Associated Protein 9 mRNA (N1-methylpseudouridine), sourced from TriLink BioTechnologies, were used without modification. These mRNAs were stored at -80°C and thawed on ice before use. MC3 and SM-102 were obtained from Echelon Biosciences, while amine headgroups and other precursors for ionizable lipid synthesis were purchased from Sigma-Aldrich and TCI America. Biodegradable lipid tails were synthesized and purified via flash column chromatography, and their structures were confirmed using ^1^H NMR (400 MHz, CDCl_3_) at the nuclear magnetic resonance facility in the Department of Chemistry, University of Toronto (Supplementary Figures S12-S37). High-resolution mass spectra of the synthesized materials were acquired using an LC-MS spectrometer at the Centre for Pharmaceutical Oncology (CPO), University of Toronto. Cell Counting Kit 8 (CCK-8, ab228554) was purchased from Abcam. Single-guide RNA (sgRNA) was chemically modified with 2’-O-methylation at the 2’-hydroxyl group and phosphorothioate bonds at the non-bridging oxygen in the phosphate backbone, specifically at or between the first three and last three nucleotides. This sgRNA was purchased from IDT [50].

### 4.7 *In vitro* high-throughput screening of ionizable lipids

To create the high-throughput screening library, we designed a modular synthesis approach that enables systematic variation of functional groups. This strategy requires key building blocks, including isocyanides, amines, carboxylic acids, and aldehydes, which serve as fundamental reactants in the formation of intermediates leading to the final products. Briefly, 96 distinct stock solutions of amines, carboxylic acids, aldehydes, and isocyanides were prepared in a 96-deep-well plate and dissolved in ethanol. Using an OT-2 liquid handler, these solutions were transferred into a 96-well PCR plate. For the synthesis of the final products, all isocyanides and amines were commercially sourced. A portion of the tails of aldehyde and carboxylic acid were synthesized as detailed in Supplementary Note S.5. Ionizable lipids were synthesized directly within each well by mixing the components in a 1:1:1:1 ratio and stirring the reactions for 18 hours. LNP formulations and high-throughput screening methods were conducted as previously described [48, 30]. For *in vitro* screening, all LNPs were formulated using a molar ratio of 35:16:46.5:2.5 for ionizable lipid, Dioleoylphosphatidylethanolamine (DOPE), cholesterol, and dimyristoyl-sn-glycero-3-phosphoethanolamine–N-(methoxy(polyethylene glycol)-2000) (C14-PEG2000), following the protocols established in our previous work [48].

### 4.8 Manual lipid synthesis and LNP characterization for post-optimization validation

The lipids of LUMI-1 to LUMI-6 and LUMI-6D were synthesized manually, and the detailed synthesis and purification methods were described in Supplementary Note S.6. All the LNPs used for I.T. injection experiments were formulated with ionizable lipid, 1,2-dioleoyl-3-trimethylammonium-propane (DOTAP), cholesterol, and C14-PEG2000 in a molar ratio of 35:28:34.5:2. For the other LNP formulations, including SM-102, and MC3, were prepared with a molar ratio of 50:10:38.5:1.5 for ionizable lipid, distearoylphosphatidylcholine (DSPC), cholesterol, and C14-PEG2000, respectively. The LNPs were synthesized using a microfluidic chip device by combining an aqueous phase containing mRNA with an ethanol phase containing the lipids. The aqueous phase was prepared using a 10 mM citrate buffer with the desired concentration of mRNA. The ethanol phase was prepared by solubilizing a lipid mixture, which included an ionizable lipid, helper lipid, cholesterol, and C14-PEG2000. The lipids were mixed with the appropriate molar ratios as determined in the LNP formulation parameters. Following synthesis, the LNPs were subjected to dialysis to remove ethanol and exchange the buffer. Dialysis was performed against phosphate-buffered saline (PBS) using either a 10,000 molecular weight cut-off (MWCO) Pierce 96-well microdialysis plate (ThermoFisher) or a 20,000 MWCO dialysis cassette (ThermoFisher). This step ensured buffer exchange while maintaining the structural integrity and stability of the LNPs. The size and polydispersity index (PDI) of LNPs were measured using Zetasizer Nano ZS (Malvern Instruments) for quality control.

### 4.9 Cell and Animal Experiments

The HBE cell line was obtained from Sigma-Aldrich and maintained in Eagle’s Minimum Essential Medium (MEM) supplemented with 10% fetal bovine serum (Gibco) and 1% Penicillin/Streptomycin (Gibco). All animal experiments were approved by the University Health Network Animal Resources Centre and conducted in compliance with Animal Use Protocol (AUP) guidelines. C57BL/6 and B6.Cg-Gt(ROSA)26Sort^tm9(CAG-tdTomato)Hze^/J (Ai9) mice were purchased from Jackson Laboratory. For the lung luminescence assay, 50,*µ*L of LNP (0.25,mg/kg Fluc mRNA) was administered via I.T. injection. Six hours post-dosing, C57BL/6 mice received an intraperitoneal (I.P.) injection of 2,mg D-luciferin (0.1,g/kg, 10,mg/mL) to facilitate luminescence imaging. Anesthesia was provided using 1.5% isoflurane in oxygen, and the mice were euthanized 10 minutes later. The lungs were excised and imaged using the Xenogen *In Vivo* Imaging System (IVIS, PerkinElmer). The total flux (photons per second) of bioluminescence in each organ was quantified. Imaging data were analyzed and quantified using Living Image Software (PerkinElmer). For CRISPR-Cas9 gene editing in Ai9 mice, 50 *µ*L of LNP-Cas9/sgRNA (containing Cas9 mRNA and sgRNA at a weight ratio of 4:1, with a Cas9 mRNA dose of 1 mg/kg) was administered via I.T. injection according to the specified dosing schedule. 9 days after the initial dose, the mice were euthanized, and their lungs were collected for flow cytometry analysis and immunofluorescence staining. Briefly, the tissues were washed twice with 1 × PBS, followed by overnight fixation in 0.5% paraformaldehyde prepared in 1 × PBS. The cells were then resuspended in 1 × PBS containing 5% FBS and analyzed using a BD Biosciences flow cytometer. The antibodies and dyes were listed in Supplementary Note S.7. To evaluate the impact of brominated and non-brominated ionizable lipids on mTP, LUMI-6 and its debrominated derivative, LUMI-6D, were synthesized. Both ionizable lipids were formulated with Fluc mRNA using the same method and molar ratio (35:16:46.5:2.5 for ionizable lipid, DOPE, cholesterol, and C14-PEG2000). The resulting LNPs were incubated with HBE cells for 18 hours. mTP was measured using a bioluminescence assay on a plate reader. The cytotoxicity of lipid candidates was assessed using the CCK-8 assay following the manufacturer’s instructions. HBE cells were seeded into a 96-well plate at a density of 5 × 10^4^ cells per well. The mRNA-LNPs containing the test lipids were added into the cells and followed by an 18-hour cell incubation. After the incubation, 10 *µ*L of CCK-8 reagent was added to each well containing 100 *µ*L of culture medium. The plate was then incubated at 37^°^C for 2 hours in a humidified cell incubator. The absorbance was measured at 450 nm using a plate reader, with the color intensity corresponding to the number of viable cells.

### 4.10 Statistical analysis

Statistical comparisons between two groups were conducted using a two-tailed Student’s t-test, while the one-way analysis of variance (ANOVA) was used for comparisons involving more than two groups. Data analysis was performed using GraphPad Prism 10.0. Statistical significance was defined as *P <* 0.05, with significance levels indicated as *\*P <* 0.05, *\*\*P <* 0.01, *\*\*\*P <* 0.001, and *\*\*\*\*P <* 0.0001.

## Data Availability

The nuclear magnetic resonance (NMR) data supporting this study are provided in the Supplementary Information. All other experimental datasets, including raw and processed transfection efficiency measurements, will be made publicly available upon publication.

## Code Availability

The source code for all LUMI-lab software modules and the design files for custom hardware components are available at https://github.com/bowenli-lab/LUMI-lab. The LUMI-model checkpoints will be released at the same repository upon publication.

## Acknowledgment

This research was funded by the Accelerate Translation grant from the Acceleration Consortium (518240), the GSK Chair Professorship, the startup fund from the Leslie Dan Faculty of Pharmacy, the operating fund from the Princess Margaret Cancer Centre, the Connaught Fund (514681), the J.P. Bickell Foundation (515159), the Canada Research Chairs Program (CRC-2022-00575), the Canadian Institutes of Health Research (PJH-185722), the New Frontiers in Research Fund (NFRFE-2023-00203), the Natural Sciences and Engineering Research Council of Canada (RGPIN-2023-05124), and the Canada Foundation for Innovation – John R. Evans Leaders Fund (43711). This research was made possible in part through computing resources provided by Calcul Québec (calculquebec.ca) and the Digital Research Alliance of Canada (alliancecan.ca). We also acknowledge Modal AI (modal.com) for providing additional computing resources to support this work. K.P. made the contributions to this work while he was an undergraduate student at University of Toronto. He is currently affiliated with Stanford University.

## Author Contributions

H.C., Y.X., and B.L. conceptualized the study and designed the experiments. Y.X. led the design and development of the chemical library and wet-lab experiments. H.C. led the design and development of AI and automated systems. K.P. had core contributions to software development, model development, and training. G.L. had core contributions to the custom hardware design and creation. F.G. had core contributions to the *in vivo* experiments. Y.X., H.C., and B.L. wrote the initial draft of the paper. All authors discussed the results and edited the paper. B.W. and B.L. acquired funding and supervised the project.

## References

[1] Milad Abolhasani and Eugenia Kumacheva. “The rise of self-driving labs in chemical and materials sciences”. In: Nature Synthesis 2.6 (2023), pp. 483–492.

[2] Tanja Dimitrov et al. “Autonomous molecular design: then and now”. In: ACS applied materials & interfaces 11.28 (2019), pp. 24825–24836.

[3] Jiayu Peng et al. “Human-and machine-centred designs of molecules and materials for sustainability and decarbonization”. In: Nature Reviews Materials 7.12 (2022), pp. 991–1009.

[4] Eric Stach et al. “Autonomous experimentation systems for materials development: A community perspective”. In: Matter 4.9 (2021), pp. 2702–2726.

[5] Benjamin P MacLeod et al. “Self-driving laboratory for accelerated discovery of thin-film materials”. In: Science Advances 6.20 (2020), eaaz8867.

[6] Brent A Koscher et al. “Autonomous, multiproperty-driven molecular discovery: From predictions to measurements and back”. In: Science 382.6677 (2023), eadi1407.

[7] Benjamin Burger et al. “A mobile robotic chemist”. In: Nature 583.7815 (2020), pp. 237–241.

[8] OpenAI. GPT-4 Technical Report. 2023. arXiv: 2303.08774 [cs.CL].

[9] Kizzmekia S Corbett et al. “SARS-CoV-2 mRNA vaccine design enabled by prototype pathogen preparedness”. In: Nature 586.7830 (2020), pp. 567–571.

[10] Annette B Vogel et al. “BNT162b vaccines protect rhesus macaques from SARS-CoV-2”. In: Nature 592.7853 (2021), pp. 283–289.

[11] Fernando P Polack et al. “Safety and efficacy of the BNT162b2 mRNA Covid-19 vaccine”. In: New England journal of medicine 383.27 (2020), pp. 2603–2615.

[12] Lindsey R Baden et al. “Efficacy and safety of the mRNA-1273 SARS-CoV-2 vaccine”. In: New England journal of medicine 384.5 (2021), pp. 403–416.

[13] PR Cullis and PL Felgner. “The 60-year evolution of lipid nanoparticles for nucleic acid delivery”. In: Nature Reviews Drug Discovery 23.9 (2024), pp. 709–722.

[14] Xuexiang Han et al. “An ionizable lipid toolbox for RNA delivery”. In: Nature communications 12.1 (2021), p. 7233.

[15] Jacob Witten et al. “Artificial intelligence-guided design of lipid nanoparticles for pulmonary gene therapy”. In: Nature biotechnology (2024), pp. 1–10.

[16] Riccardo Rampado and Dan Peer. “Design of experiments in the optimization of nanoparticle-based drug delivery systems”. In: Journal of Controlled Release 358 (2023), pp. 398–419.

[17] Rishi Bommasani et al. “On the opportunities and risks of foundation models”. In: arXiv preprint arXiv:2108.07258 (2021).

[18] Bowen Li et al. “Combinatorial design of nanoparticles for pulmonary mRNA delivery and genome editing”. In: Nature biotechnology 41.10 (2023), pp. 1410–1415.

[19] Shuying Chen et al. “Nanotechnology-based mRNA vaccines”. In: Nature Reviews Methods Primers 3.1 (2023), p. 63.

[20] Lulu Xue et al. “Responsive biomaterials: optimizing control of cancer immunotherapy”. In: Nature Reviews Materials 9.2 (2024), pp. 100–118.

[21] Xucheng Hou et al. “Lipid nanoparticles for mRNA delivery”. In: Nature Reviews Materials 6.12 (2021), pp. 1078–1094.

[22] Yue Xu et al. “Rational design and combinatorial chemistry of ionizable lipids for RNA delivery”. In: Journal of Materials Chemistry B 11.28 (2023), pp. 6527–6539.

[23] Kevin J Kauffman et al. “Optimization of lipid nanoparticle formulations for mRNA delivery in vivo with fractional factorial and definitive screening designs”. In: Nano letters 15.11 (2015), pp. 7300–7306.

[24] Yuyang Wang et al. “Molecular contrastive learning of representations via graph neural networks”. In: Nature Machine Intelligence 4.3 (2022), pp. 279–287.

[25] Ashish Vaswani et al. “Attention is all you need”. In: Advances in neural information processing systems 30 (2017).

[26] Hirotomo Moriwaki et al. “Mordred: a molecular descriptor calculator”. In: Journal of cheminformatics 10 (2018), pp. 1–14.

[27] Zubao Gan et al. “Nanoparticles containing constrained phospholipids deliver mRNA to liver immune cells in vivo without targeting ligands”. In: Bioengineering & translational medicine 5.3 (2020), e10161.

[28] Siddharth Patel et al. “Naturally-occurring cholesterol analogues in lipid nanoparticles induce polymorphic shape and enhance intracellular delivery of mRNA”. In: Nature communications 11.1 (2020), p. 983.

[29] Lining Zheng et al. “Lipid nanoparticle topology regulates endosomal escape and delivery of RNA to the cytoplasm”. In: Proceedings of the National Academy of Sciences 120.27 (2023), e2301067120.

[30] Bowen Li et al. “Accelerating ionizable lipid discovery for mRNA delivery using machine learning and combinatorial chemistry”. In: Nature Materials (2024), pp. 1–7.

[31] Brandon R Smith, Candice M Eastman, and Jon T Njardarson. “Beyond C, H, O, and N! Analysis of the elemental composition of US FDA approved drug architectures: Miniperspective”. In: Journal of Medicinal Chemistry 57.23 (2014), pp. 9764–9773.

[32] Hyung Cho, Kenneth J Wolf, and Eric J Wolf. “Management of ocular inflammation and pain following cataract surgery: focus on bromfenac ophthalmic solution”. In: Clinical Ophthalmology (2009), pp. 199–210.

[33] Eduard Potapskyi et al. “Introducing bromine in the molecular structure as a good strategy to the drug design”. In: Journal of Medical Science 93.3 (2024), e1128–e1128.

[34] Zhijian Xu et al. “Halogen bond: its role beyond drug–target binding affinity for drug discovery and development”. In: Journal of chemical information and modeling 54.1 (2014), pp. 69–78.

[35] Gengmo Zhou et al. “Uni-mol: A universal 3d molecular representation learning framework”. In: The Eleventh International Conference on Learning Representations. 2023.

[36] Chengxuan Ying et al. “Do transformers really perform badly for graph representation?” In: Advances in neural information processing systems 34 (2021), pp. 28877–28888.

[37] Muhammed Shuaibi et al. “Rotation invariant graph neural networks using spin convolutions”. In: arXiv preprint arXiv:2106.09575 (2021).

[38] Josiah Willard Gibbs. Elementary principles in statistical mechanics: developed with especial reference to the rational foundations of thermodynamics. C. Scribner”s sons, 1902.

[39] Jimmy Lei Ba. “Layer normalization”. In: arXiv preprint arXiv:1607.06450 (2016).

[40] Ting Chen et al. “A simple framework for contrastive learning of visual representations”. In: International conference on machine learning. PMLR. 2020, pp. 1597–1607.

[41] James Kirkpatrick et al. “Overcoming catastrophic forgetting in neural networks”. In: Proceedings of the national academy of sciences 114.13 (2017), pp. 3521–3526.

[42] Gui Citovsky et al. “Batch active learning at scale”. In: Advances in Neural Information Processing Systems 34 (2021), pp. 11933–11944.

[43] Sereina Riniker and Gregory A Landrum. “Better informed distance geometry: using what we know to improve conformation generation”. In: Journal of chemical information and modeling 55.12 (2015), pp. 2562–2574.

[44] Shuzhe Wang et al. “Improving conformer generation for small rings and macrocycles based on distance geometry and experimental torsional-angle preferences”. In: Journal of chemical information and modeling 60.4 (2020), pp. 2044–2058.

[45] Greg Landrum. RDKit: Open-source cheminformatics. 2006. url: https://www.rdkit.org.

[46] Paolo Tosco, Nikolaus Stiefl, and Gregory Landrum. “Bringing the MMFF force field to the RDKit: implementation and validation”. In: Journal of cheminformatics 6 (2014), pp. 1–4.

[47] Thomas A Halgren. “Merck molecular force field. I. Basis, form, scope, parameterization, and performance of MMFF94”. In: Journal of computational chemistry 17.5-6 (1996), pp. 490–519.

[48] Yue Xu et al. “AGILE platform: a deep learning powered approach to accelerate LNP development for mRNA delivery”. In: Nature Communications 15.1 (2024), p. 6305.

[49] Zepeng He et al. “A Multidimensional Approach to Modulating Ionizable Lipids for High-Performing and Organ-Selective mRNA Delivery”. In: Angewandte Chemie International Edition 62.43 (2023), e202310401.

[50] Tuo Wei et al. “Lung SORT LNPs enable precise homology-directed repair mediated CRISPR/Cas genome correction in cystic fibrosis models”. In: Nature communications 14.1 (2023), p. 7322.

