## Supplementary Notes and Figures for "LUMI-lab: a Foundation Model-Driven Autonomous Platform Enabling Discovery of New Ionizable Lipid Designs for mRNA Delivery"

**Supplementary Figures** **17**

#### S.1 Training details for LUMI-model

Supplementary Table S1: Unsupervised Pretraining Parameters

| **Parameter** | **Value** |
|---|---|
| Batch size | 448 |
| Scheduler | Polynomial decay |
| Learning rate | $1 \times 10^{-4}$ |
| Optimizer | Adam |
| Optimizer betas | 0.9, 0.99 |
| Epsilon | $1 \times 10^{-6}$ |
| Weight decay | $1 \times 10^{-4}$ |
| Dropout | 0.0 |
| Warmup | 1000 steps |
| Training steps | 200000 steps |
| Model width | 512 |
| Number of attention heads | 64 |
| Atom mask probability | 0.15 |
| Atom coordinate noise type | uniform |
| Atom coordinate noise scale | 1.0 |
| Masked atom loss scale | 0.5 |
| Coordinate recovery loss scale | 5.0 |
| Contrastive loss scale | 10.0 |

Supplementary Table S2: Continual Pretraining Parameters

| **Parameter** | **Value** |
|---|---|
| Batch size | 448 |
| Scheduler | Polynomial decay |
| Learning rate | $2 \times 10^{-5}$ |
| Optimizer | Adam |
| Optimizer betas | 0.9, 0.99 |
| Epsilon | $1 \times 10^{-6}$ |
| Weight decay | $1 \times 10^{-4}$ |
| Dropout | 0.0 |
| Warmup | 1000 steps |
| Training steps | 40000 steps |
| Model width | 512 |
| Number of attention heads | 64 |
| Atom mask probability | 0.15 |
| Atom coordinate noise type | uniform |
| Atom coordinate noise scale | 1.0 |
| Masked atom loss scale | 0.5 |
| Coordinate recovery loss scale | 5.0 |
| Contrastive loss scale | 10.0 |
| lipid:others sampling ratio | 1:0.5 |

#### S.2 Chemical library for expanded lipid dataset

| ID | SMILES |
|---|---|
| **Category A: Amines** | |
| A1 | NCCN(CC1)CCN1CCN |
| A2 | CCN(CC)CCCC(C)N |
| A3 | NCC1CCN(C)CC1 |
| A4 | NC1C[N@@]2CC[C@H]1CC2 |
| A5 | NCCCN(CCCN)CCCN |
| A6 | NCC1CCN(CCCO)CC1 |
| A7 | NCC1CCN(CCCCO)CC1 |
| A8 | NCC1CCN(CCCCCO)CC1 |
| A9 | NC1CCN(CCCO)CC1 |
| A10 | NC1CCN(CCCCO)CC1 |
| A11 | NC1CCN(CCCCCO)CC1 |
| A12 | NCCC1CCN(CCO)CC1 |
| A13 | NCCC1CCN(CCCO)CC1 |
| A14 | NCCC1CCN(CCCCO)CC1 |
| A15 | NCCN(CC)CC |
| A16 | NC(CC1)CCC1N(CC)CC |
| A17 | NC(CC1)CCC1N2CCCC2 |
| A18 | NC(CC1)CCC1N2CCCCC2 |
| A19 | NC(CC1)CCC1N2CCCCCC2 |
| A20 | NCCCCN1CCCC1 |
| A21 | NCC1CN(C)CC1 |
| A22 | NCCCCN(C)C |
| A23 | NCCCCCN(C)C |
| A24 | NCCN1CCN(C)CC1 |
| A25 | NCN(CC1)CCN1CN |
| A26 | NCCCN(CC1)CCN1CCCN |
| A27 | O=C(C(CCCCN)N1)NC(CCCCN)C1=O |
| A28 | O=C(C(CCCN)N1)NC(CCCN)C1=O |
| A29 | O=C([C@H](CCN)N1)N[C@H](CCN)C1=O |
| A30 | O=C(C(CN)N1)NC(CN)C1=O |
| A31 | O=C(C(N)N1)NC(N)C1=O |
| A32 | OC(O)N1CCC(CCN)CC1 |
| A33 | NCCN(C)C |
| A34 | NN1CCCCC1 |
| A35 | NCCCN1CCOCC1 |
| A36 | NCCN1CCNCC1 |
| A37 | NCCCN(C)C |
| A38 | NCC(CCC1)N1CC |
| A39 | NC1CCN(CC1)C |
| A40 | NCCN(CCN)CCN |
| A41 | NN1CCOCC1 |
| A42 | NCCN(CC)CCC |
| A43 | NCCCN(C)CCCN |
| A44 | NCCN1CCCC1 |
| A45 | NCCN(C(C)C)C(C)C |
| A46 | NC1=CC(C)=NN1C |

| ID | SMILES |
|---|---|
| A47 | NCCCN(CC)CC |
| A48 | CN(N)C |
| A49 | NCCCN1CCCC1 |
| A50 | NCCN1CCCCC1 |
| A51 | NCCCN(CCCC)CCCC |
| A52 | NC(CC1)CCC1N(C)C |
| A53 | NCCCN1C(C)CCCC1 |
| A54 | NC1=CNN=C1 |
| A55 | NC1CCN(C(C)C)CC1 |
| A56 | NC1=NC2=CC=CC=C2N1 |
| A57 | NCC1CCN(CCO)CC1 |
| A58 | NCC1CCN(O)CC1 |
| A59 | OCC1CN(CCCN)CCC1 |
| A60 | OCN1CCC(CCN)CC1 |
| A61 | OCC1CCN(CC2=CC(N)=CC=C2)CC1 |
| A62 | CN1C=CN=C1CN |
| A63 | NC1=NC=CS1 |
| A64 | NCCCN1CCCCCC1 |
| **Category B: Isocyanides** | |
| B1 | CCCCCC(C)[N+]#[C-] |
| B2 | CCCCCCC(C)[N+]#[C-] |
| B3 | CCC(C)[N+]#[C-] |
| B4 | CC(C)C[N+]#[C-] |
| B5 | CCC(C)C[N+]#[C-] |
| B6 | CCCC(C)C[N+]#[C-] |
| B7 | CCCCC(C)C[N+]#[C-] |
| B8 | CCCCCC(C)C[N+]#[C-] |
| B9 | CCC(CC)[N+]#[C-] |
| B10 | CCC(CC)C[N+]#[C-] |
| B11 | CC(C)CC[N+]#[C-] |
| B12 | CCC(C)CC[N+]#[C-] |
| B13 | CCCC(C)CC[N+]#[C-] |
| B14 | CCCCC(C)CC[N+]#[C-] |
| B15 | CCCC(CCC)[N+]#[C-] |
| B16 | CCCC(CCC)C[N+]#[C-] |
| B17 | CCCC(CC[N+]#[C-])CCC |
| B18 | CCC(CC)CC[N+]#[C-] |
| B19 | CCCC(CC)CC[N+]#[C-] |
| B20 | CCCCC(CC)CC[N+]#[C-] |
| B21 | CC1CCC([N+]#[C-])CC1 |
| B22 | CCC1CCC([N+]#[C-])CC1 |
| B23 | CC(C)C1CCC([N+]#[C-])CC1 |
| B24 | CC(CC)C1CCC([N+]#[C-])CC1 |
| B25 | CCC(CC)C1CCC([N+]#[C-])CC1 |
| B26 | [C-]#[N+]C1CCC(C2CCCC2)CC1 |
| B27 | CN1CCC([N+]#[C-])CC1 |
| B28 | CCN1CCC([N+]#[C-])CC1 |
| B29 | CC(C)N1CCC([N+]#[C-])CC1 |
| B30 | CC(CC)N1CCC([N+]#[C-])CC1 |
| B31 | CCC(CC)N1CCC([N+]#[C-])CC1 |

| ID | SMILES |
|---|---|
| B32 | [C-]#[N+]C1CCN(C2CCCC2)CC1 |
| B33 | CCCN(CCC)[N+]#[C-] |
| B34 | CCCN(CCC)C[N+]#[C-] |
| B35 | CCN(CC[N+]#[C-])CC |
| B36 | CCCN(CCC)CC[N+]#[C-] |
| B37 | CCCN(CC)CC[N+]#[C-] |
| B38 | CCCCN(CC)CC[N+]#[C-] |
| B39 | CC1CCC([N+]#[C-])C1 |
| B40 | CCC1CCC([N+]#[C-])C1 |
| B41 | CC(C)C1CCC([N+]#[C-])C1 |
| B42 | CC(CC)C1CCC([N+]#[C-])C1 |
| B43 | CCC(CC)C1CCC([N+]#[C-])C1 |
| B44 | [C-]#[N+]C(C1)CCC1C2CCCC2 |
| B45 | CN1CCC([N+]#[C-])C1 |
| B46 | CCN1CCC([N+]#[C-])C1 |
| B47 | CC(C)N1CCC([N+]#[C-])C1 |
| B48 | CC(CC)N1CCC([N+]#[C-])C1 |
| B49 | CCC(CC)N1CCC([N+]#[C-])C1 |
| B50 | [C-]#[N+]C(C1)CCN1C2CCCC2 |
| B51 | C[N+]#[C-] |
| B52 | O=S(C[N+]#[C-])(C1=CC=C(C)C=C1)=O |
| B53 | [C-]#[N+]CN1C(C=CC=C2)=C2N=N1 |
| B54 | COC(C[N+]#[C-])=O |
| B55 | O=C(C[N+]#[C-])OCC |
| B56 | [C-]#[N+]C1CCCC1 |
| B57 | [C-]#[N+]C1CCCCC1 |
| B58 | [C-]#[N+]CCN1CCOCC1 |
| B59 | CC(CC(C)([N+]#[C-])C)(C)C |
| B60 | CCCC[N+]#[C-] |
| B61 | CC(C)([N+]#[C-])C |
| B62 | [C-]#[N+]C1(C[C@@H]2C3)C[C@@H]3C[C@@H](C2) |
| B1 | CCCCCC(C)[N+]#[C-] |
| B2 | CCCCCCC(C)[N+]#[C-] |
| B3 | CCC(C)[N+]#[C-] |
| B4 | CC(C)C[N+]#[C-] |
| B5 | CCC(C)C[N+]#[C-] |
| B6 | CCCC(C)C[N+]#[C-] |
| B7 | CCCCC(C)C[N+]#[C-] |
| B8 | CCCCCC(C)C[N+]#[C-] |
| B9 | CCC(CC)[N+]#[C-] |
| B10 | CCC(CC)C[N+]#[C-] |
| B11 | CC(C)CC[N+]#[C-] |
| B12 | CCC(C)CC[N+]#[C-] |
| B13 | CCCC(C)CC[N+]#[C-] |
| B14 | CCCCC(C)CC[N+]#[C-] |
| B15 | CCCC(CCC)[N+]#[C-] |
| B16 | CCCC(CCC)C[N+]#[C-] |
| B17 | CCCC(CC[N+]#[C-])CCC |
| B18 | CCC(CC)CC[N+]#[C-] |
| B19 | CCCC(CC)CC[N+]#[C-] |

| ID | SMILES |
|---|---|
| B20 | CCCCC(CC)CC[N+]#[C-] |
| B21 | CC1CCC([N+]#[C-])CC1 |
| B22 | CCC1CCC([N+]#[C-])CC1 |
| B23 | CC(C)C1CCC([N+]#[C-])CC1 |
| B24 | CC(CC)C1CCC([N+]#[C-])CC1 |
| B25 | CCC(CC)C1CCC([N+]#[C-])CC1 |
| B26 | [C-]#[N+]C1CCC(C2CCCC2)CC1 |
| B27 | CN1CCC([N+]#[C-])CC1 |
| B28 | CCN1CCC([N+]#[C-])CC1 |
| B29 | CC(C)N1CCC([N+]#[C-])CC1 |
| B30 | CC(CC)N1CCC([N+]#[C-])CC1 |
| B31 | CCC(CC)N1CCC([N+]#[C-])CC1 |
| B32 | [C-]#[N+]C1CCN(C2CCCC2)CC1 |
| B33 | CCCN(CCC)[N+]#[C-] |
| B34 | CCCN(CCC)C[N+]#[C-] |
| B35 | CCN(CC[N+]#[C-])CC |
| B36 | CCCN(CCC)CC[N+]#[C-] |
| B37 | CCCN(CC)CC[N+]#[C-] |
| B38 | CCCCN(CC)CC[N+]#[C-] |
| B39 | CC1CCC([N+]#[C-])C1 |
| B40 | CCC1CCC([N+]#[C-])C1 |
| B41 | CC(C)C1CCC([N+]#[C-])C1 |
| B42 | CC(CC)C1CCC([N+]#[C-])C1 |
| B43 | CCC(CC)C1CCC([N+]#[C-])C1 |
| B44 | [C-]#[N+]C(C1)CCC1C2CCCC2 |
| B45 | CN1CCC([N+]#[C-])C1 |
| B46 | CCN1CCC([N+]#[C-])C1 |
| B47 | CC(C)N1CCC([N+]#[C-])C1 |
| B48 | CC(CC)N1CCC([N+]#[C-])C1 |
| B49 | CCC(CC)N1CCC([N+]#[C-])C1 |
| B50 | [C-]#[N+]C(C1)CCN1C2CCCC2 |
| B51 | C[N+]#[C-] |
| B52 | O=S(C[N+]#[C-])(C1=CC=C(C)C=C1)=O |
| B53 | [C-]#[N+]CN1C(C=CC=C2)=C2N=N1 |
| B54 | COC(C[N+]#[C-])=O |
| B55 | O=C(C[N+]#[C-])OCC |
| B56 | [C-]#[N+]C1CCCC1 |
| B57 | [C-]#[N+]C1CCCCC1 |
| B58 | [C-]#[N+]CCN1CCOCC1 |
| B59 | CC(CC(C)([N+]#[C-])C)(C)C |
| B60 | CCCC[N+]#[C-] |
| B61 | CC(C)([N+]#[C-])C |
| B62 | [C-]#[N+]C1(C[C@@H]2C3)C[C@@H]3C[C@@H](C2) |
| **Category C: Lipid Aldehydes** | |
| C1 | CCCCC(C)C=O |
| C2 | CCCCCC(C)C=O |
| C3 | CCCCCCC(C)C=O |
| C4 | CCCCCCCC(C)C=O |
| C5 | CCCCCCCCC(C)C=O |
| C6 | CCCCCCCCCCC(C)C=O |

| ID | SMILES |
|---|---|
| C7 | CCCCCCCCCC(C)C=O |
| C8 | C=CCCCCCCCC(C)C=O |
| C9 | O=C(C(CCCCCCCC)CCCCCC)OCCCCC(C)C=O |
| C10 | CCCCCCC(CCCC)C(OCCCCC(C)C=O)=O |
| C11 | CCCCCCCC(C)C(CCCCCC)C=O |
| C12 | CCCCCCCCCCCCCCC(C)C=O |
| C13 | CCCCCCCCCCCCCCCC(C)C=O |
| C14 | CCCCCCCCCCCCCCCCC(C)C=O |
| C15 | CCCCCCCC/C=C\CCCCCCC(C)C=O |
| C16 | CCCCC/C=C\C/C=C\CCCCCCC(C)C=O |
| C17 | CCCC(C)CC=O |
| C18 | CCCCC(C)CC=O |
| C19 | CCCCCC(C)CC=O |
| C20 | CCCCCCC(C)CC=O |
| C21 | CCCCCCCC(C)CC=O |
| C22 | CCCCCCCCCC(C)CC=O |
| C23 | CCCCCCCCC(C)CC=O |
| C24 | C=CCCCCCCC(C)CC=O |
| C25 | O=C(C(CCCCCCCC)CCCCCC)OCCCC(C)CC=O |
| C26 | CCCCCCC(CCCC)C(OCCCC(C)CC=O)=O |
| C27 | CCCCCCCCC(C(C)CCCCC)C=O |
| C28 | CCCCCCCCCCCCCCC(CC)C=O |
| C29 | CCCCCCCCCCCCCCCC(CC)C=O |
| C30 | CCCCCCCCCCCCCCCCC(CC)C=O |
| C31 | CCCCCCCC/C=C\CCCCCCC(CC)C=O |
| C32 | CCCCC/C=C\C/C=C\CCCCCCC(CC)C=O |
| C33 | CCCC(CC)CC=O |
| C34 | CCCCC(CC)CC=O |
| C35 | CCCCCC(CC)CC=O |
| C36 | CCCCCCC(CC)CC=O |
| C37 | CCCCCCCC(CC)CC=O |
| C38 | CCCCCCCCCC(CC)CC=O |
| C39 | CCCCCCCCC(CC)CC=O |
| C40 | C=CCCCCCCC(CC)CC=O |
| C41 | O=C(C(CCCCCCCC)CCCCCC)OCCCC(CC)CC=O |
| C42 | CCCCCCC(CCCC)C(OCCCC(CC)CC=O)=O |
| C43 | CCCCCCCCC(C(CC)CCCCC)C=O |
| C44 | CCCCCCCCCCCCCCC(CCC)C=O |
| C45 | CCCCCCCCCCCCCCCC(CCC)C=O |
| C46 | CCCCCCCCCCCCCCCCC(CCC)C=O |
| C47 | CCCCCCCC/C=C\CCCCCCC(CCC)C=O |
| C48 | CCCCC/C=C\C/C=C\CCCCCCC(CCC)C=O |
| C49 | CCCCCC=O |
| C50 | CCCCCCC=O |
| C51 | CCCCCCCC=O |
| C52 | CCCCCCCCC=O |
| C53 | CCCCCCCCCC=O |
| C54 | CCCCCCCCCCCC=O |
| C55 | CCCCCCCCCCC=O |
| C56 | C=CCCCCCCCCC=O |

| ID | SMILES |
|---|---|
| C57 | O=C(C(CCCCCCCC)CCCCCC)OCCCCCC=O |
| C58 | CCCCCCC(CCCC)C(OCCCCCC=O)=O |
| C59 | CCCCCCCCC(CCCCCC)C=O |
| C60 | CCCCCCCCCCCCCCCCC=O |
| C61 | CCCCCCCCCCCCCCCCCC=O |
| C62 | CCCCCCCCCCCCCCCCCCC=O |
| C63 | CCCCCCCC/C=C\CCCCCCCC=O |
| C64 | CCCCC/C=C\C/C=C\CCCCCCCC=O |
| **Category D: Lipid Carboxylic Acids** | |
| D1 | OC(CC(C1)(C2)C[C@@H]3C[C@H]2C[C@H]1C3)=O |
| D2 | OC(C[C@@](C1)(C2)C[C@]3(O)C[C@H]2C[C@H]1C3)=O |
| D3 | OC(CCN1CCCCC1)=O |
| D4 | CN(CC(O)=O)C |
| D5 | CN(CCC(O)=O)C |
| D6 | CN(CCCC(O)=O)C |
| D7 | OC(CN(CC)CC)=O |
| D8 | OC(CN(CCO)CCO)=O |
| D9 | O=C(O)CCCCBr |
| D10 | O=C(O)CCCCCBr |
| D11 | O=C(O)CCCCCCCBr |
| D12 | CCCCCCC(CCCC)C(O)=O |
| D13 | OC(C(CCCCCCCC)CCCCCC)=O |
| D14 | CCCCCCCC(O)=O |
| D15 | OC(CCCCCCCC)=O |
| D16 | CC(CCC(O)=O)CCCCC |
| D17 | OC(CCCCCCCCC)=O |
| D18 | OC(/C=C/CCCCCCC)=O |
| D19 | OC(CCCCCCCCCC)=O |
| D20 | OC(CCCCCCCCC#C)=O |
| D21 | OC(CCCCCCCCC=C)=O |
| D22 | OC(CCCCCCCCCCC)=O |
| D23 | CCCCCCCCCCCCCC(O)=O |
| D24 | CCCCCCCCCCCCCCCC(O)=O |
| D25 | CCCCCCCCCCCCCCCCCC(O)=O |
| D26 | CCCCCCCC/C=C\CCCCCCCC(O)=O |
| D27 | CCCCC/C=C\C/C=C\CCCCCCCC(O)=O |
| D28 | OC(CCCCC(OCC(CCCCCC)CCCCCCCC)=O)=O |
| D29 | OC(CCCCC(OC(C)CCCCCCCCC)=O)=O |
| D30 | CCC(OC(CCCCC(O)=O)=O)CCCCCCCC |
| D31 | OC(CCCCC(OC(CCCCCC)CCC)=O)=O |
| D32 | CCCCCCCCC(OC(CCCCC(O)=O)=O)CCCCCCCC |
| D33 | CCC(CCC(OC(CCCCC(O)=O)=O)CC(C)C)CCCC |
| D34 | OC(CCCCC(OCCCCCCCCCC)=O)=O |
| D35 | OC(CCCCC(OCCCCCCCCCCC)=O)=O |
| D36 | OC(CCCCC(OCCCCCCCCCC=C)=O)=O |
| D37 | CCCCCCC(CCCCC)C(O)=O |
| D38 | OC(C(CCCCCCCC)CCCCCCC)=O |
| D39 | CCCCCCC(C)C(O)=O |
| D40 | OC(C(C)CCCCCCC)=O |
| D41 | OC(CCC(CCCCC)CC)=O |

| ID | SMILES |
|---|---|
| D42 | OC(C(C)CCCCCCCC)=O |
| D43 | OC(/C=C/C(C)CCCCCC)=O |
| D44 | OC(C(C)CCCCCCCCC)=O |
| D45 | OC(C(C)CCCCCCCC#C)=O |
| D46 | OC(C(C)CCCCCCCC=C)=O |
| D47 | OC(C(C)CCCCCCCCCC)=O |
| D48 | CCCCCCCCCCCCC(C)C(O)=O |
| D49 | OC(CCCCC(OC(CC)CCCCCCCCC=C)=O)=O |
| D50 | CCCCCCCCCCCCCCC(C)C(O)=O |
| D51 | CCCCCCCCCCCCCCCCC(C)C(O)=O |
| D52 | CCCCCCCC/C=C\CCCCCCC(C)C(O)=O |
| D53 | CCCCC/C=C\C/C=C\CCCCCCC(C)C(O)=O |
| D54 | OC(CCCCC(OCC(CCCCCCC)CCCCCCCC)=O)=O |
| D55 | OC(CCCCC(OC(CC)CCCCCCCCC)=O)=O |
| D56 | CCCCCCCCC(OC(CCCCC(O)=O)=O)CCC |
| D57 | OC(CCCCC(OC(CCCCCC)CCCC)=O)=O |
| D58 | OC(CCCCC(OC(CCCCCCCCC)CCCCCCCCC)=O)=O |
| D59 | CC(C)CC(OC(CCCCC(O)=O)=O)CCC(CCCC)CCC |
| D60 | OC(CCCCC(OC(CCC)CCCCCCCCC)=O)=O |
| D61 | OC(CCCCC(OC(CC)CCCCCCCCCC)=O)=O |

#### S.3 Design of open-source mechanical modules for LUMI-lab

**Liquid Sampler** The sampler is designed with two main components: a storage area and a sample loading module. The storage area comprises 96 peristaltic pumps, each paired with a 5 mL syringe. Each pump is dedicated to dispensing a single type of chemical liquid. Through silicone tubes, the peristaltic pumps transfer the liquid from the syringes to the corresponding wells of the 96 well sampler cap. The precise alignment of the loading heads on the sampler cap with the wells ensures accurate liquid dispensing (Supplementary Figure S7).

To optimize space utilization and maintain modular expandability, the 96 pumps are arranged across two layers. Each syringe is labeled with a unique QR code for tracking raw material usage. During operation, the 96-well plate is positioned on a motorized base that moves into the loading zone via a screw-driven mechanism. Based on the loading volumes provided by our model, precise and individualized dispensing is executed.

To mitigate errors caused by evaporation of residual liquid within the tubing during periods of inactivity, the system pre-wets the tubes by pumping out liquid before each operation. Subsequently, the residual liquid is completely retracted, followed by the formal loading process. To address discrepancies arising from variations in tubing lengths, a linear correction model is applied to adjust loading times, ensuring high accuracy and efficiency.

The sampler's controller utilizes a modular stacking driver board system, enabling scalability. This design allows for the potential integration of additional peristaltic pumps, significantly expanding the range of chemical liquids that can be sampled, thus demonstrating considerable future potential.

**Feeder for pipette tip racks and well plates** The feeder is an automated system designed to replenish experimental consumables, such as tip racks, with high precision and efficiency. Its structural components are composed of aluminum extrusion frames and 3D-printed parts, ensuring a lightweight yet durable design (Supplementary Figure S8). The core mechanism relies on a stepper motor coupled with a screw-driven lifting platform. This synchronous lifting mechanism enables the smooth and accurate positioning of consumables during the replenishment process, minimizing the risk of mechanical errors.

To enhance usability, the feeder includes a manual control interface that allows operators to lower the platform for restocking experimental consumables manually. This design ensures operational flexibility and facilitates straightforward maintenance and replenishment tasks. Additionally, the system features a modular design, allowing for the seamless connection of multiple feeders via standardized connectors. This modularity not only supports scalability but also enhances adaptability to varying experimental setups, making it suitable for a wide range of laboratory environments.

The control method of the feeder is designed for consistency and interoperability. It employs the same control methodology as the clamper system, utilizing a Raspberry Pi and a motor driver board for precise actuation. This integration ensures compatibility with other automated laboratory equipment, streamlining system management and reducing the complexity of multi-device coordination.

#### S.4 Design of software modules for LUMI-lab

We developed an integrated software framework to orchestrate LUMI-lab's mechanical components and manage complex, high-throughput experimental workflows. The overall architecture for the software framework is illustrated in Supplementary Figure S1, which interconnects the hardware resources, local control panel, and cloud computing.

The system architecture addresses three fundamental challenges in automated experimentation: (1) parallel control of multiple hardware operations, (2) closed-loop integration between computational modeling (LUMI-model) and wet experiments (LUMI-lab), and (3) the human-computer interface for progress monitoring.

To achieve the parallel control of hardware, we design a task planner for scheduling the tasks based on the pre-defined experiment protocol and the availability of the labware and consumables. This accelerates the experiment by maximizing the utilization of the available resources as depicted in Supplementary Figure S2. Most of the deployed hardware is controlled through HTTP or SSH protocol, including the robot arm (via Universal Robot UR5e Dashboard Server), liquid handler (via Opentrons API v2), and plate reader (via Microsoft Windows OLE service with Fastapi interface). For other hardware, including liquid sampler, cell incubator, and plate feeders, we use two Raspberry Pi's for controlling the malicious motors and sensors, we use Fastapi for communicating with Raspberry Pi.

Integration of LUMI-lab and LUMI-model is achieved through a distributed architecture optimized for high-throughput data processing. Experimental readouts from LUMI-lab are managed through a MongoDB database, which serves as the central data repository for model refinement. The LUMI-model framework implements a hybrid computing strategy: model fine-tuning is performed locally with one A6000 Ada GPU, while inference tasks are distributed across cloud-based GPU clusters. This architecture is essential for processing our extensive virtual library, consisting of over 221K molecules with more than 2.4M conformers to predict per ensemble model. We implement cloud-based parallel inference using Modal's serverless API and Docker containers with up to ten A100 GPUs. Prediction results are aggregated locally for subsequent iterations.

We developed an intuitive web-based control interface that enables real-time monitoring and remote operation of LUMI-lab. We deploy a control panel using Streamlit, which presents key metrics for LUMI-lab, such as the current experiment plan, latest readouts, and consumable status. The platform supports remote intervention capabilities, such as labware reconfiguration for consumable replenishment. System events, including experimental milestones and potential failures, are communicated through automated Slack notifications, ensuring continuous experimental oversight while minimizing the need for constant human supervision.

#### S.5 Synthesis of lipid Tails

Aldehyde tails were synthesized through either a one-step process (Route A) or a two-step process (Route B)(Supplementary Figure S10):

**Route A:** Aldehyde tails were synthesized directly from alcohol tails using Dess-Martin Periodinane (DMP) oxidation. Alcohol (1 mmol) was dissolved in 30 ml of anhydrous dichloromethane (DCM), and DMP was added. The mixture was stirred under nitrogen at room temperature for 2 hours. After confirming reaction completion via thin-layer chromatography (TLC), 200 ml of 50% (w/v) sodium thiosulfate pentahydrate solution was added and stirred for an additional 15 minutes. The organic layers were combined, washed with brine, dried over anhydrous sodium sulfate ($Na_2SO_4$), and concentrated to yield a crude colorless oil. Purification was carried out using silica gel chromatography with a gradient of 0–50% ethyl acetate in hexane to obtain the aldehyde tails.

**Route B:** Aldehyde tails were synthesized in two steps. First, esterification was carried out by dissolving alcohol (10.0 mmol), carboxylic acid (1.0 mmol), dicyclohexylcarbodiimide (DCC, 1.1 mmol), and 4-dimethylaminopyridine (DMAP, 0.2 mmol) in 20 ml of anhydrous DCM in a round-bottom flask. The reaction was stirred at room temperature under nitrogen for 24 hours. The reaction mixture was filtered to remove dicyclohexylurea byproducts, and the filtrate was evaporated under vacuum. The residue was purified using a CombiFlash BUCHI C-815 chromatography system with gradient elution (0%–20% ethyl acetate in hexane) to isolate the ester product. The product was then oxidized to aldehyde using the same DMP oxidation method described in Route A.

**6-oxohexyl 2-butyloctanoate (C9):** Follow the synthesis method (Route B) as above. The residue was purified by silica gel chromatography to give 6-hydroxyhexyl 2-butyloctanoate as a colorless oil (yield 58%). $^1$H NMR (400 MHz, $CDCl_3$) $\delta$ 4.04 (t, $J = 6.7$ Hz, 2H), 3.61 (t, $J = 6.6$ Hz, 2H), 2.28 (tt, $J = 9.0, 5.3$ Hz, 1H), 1.64–1.51 (m, 6H), 1.39–1.21 (m, 18H), 0.87–0.84 (m, 6H). After the product was oxidized, 6-oxohexyl 2-butyloctanoate (C9) was obtained as a colorless oil (yield 95%). $^1$H NMR (400 MHz, $CDCl_3$) $\delta$ 4.06 (t, $J = 6.6$ Hz, 2H), 2.53–2.38 (m, 2H), 2.29 (tt, $J = 9.0, 5.4$ Hz, 1H), 1.74–1.50 (m, 6H), 1.48–1.35 (m, 4H), 1.33–1.17 (m, 13H), 0.86 (td, $J = 7.0, 2.2$ Hz, 6H).

**6-oxohexyl 2-hexyldecanoate (C10):** Follow the synthesis method (Route B) as above. The residue was purified by silica gel chromatography to give 6-hydroxyhexyl 2-hexyldecanoate as a colorless oil (yield 52%). $^1$H NMR (400 MHz, $CDCl_3$) $\delta$ 4.05 (t, $J = 6.6$ Hz, 2H), 3.62 (t, $J = 6.7$ Hz, 2H), 2.29 (tt, $J = 9.0, 5.3$ Hz, 1H), 1.64–1.52 (m, 6H), 1.41–1.18 (m, 25H), 0.86 (t, $J = 6.7$ Hz, 6H). After the product was oxidized, 6-oxohexyl 2-hexyldecanoate (C10) was obtained as a colorless oil (yield 98%). $^1$H NMR (400 MHz, $CDCl_3$) $\delta$ 9.76 (dt, $J = 3.9, 1.9$ Hz, 1H), 4.06 (td, $J = 6.6, 3.5$ Hz, 2H), 2.61–1.98 (m, 3H), 1.69–1.22 (m, 29H), 0.86 (dq, $J = 6.9, 3.4$ Hz, 6H).

**2-hexyldecanal (C11):** Follow the synthesis method (Route A) as above. The residue was purified by silica gel chromatography to give 2-hexyldecanal (C11) as a colorless oil (yield 92%). $^1$H NMR (400 MHz, $CDCl_3$) $\delta$ 9.55 (d, $J = 3.2$ Hz, 1H), 2.22 (dqd, $J = 11.2, 5.4, 3.2$ Hz, 1H), 1.46–1.40 (m, 2H), 1.28–1.25 (m, 21H), 0.88 (d, $J = 2.2$ Hz, 6H).

**Palmitaldehyde (C12):** Follow the synthesis method (Route A) as above. The residue was purified by silica gel chromatography to give palmitaldehyde (C12) as a white powder (yield 95%). $^{1}$H NMR (400 MHz, $CDCl_3$) $\delta$ 9.75 (t, $J = 1.9$ Hz, 1H), 2.41 (td, $J = 7.4, 1.9$ Hz, 2H), 1.69–1.59 (m, 2H), 1.25 (s, 23H), 0.89–0.85 (m, 3H).

**Heptadecanal (C13):** Follow the synthesis method (Route A) as above. The residue was purified by silica gel chromatography to give heptadecanal (C13) as a white powder (yield 96%). $^{1}$H NMR (400 MHz, $CDCl_3$) $\delta$ 9.76 (t, $J = 1.9$ Hz, 1H), 2.42 (td, $J = 7.4, 1.9$ Hz, 2H), 1.64–1.61 (m, 2H), 1.26 (d, $J = 3.8$ Hz, 25H), 0.86 (s, 3H).

**Stearaldehyde (C14):** Follow the synthesis method (Route A) as above. The residue was purified by silica gel chromatography to give stearaldehyde (C14) as a white powder (yield 95%). $^{1}$H NMR (400 MHz, $CDCl_3$) $\delta$ 9.75 (t, $J = 1.9$ Hz, 1H), 2.41 (td, $J = 7.4, 1.9$ Hz, 2H), 1.78–1.56 (m, 2H), 1.24 (s, 27H), 0.96–0.79 (m, 3H).

**Olealdehyde (C15):** Follow the synthesis method (Route A) as above. The residue was purified by silica gel chromatography to give olealdehyde (C15) as a colorless oil (yield 90%). $^{1}$H NMR (400 MHz, $CDCl_3$) $\delta$ 9.76 (t, $J = 1.9$ Hz, 1H), 5.49–5.12 (m, 2H), 2.41 (td, $J = 7.4, 1.9$ Hz, 2H), 2.00 (q, $J = 5.9$ Hz, 4H), 1.68–1.57 (m, 2H), 1.39–1.23 (m, 20H), 0.87 (t, $J = 6.8$ Hz, 3H).

**(9Z,12Z)-octadeca-9,12-dienal (C16):** Follow the synthesis method (Route A) as above. The residue was purified by silica gel chromatography to give (9Z,12Z)-octadeca-9,12-dienal (C16) as a colorless oil (yield 90%). $^{1}$H NMR (400 MHz, $CDCl_3$) $\delta$ 9.76 (t, $J = 1.9$ Hz, 1H), 5.54–5.13 (m, 4H), 2.87–2.68 (m, 2H), 2.42 (td, $J = 7.4, 1.9$ Hz, 2H), 2.05 (q, $J = 6.9$ Hz, 4H), 1.62 (dd, $J = 8.6, 5.9$ Hz, 2H), 1.36–1.28 (m, 14H), 0.88 (d, $J = 2.7$ Hz, 3H).

Carboxylic acid tails were synthesized through either a one-step process: esterification was carried out by dissolving alcohol (1.0 mmol), adipic acid (10.0 mmol), dicyclohexylcarbodiimide (DCC, 1.1 mmol), and 4-dimethylaminopyridine (DMAP, 0.2 mmol) in 20 ml of anhydrous DCM in a round-bottom flask. The reaction was stirred at room temperature under nitrogen for 24 hours. The reaction mixture was filtered to remove dicyclohexylurea byproducts, and the filtrate was evaporated under vacuum. The residue was purified using a CombiFlash BUCHI C-815 chromatography system with gradient elution (0%-30% ethyl acetate in hexane) to isolate the ester product (Supplementary Figure S11).

**6-((2-hexyldecyl)oxy)-6-oxohexanoic acid (D28):** Follow the synthesis method as above. The residue was purified by silica gel chromatography to give 6-((2-hexyldecyl)oxy)-6-oxohexanoic acid (D28) as a white crystal (yield 75%). $^{1}$H NMR (400 MHz, $CDCl_3$) $\delta$ 3.94 (dd, $J = 5.9, 2.8$ Hz, 2H), 2.48–2.17 (m, 4H), 1.72–1.59 (m, 5H), 1.24 (d, $J = 3.3$ Hz, 24H), 0.86 (t, $J = 6.7$ Hz, 6H).

**6-oxo-6-(undecan-2-yloxy)hexanoic acid (D29):** Follow the synthesis method as above. The residue was purified by silica gel chromatography to give 6-oxo-6-(undecan-2-yloxy)hexanoic acid (D29) as a white crystal (yield 72%). $^{1}$H NMR (400 MHz, $CDCl_3$) $\delta$ 5.03–4.71 (m, 1H), 2.46–2.22 (m, 4H), 1.72–1.53 (m, 5H), 1.47–1.06 (m, 18H), 0.99–0.68 (m, 3H).

**6-oxo-6-(undecan-3-yloxy)hexanoic acid (D30):** Follow the synthesis method as above. The residue was purified by silica gel chromatography to give 6-oxo-6-(undecan-3-yloxy)hexanoic acid (D30) as a white crystal (yield 69%). $^1$H NMR (400 MHz, $CDCl_3$) $\delta$ 4.80 (ddd, $J = 12.3, 6.8, 5.5$ Hz, 1H), 2.52–2.19 (m, 4H), 1.71–1.46 (m, 8H), 1.24 (s, 12H), 0.86 (td, $J = 7.1, 2.1$ Hz, 6H).

**6-(decan-4-yloxy)-6-oxohexanoic acid (D31):** Follow the synthesis method as above. The residue was purified by silica gel chromatography to give 6-(decan-4-yloxy)-6-oxohexanoic acid (D31) as a white crystal (yield 75%). $^1$H NMR (400 MHz, $CDCl_3$) $\delta$ 4.87 (tt, $J = 7.1, 5.4$ Hz, 1H), 2.47–2.18 (m, 4H), 1.72–1.60 (m, 4H), 1.56–1.41 (m, 4H), 1.34–1.15 (m, 10H), 0.87 (dt, $J = 9.8, 7.2$ Hz, 6H).

**6-(heptadecan-9-yloxy)-6-oxohexanoic acid (D32):** Follow the synthesis method as above. The residue was purified by silica gel chromatography to give 6-(heptadecan-9-yloxy)-6-oxohexanoic acid (D32) as a white crystal (yield 68%). $^1$H NMR (400 MHz, $CDCl_3$) $\delta$ 4.95–4.79 (m, 1H), 2.50–2.18 (m, 4H), 1.96–1.06 (m, 26H), 0.87 (dt, $J = 9.5, 7.2$ Hz, 6H).

**6-((7-ethyl-2-methylundecan-4-yl)oxy)-6-oxohexanoic acid (D33):** Follow the synthesis method as above. The residue was purified by silica gel chromatography to give 6-((7-ethyl-2-methylundecan-4-yl)oxy)-6-oxohexanoic acid (D33) as a white crystal (yield 73%). $^1$H NMR (400 MHz, $CDCl_3$) $\delta$ 4.94 (dtd, $J = 8.6, 6.1, 4.3$ Hz, 1H), 2.39–2.19 (m, 4H), 1.68–1.45 (m, 7H), 1.33–1.09 (m, 13H), 1.03–0.59 (m, 12H).

**6-(decyloxy)-6-oxohexanoic acid (D34):** Follow the synthesis method as above. The residue was purified by silica gel chromatography to give 6-(decyloxy)-6-oxohexanoic acid (D34) as a white crystal (yield 76%). $^1$H NMR (400 MHz, $CDCl_3$) $\delta$ 3.98 (t, $J = 6.8$ Hz, 2H), 2.40–2.13 (m, 4H), 1.60 (h, $J = 3.1$ Hz, 4H), 1.52–1.04 (m, 17H), 0.84–0.76 (m, 3H).

**6-oxo-6-(undecyloxy)hexanoic acid (D35):** Follow the synthesis method as above. The residue was purified by silica gel chromatography to give 6-oxo-6-(undecyloxy)hexanoic acid (D35) as a white crystal (yield 77%). $^1$H NMR (400 MHz, $CDCl_3$) $\delta$ 3.98 (q, $J = 6.4$ Hz, 2H), 2.28 (ddd, $J = 18.7, 8.0, 4.8$ Hz, 4H), 1.67–1.51 (m, 6H), 1.22 (dt, $J = 21.2, 6.0$ Hz, 16H), 0.94–0.68 (m, 3H).

**6-oxo-6-(undec-10-en-1-yloxy)hexanoic acid (D36):** Follow the synthesis method as above. The residue was purified by silica gel chromatography to give 6-oxo-6-(undec-10-en-1-yloxy)hexanoic acid (D36) as a white crystal (yield 66%). $^1$H NMR (400 MHz, $CDCl_3$) $\delta$ 5.79 (ddt, $J = 16.9, 10.2, 6.7$ Hz, 1H), 5.06–4.74 (m, 2H), 4.04 (t, $J = 6.8$ Hz, 2H), 2.39–2.24 (m, 4H), 2.10–1.96 (m, 2H), 1.72–1.53 (m, 7H), 1.45–1.09 (m, 14H).

#### S.6 General synthesis method of top lipid candidates

The final Ugi lipids were synthesized according to the reported method[1, 2]. Briefly, for the synthesis of the ionizable lipid library, 4CR-Ugi chemistry was employed to generate ionizable cationic lipids through reactions involving amine groups (-$NH_2$), aldehyde groups (-CHO), carboxylic acids (-COOH), and isocyanide groups (-NC). The amines, isocyanides, aldehyde tails, and carboxylic acids were sourced from TCI and Sigma Aldrich or synthesized following previously reported methods. In brief, the reactants were dissolved in ethanol and reacted in a round-bottom flask for 18 hours at a molar ratio of 1:1:1:1 (aldehyde:amine:isocyanide:carboxylic acid). The reaction mixture was evaporated under vacuum. The residue was purified using a CombiFlash BUCHI C-815 chromatography system with gradient elution (1% ammonia water, 0%–10% MeOH in DCM) to isolate the final product.

**LUMI-1:** Follow the synthesis method as above. The residue was purified by silica gel chromatography to give LUMI-1 as a colorless oil (yield 85%). MS (ESI) $m/z$: $[M+H]^+$ calcd. for $C_{40}H_{75}BrN_5O_2$, 737.97; found, 737.50. $^1$H NMR (400 MHz, $CDCl_3$) $\delta$ 5.49–5.23 (m, 4H), 3.57–3.31 (m, 3H), 2.77 (s, 1H), 2.22 (s, 2H), 2.12–1.88 (m, 16H), 1.84 (s, 1H), 1.67 (s, 6H), 1.41–1.12 (m, 34H), 0.89 (t, $J = 6.8$ Hz, 3H).

**LUMI-2:** Follow the synthesis method as above. The residue was purified by silica gel chromatography to give LUMI-2 as a colorless oil (yield 82%). MS (ESI) $m/z$: $[M+H]^+$ calcd. for $C_{42}H_{76}BrN_3O_2$, 734.51; found, 734.92. $^1$H NMR (400 MHz, $CDCl_3$) $\delta$ 5.42–5.30 (m, 2H), 3.39 (td, $J = 6.9, 1.2$ Hz, 4H), 2.50–2.43 (m, 2H), 2.34 (s, 7H), 1.93 (d, $J = 2.9$ Hz, 5H), 1.64 (d, $J = 6.1$ Hz, 6H), 1.48–1.21 (m, 40H), 0.86 (dd, $J = 7.0, 1.2$ Hz, 3H).

**LUMI-3:** Follow the synthesis method as above. The residue was purified by silica gel chromatography to give LUMI-3 as a colorless oil (yield 81%). MS (ESI) $m/z$: $[M+H]^+$ calcd. for $C_{39}H_{71}BrN_3O_4$, 724.91; found, 724.75. $^1$H NMR (400 MHz, $CDCl_3$) $\delta$ 4.10 (q, $J = 7.1$ Hz, 6H), 3.40 (t, $J = 6.8$ Hz, 8H), 2.55 (d, $J = 1.7$ Hz, 2H), 2.33 (t, $J = 7.4$ Hz, 6H), 2.03 (s, 6H), 1.89–1.83 (m, 6H), 1.66–1.61 (m, 10H), 1.50–1.45 (m, 6H), 1.25 (d, $J = 7.2$ Hz, 12H), 0.86 (td, $J = 7.0, 2.0$ Hz, 6H).

**LUMI-4:** Follow the synthesis method as above. The residue was purified by silica gel chromatography to give LUMI-4 as a colorless oil (yield 88%). MS (ESI) $m/z$: $[M+H]^+$ calcd. for $C_{41}H_{73}BrN_4O_2$, 733.97; found, 733.58. $^1$H NMR (400 MHz, $CDCl_3$) $\delta$ 5.39–5.18 (m, 2H), 2.52 (ddd, $J = 69.7, 6.6, 4.2$ Hz, 14H), 2.08–1.74 (m, 13H), 1.62 (q, $J = 4.7$ Hz, 10H), 1.35–1.04 (m, 28H), 0.96–0.61 (m, 3H).

**LUMI-5:** Follow the synthesis method as above. The residue was purified by silica gel chromatography to give LUMI-5 as a colorless oil (yield 80%). MS (ESI) $m/z$: $[M+H]^+$ calcd. for $C_{41}H_{74}BrN_5O_2$, 748.50; found, 748.50. $^1$H NMR (400 MHz, $CDCl_3$) $\delta$ 5.33 (d, $J = 7.3$ Hz, 4H), 2.85 – 2.61 (m, 12H), 2.08 – 1.91 (m, 20H), 1.31 – 1.24 (m, 27H), 0.87 (d, J = 2.8 Hz, 3H).

**LUMI-6:** Follow the synthesis method as above. The residue was purified by silica gel chromatography to give LUMI-6 as a colorless oil (yield 81%). MS (ESI) $m/z$: $[M+H]^+$ calcd. for $C_{45}H_{81}BrN_4O_4$, 821.54; found, 821.67. $^1$H NMR (400 MHz, $CDCl_3$) $\delta$ 6.67 – 6.42 (m, 1H), 4.04 (td, $J = 6.7$, 4.0 Hz, 2H), 3.40 (td, $J = 6.7$, 3.0 Hz, 3H), 2.84 – 2.39 (m, 7H), 2.37 – 2.12 (m, 4H), 1.97 (d, $J = 2.9$ Hz, 4H), 1.92 – 1.84 (m, 2H), 1.75 – 1.19 (m, 41H), 0.91 – 0.81 (m, 6H).

**LUMI-6D:** Follow the synthesis method as above. The residue was purified by silica gel chromatography to give LUMI-6D as a colorless oil (yield 75%). MS (ESI) $m/z$: $[M+H]^+$ calcd. for $C_{46}H_{84}N_4O_4$ 757.65, found 757.50.$^1$H NMR (400 MHz, $CDCl_3$) $\delta$ 6.61 (s, 1H), 4.00 (t, $J = 6.7$ Hz, 2H), 3.33 (q, $J = 5.7$ Hz, 2H), 2.67 (t, $J = 6.2$ Hz, 1H), 2.49 (t, $J = 6.0$ Hz, 9H), 2.30 – 2.20

(m, 2H), 2.19 – 2.06 (m, 2H), 2.03 (t, $J$ = 3.2 Hz, 3H), 1.69 – 1.12 (m, 49H), 0.83 (dtd, $J$ = 7.0, 4.4, 2.2 Hz, 9H).

#### S.7 Antibody and Dye used in the flow cytometry and immunofluorescence

Supplementary Table S4: Antibodies and dyes used for flow cytometry staining and immunofluorescence staining.

| **Antibody/Dye** | **Conjugate** | **Source (Cat #)** |
|---|---|---|
| **Flow Cytometry Staining** | | |
| TruStain FcX™ blocker | - | BioLegend (Cat # 101320) |
| CD31 Antibody | Alexa Fluor 594 | BioLegend (Cat # 102520) |
| EpCAM Antibody | BrilliantViolet 421 | BioLegend (Cat # 118225) |
| Zombie NIR™ Fixable Viability Kit | - | BioLegend (Cat # 423106) |
| Fix/Perm Kit | - | BD Biosciences (Cat # 554714) |
| **Immunofluorescence Staining** | | |
| Acetylated Tubulin Antibody | Alexa Fluor 647 | Santa Cruz (Cat # sc-23950) |
| CC10 Antibody | Alexa Fluor 488 | Santa Cruz (Cat # sc-390313) |
| DAPI | - | Invitrogen (Cat # P36931) |

### References

[1] Shufen Xu et al. "Tumor-Tailored Ionizable Lipid Nanoparticles Facilitate IL-12 Circular RNA Delivery for Enhanced Lung Cancer Immunotherapy". In: *Advanced Materials* (2024), p. 2400307.

[2] Zepeng He et al. "A Multidimensional Approach to Modulating Ionizable Lipids for High-Performing and Organ-Selective mRNA Delivery". In: *Angewandte Chemie International Edition* 62.43 (2023), e202310401.

### Supplementary Figures

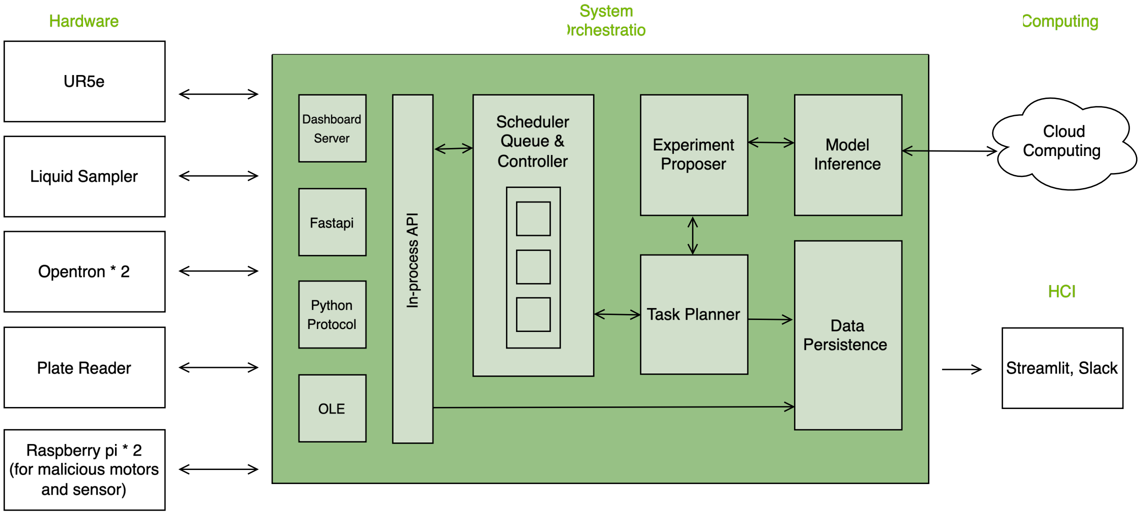

Supplementary Figure S1: The diagram of software modules and their communications with LUMI-model, and the hardware modules.

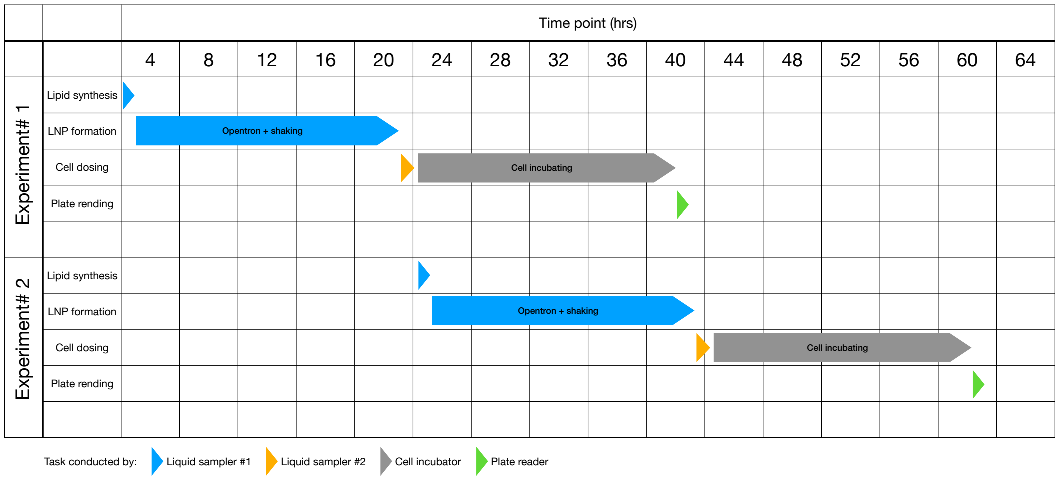

Supplementary Figure S2: The timeline of each experiment in LUMI-lab and the shift of tasks between experiments. Two experiments can be conducted simultaneously with different hardware modules on distinct tasks. This strategy increases the overall experiment throughput of the system.

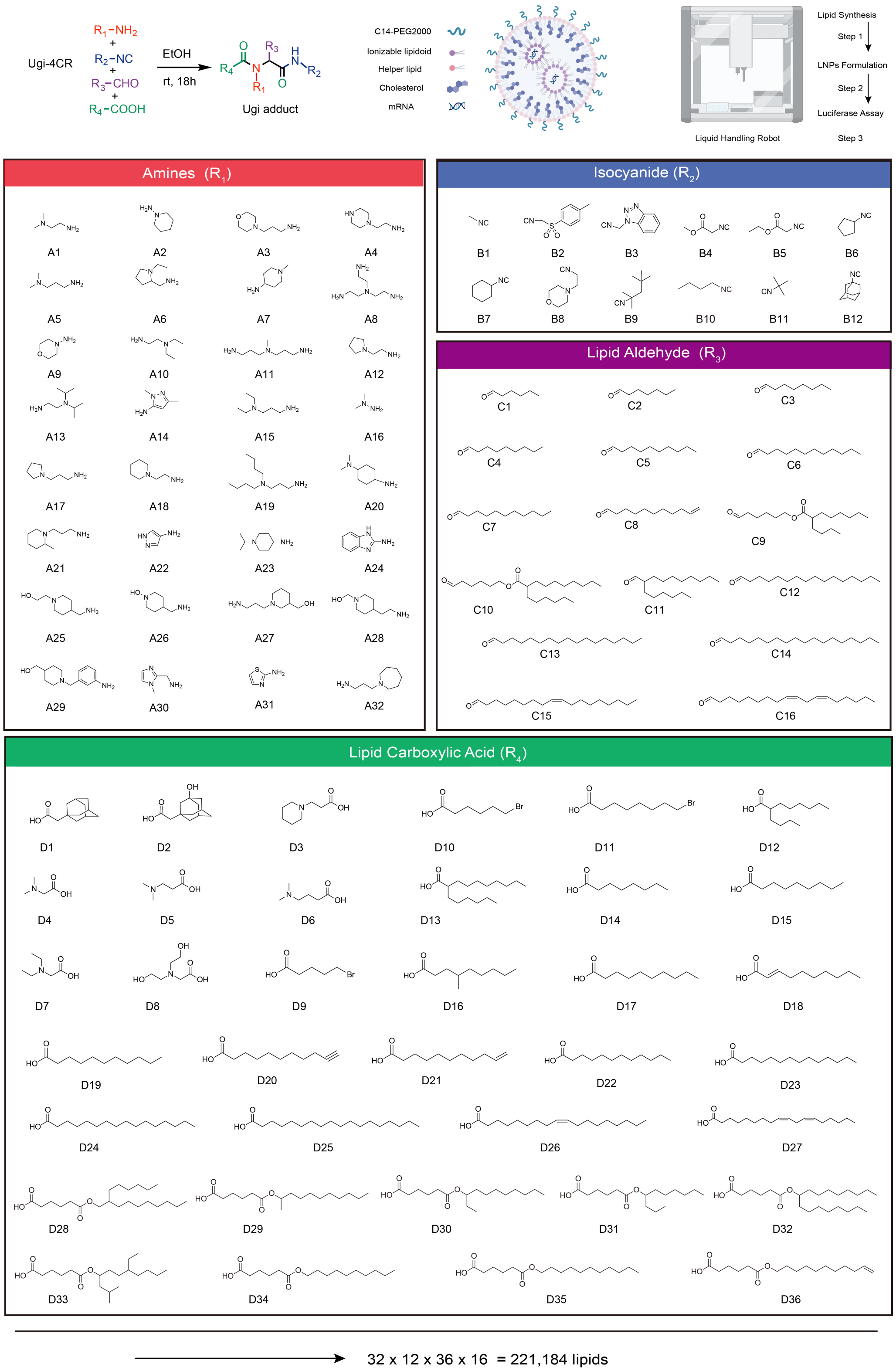

Supplementary Figure S3: The chemical library for building the Ugi-4CR reaction used in lipid synthesis.

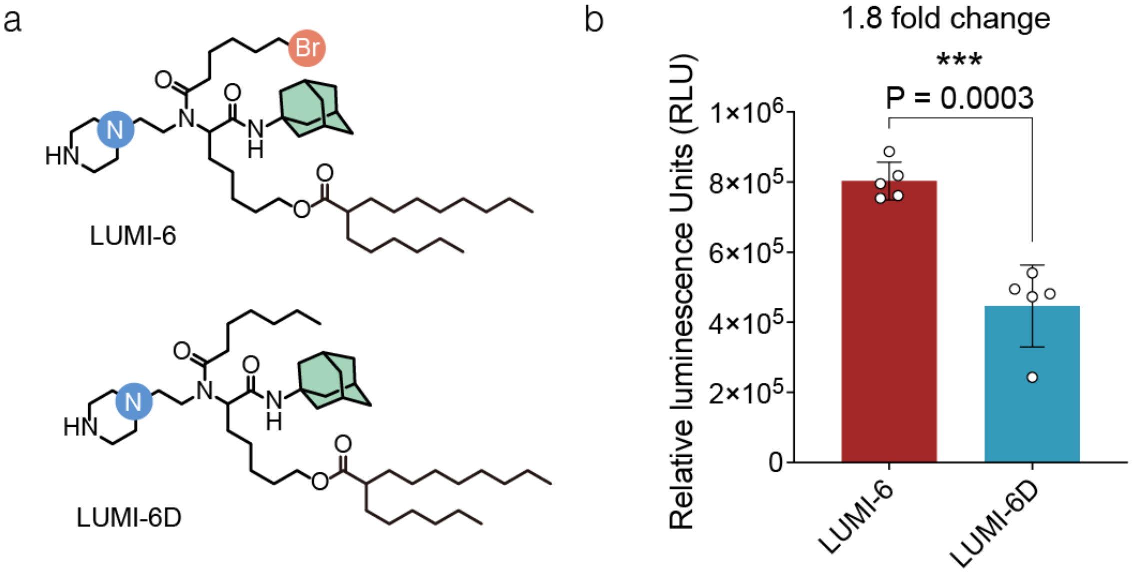

Supplementary Figure S4: Head to head comparison between brominated and non-brominated ionizable lipids. **(a)** The structures of LUMI-6 and LUMI-6 debrominated (LUMI-6D) ionizable lipids. **(b)** The relative luminescence units between LUMI-6 and LUMI-6D LNPs in HBE cells after 18 hours incubation. Statistical significance evaluated using a two-tailed unpaired $t$-test ($*P < 0.05$; $**P < 0.01$; $***P < 0.001$; $****P < 0.0001$).

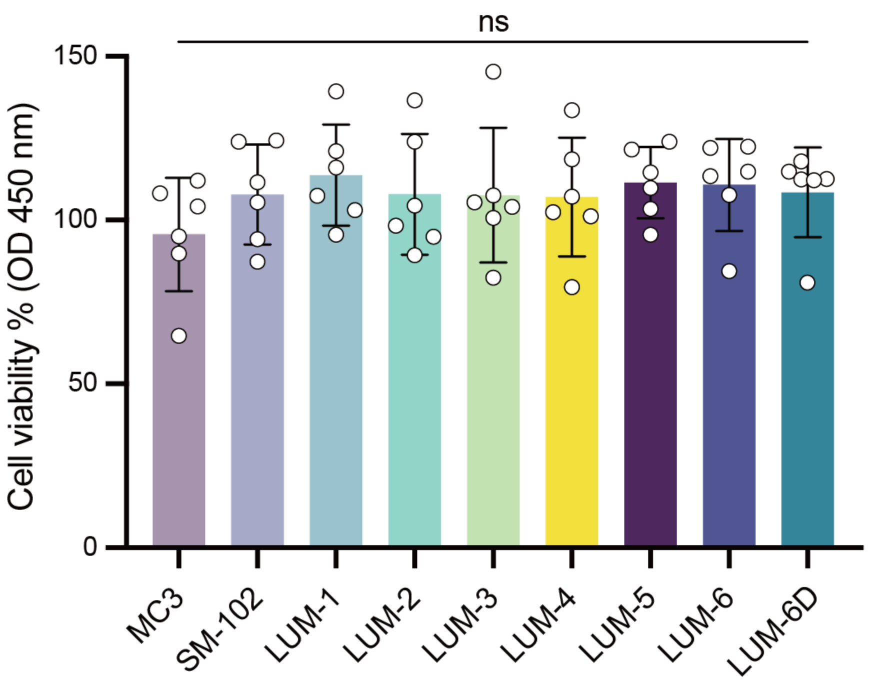

Supplementary Figure S5: The cytotoxicity of both brominated and non-brominated lipids was assessed using the CCK-8 assay. Statistical significance evaluated using two-way ANOVA (*$P <$ 0.05; **$P < 0.01$; ***$P < 0.001$; ****$P < 0.0001$).

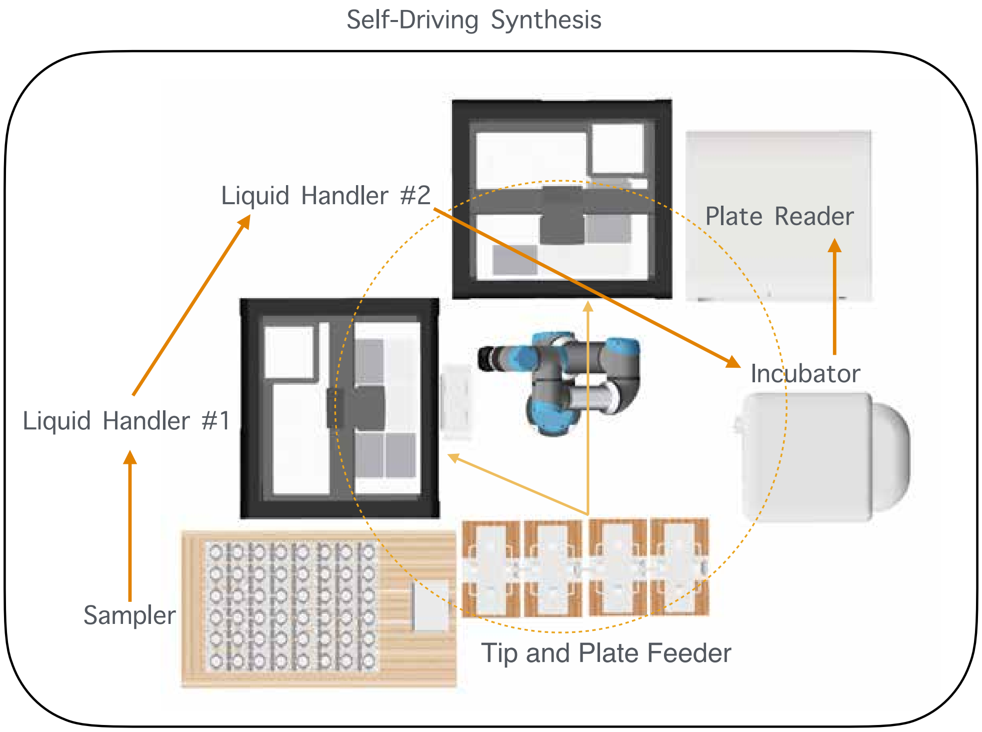

Supplementary Figure S6: Top view of the LUMI-lab hardware components and setup. Arrows indicate the directions of transferring materials and reagents between components.

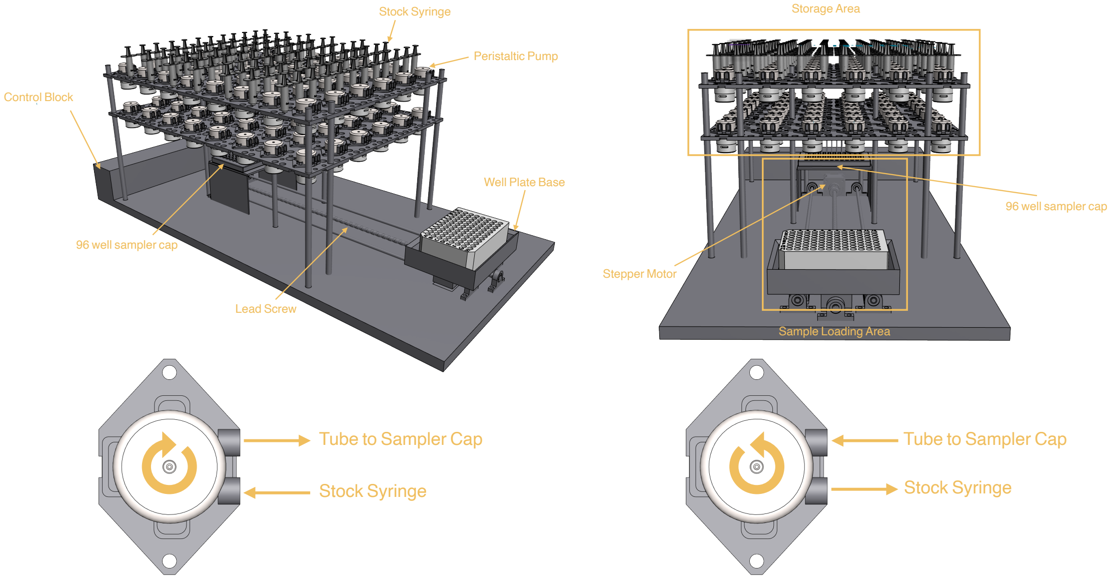

Supplementary Figure S7: The 3D design of the liquid sampler module in LUMI-lab.

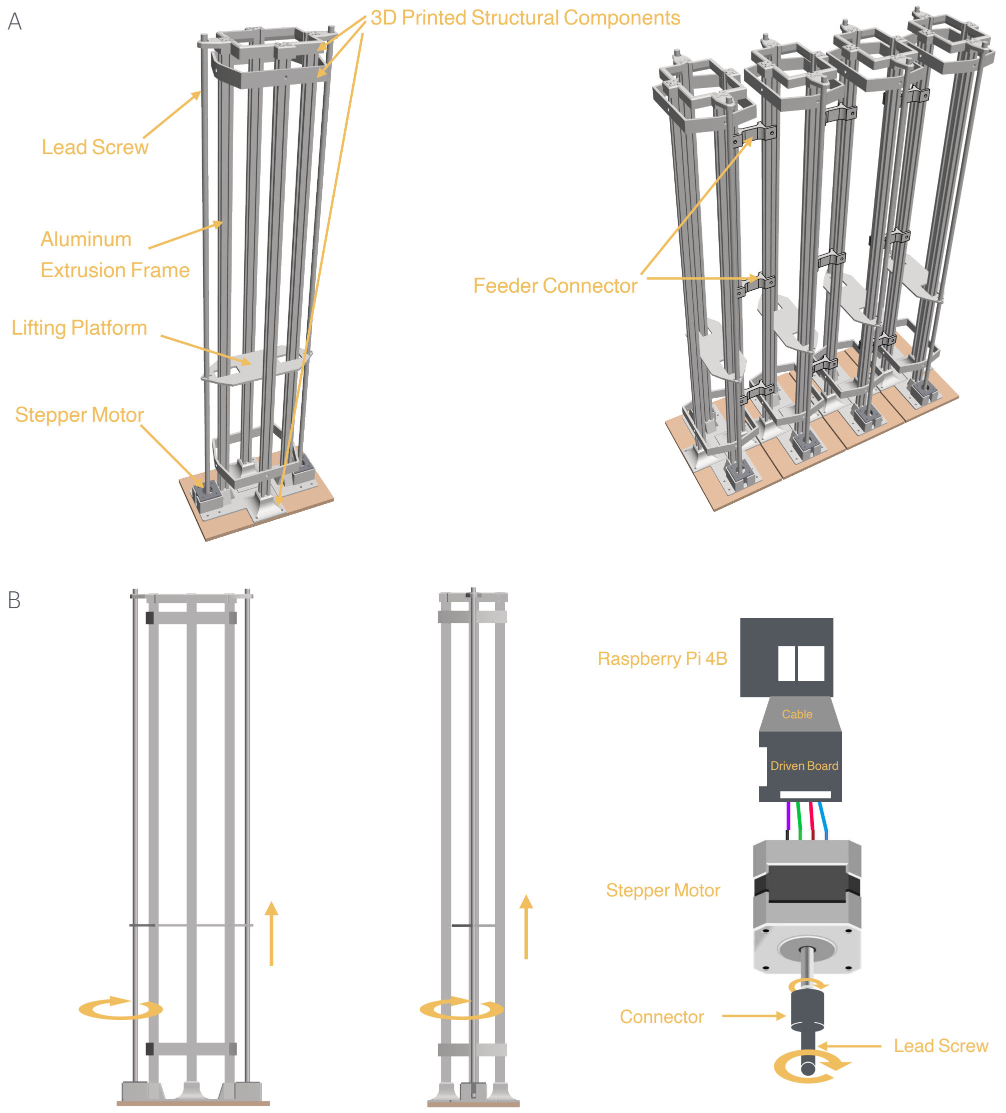

Supplementary Figure S8: The 3D design of the feeders for pipette tip racks and well plates in LUMI-lab. **(a)** The composition of single feeder and multiple feeders in a group. **(b)** Diagram visualizing the control and movement of the feeder.

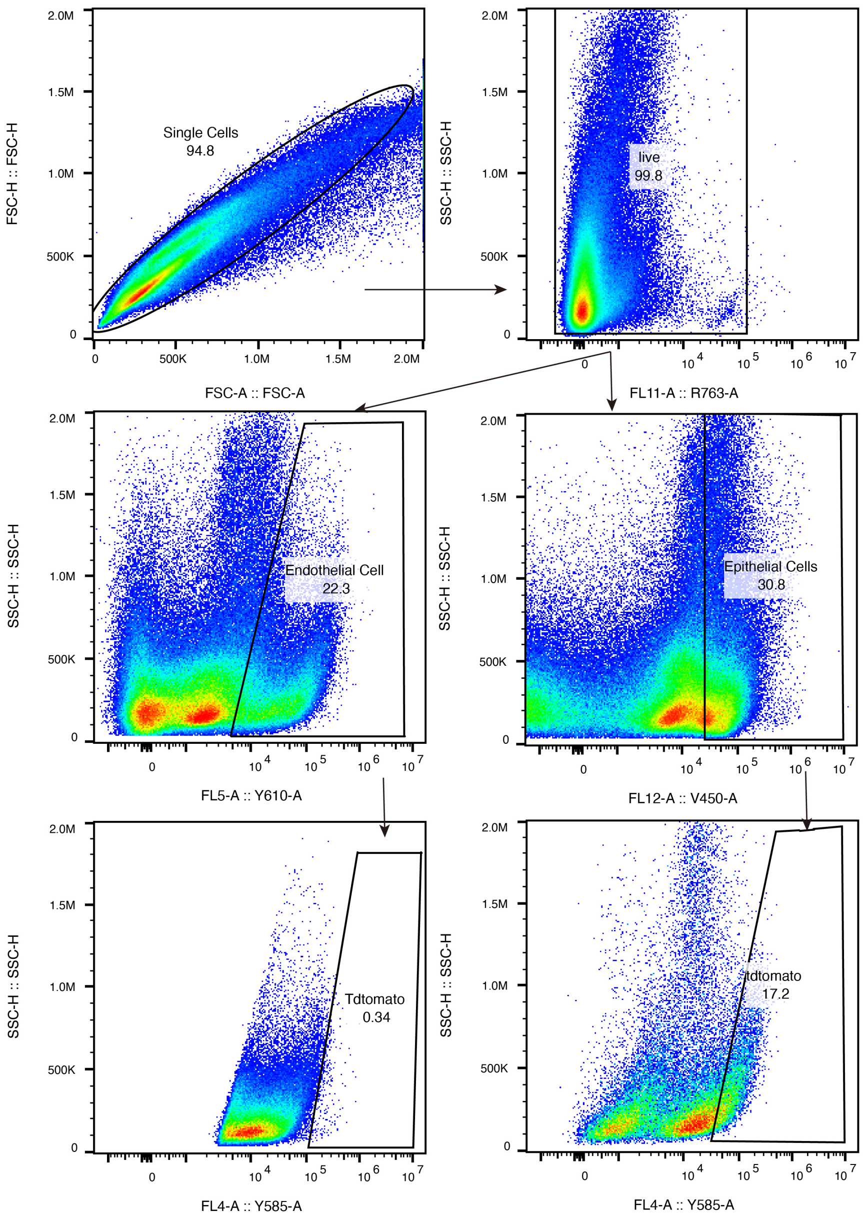

Supplementary Figure S9: Flow cytometry gating strategy for lung endothelial and epithelial cells in Ai9 reporter mice.

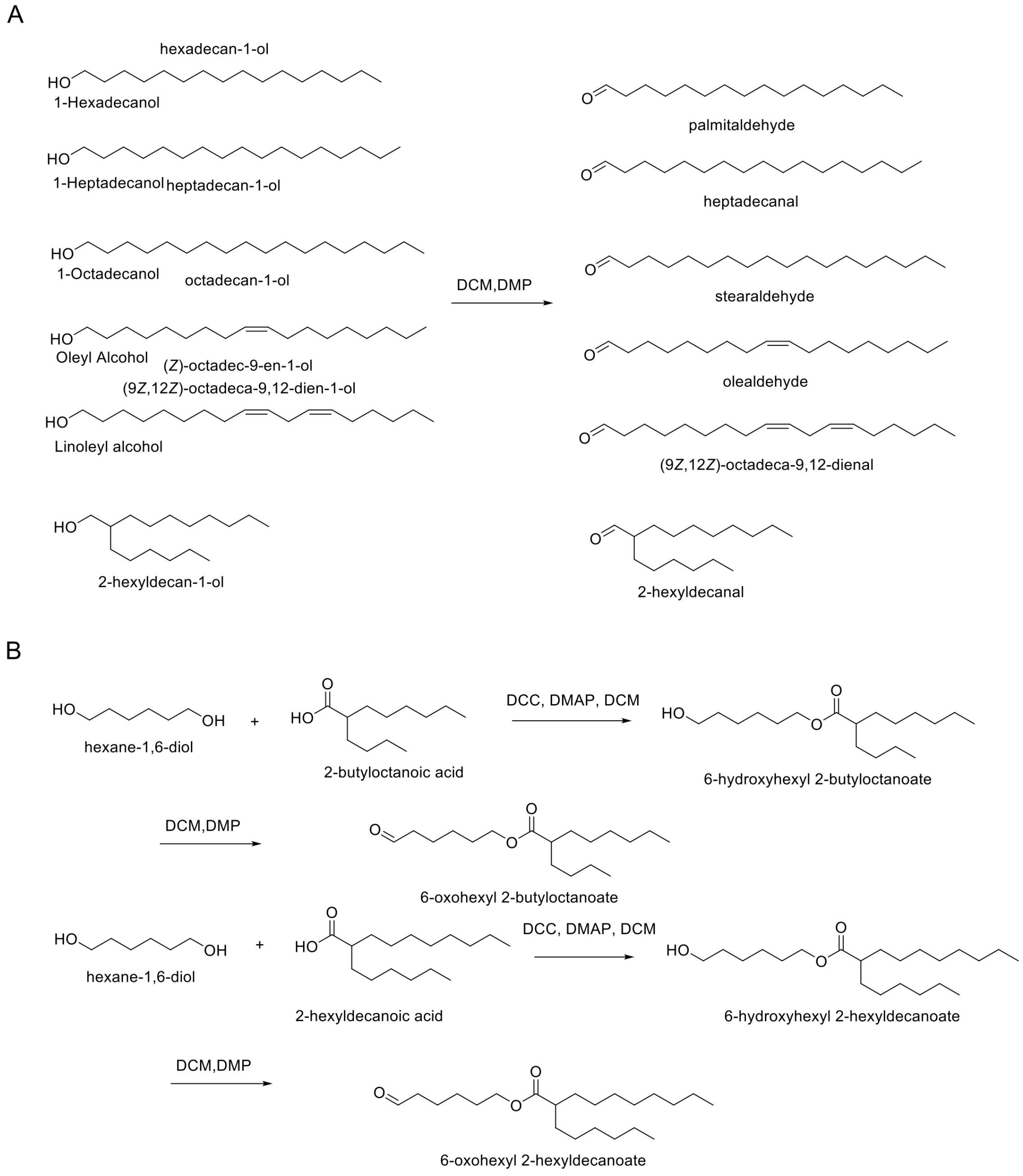

Supplementary Figure S10: General synthesis method for aldehyde tail (C9-C16), which are included in the chemical reagent library for LUMI-lab experiments.

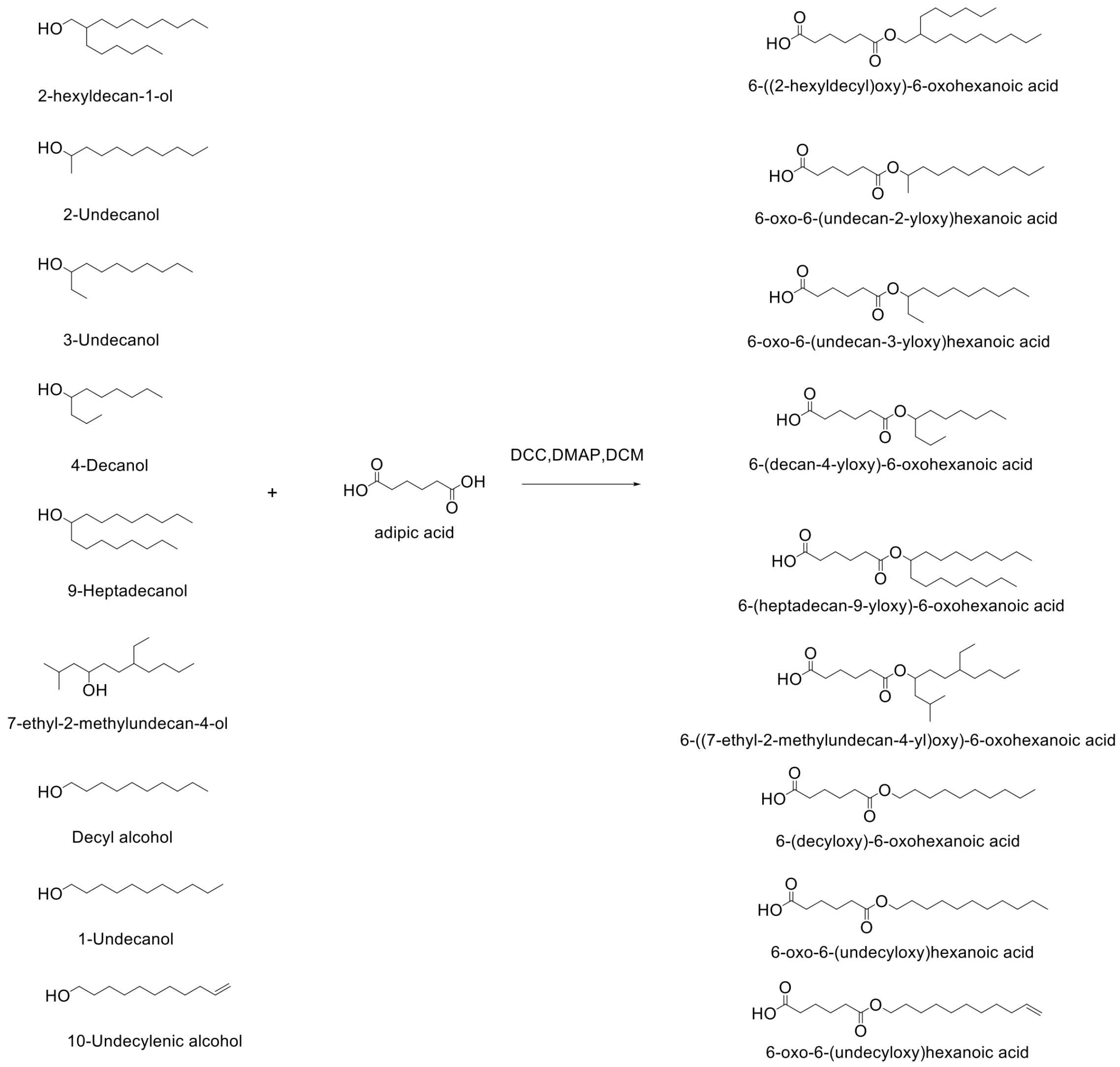

Supplementary Figure S11: General synthesis method for carboxylic acid tail (D28-D36), which are included in the chemical reagent library for LUMI-lab experiments.

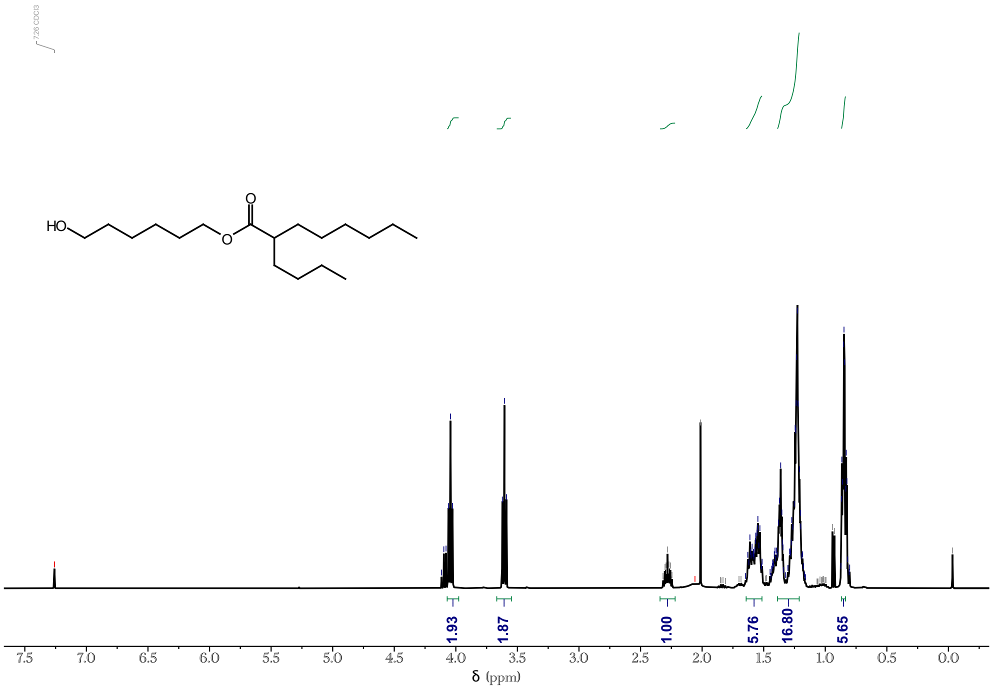

Supplementary Figure S12: ¹H NMR for tail C9-OH.

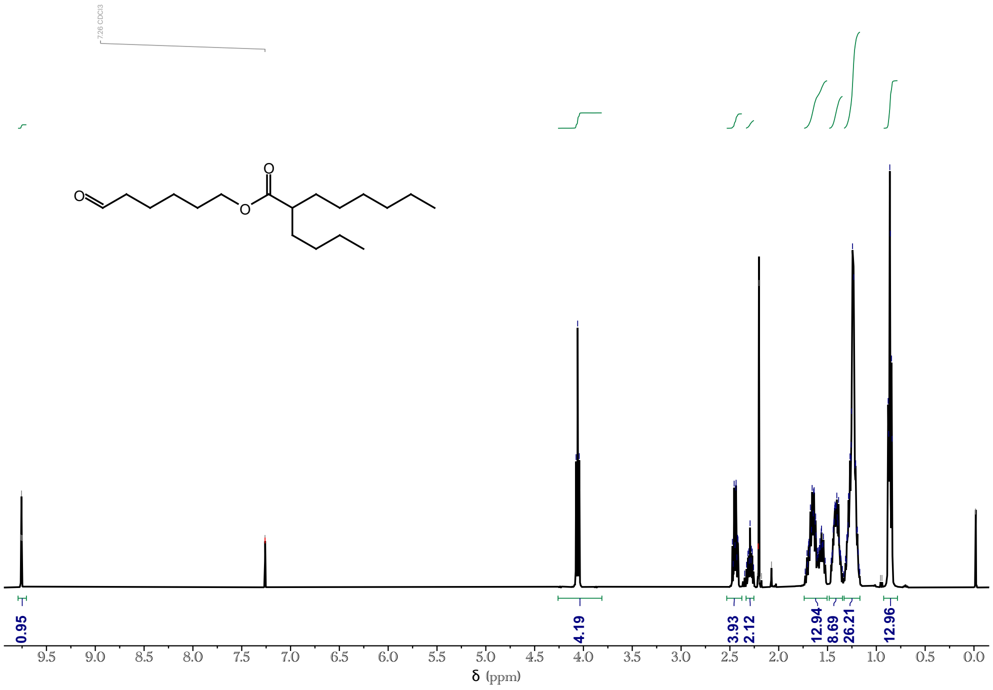

Supplementary Figure S13: ¹H NMR for tail C9-CHO.

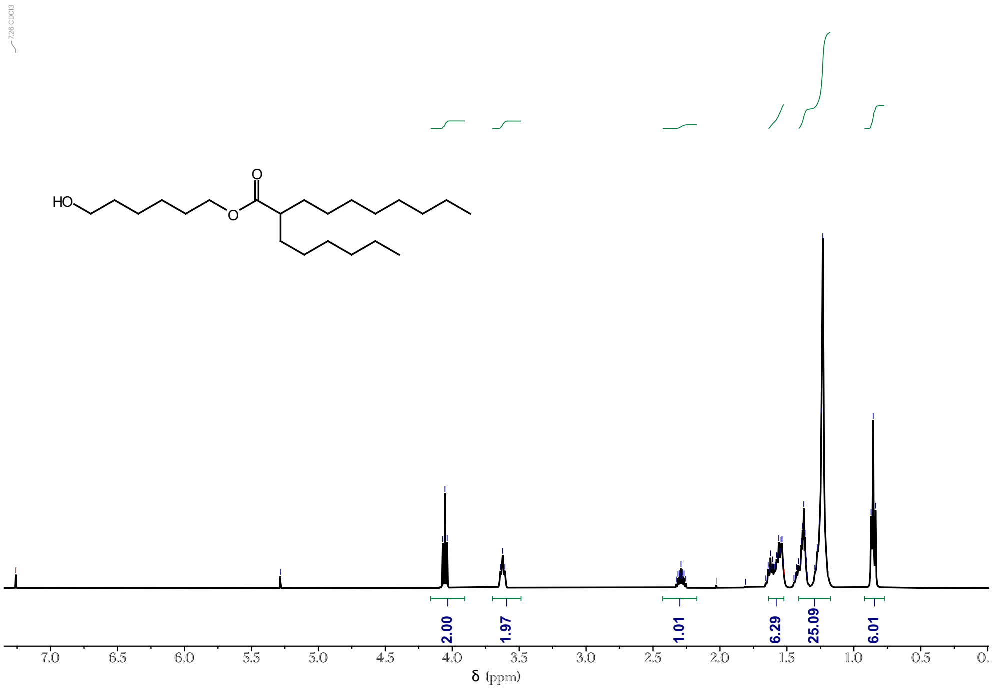

Supplementary Figure S14: $^1$H NMR for tail C10-OH.

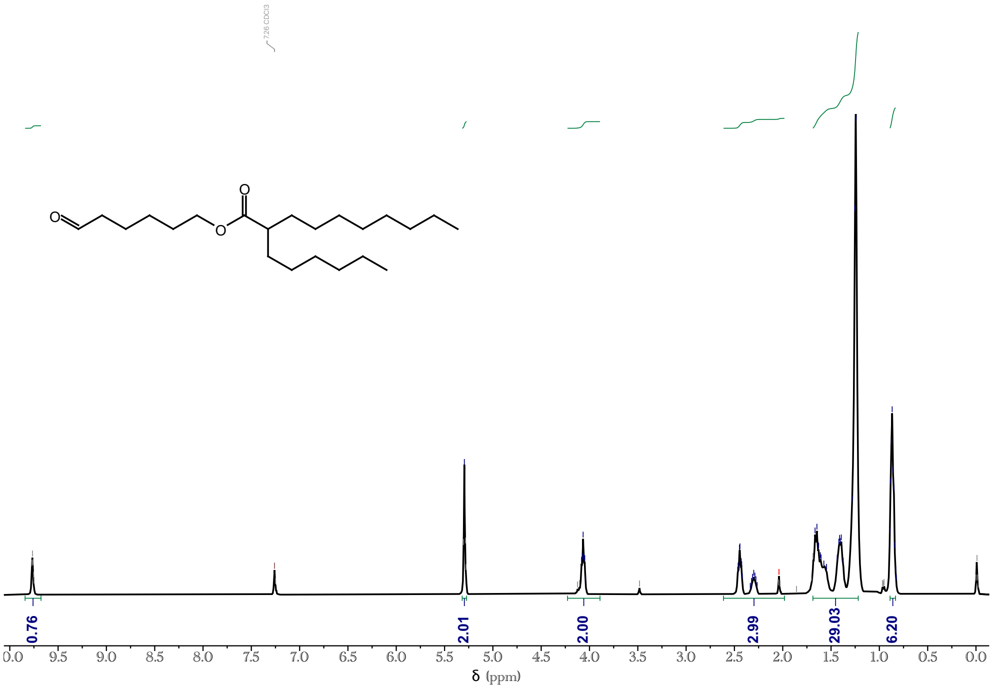

Supplementary Figure S15: $^1$H NMR for tail C10-CHO.

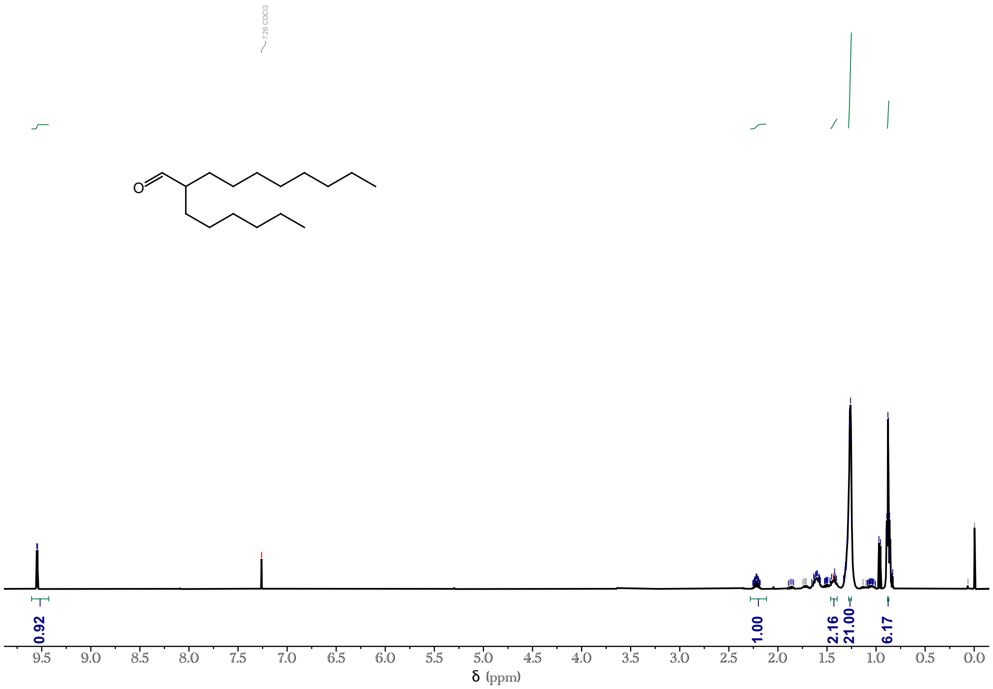

Supplementary Figure S16: $^1$H NMR for tail C11-CHO.

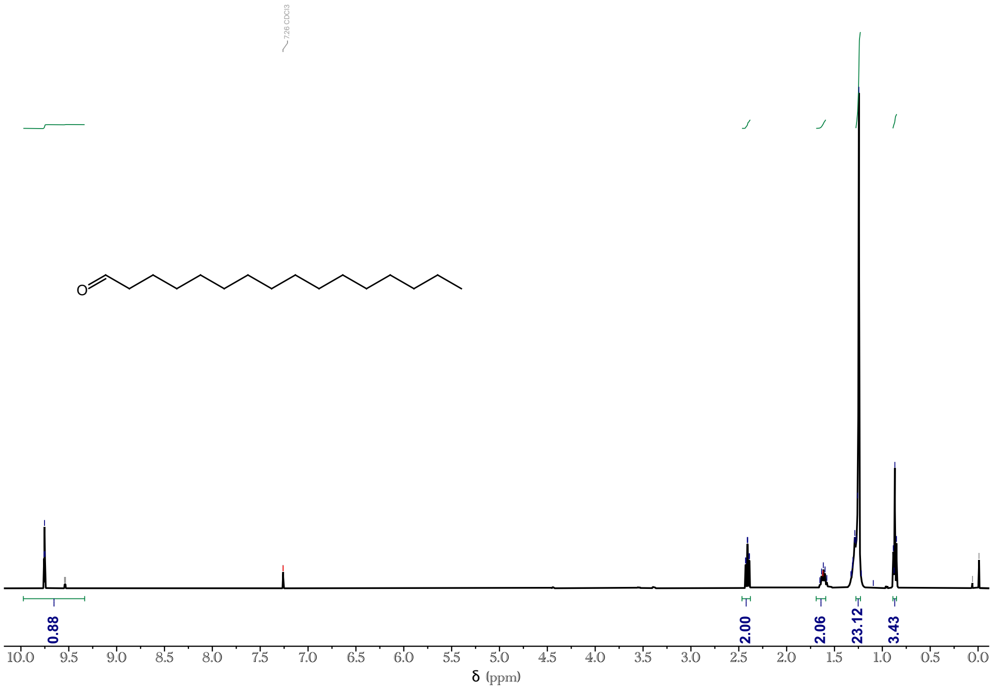

Supplementary Figure S17: $^1$H NMR for tail C12-CHO.

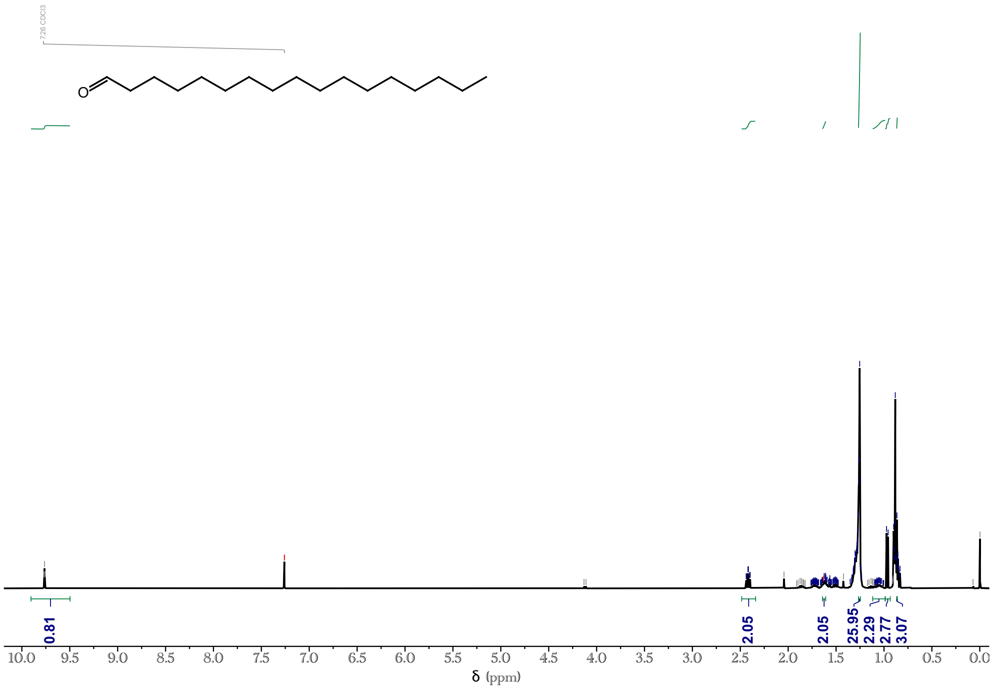

Supplementary Figure S18: $^{1}$H NMR for tail C13-CHO.

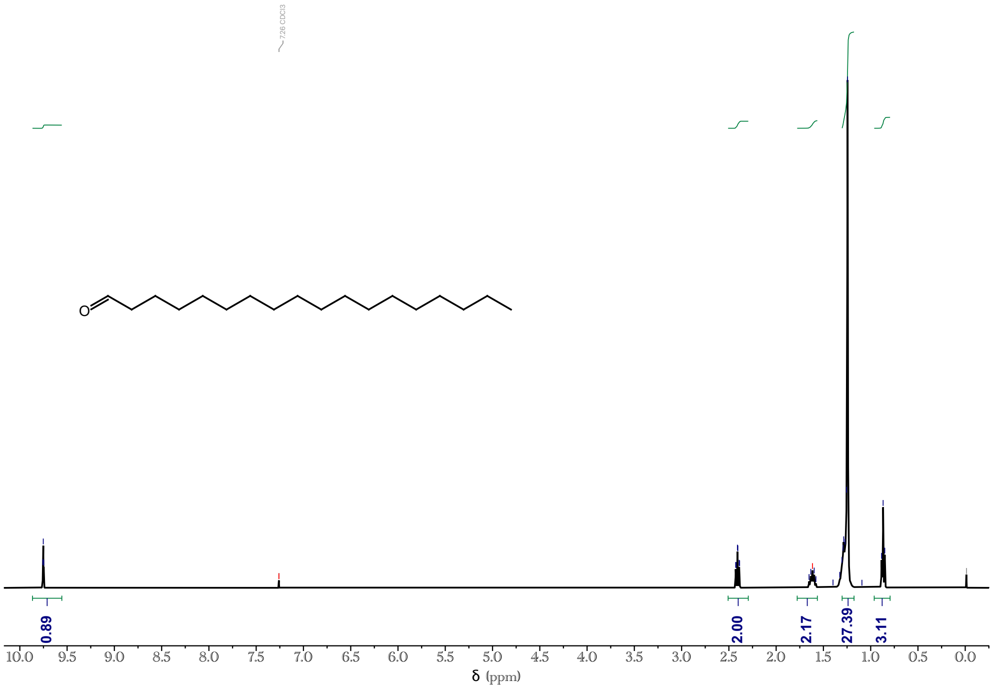

Supplementary Figure S19: $^{1}$H NMR for tail C14-CHO.

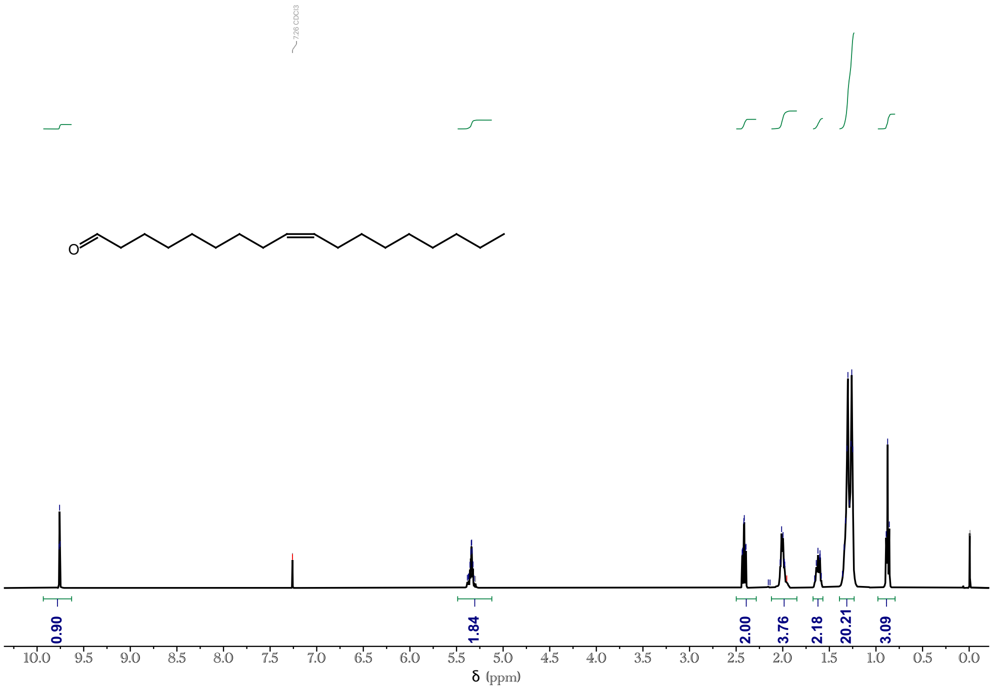

Supplementary Figure S20: $^{1}$H NMR for tail C15-CHO.

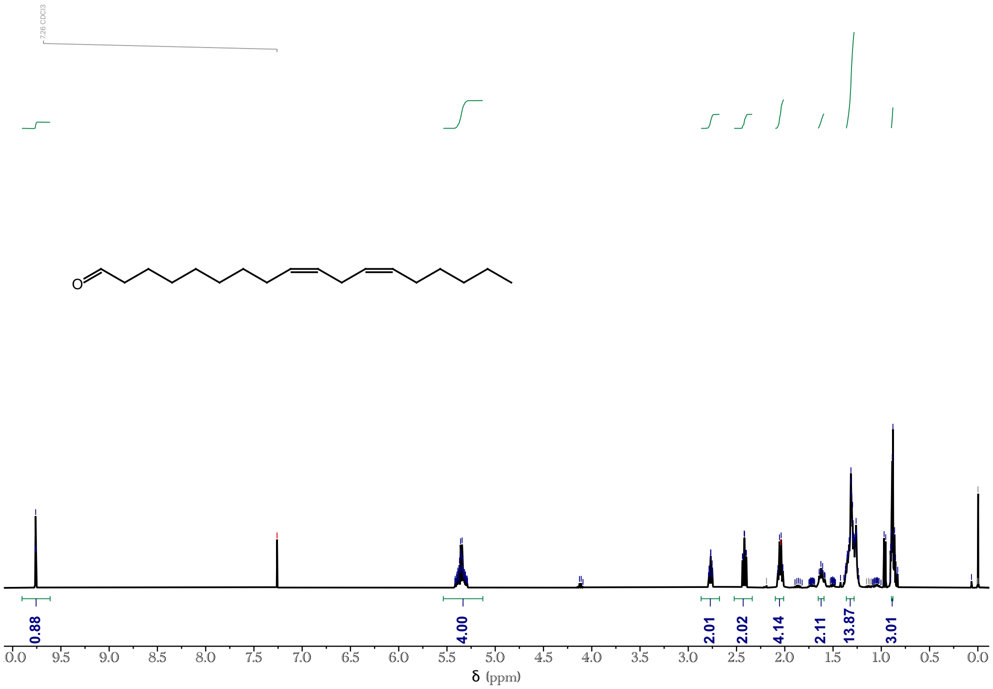

Supplementary Figure S21: $^{1}$H NMR for tail C16-CHO.

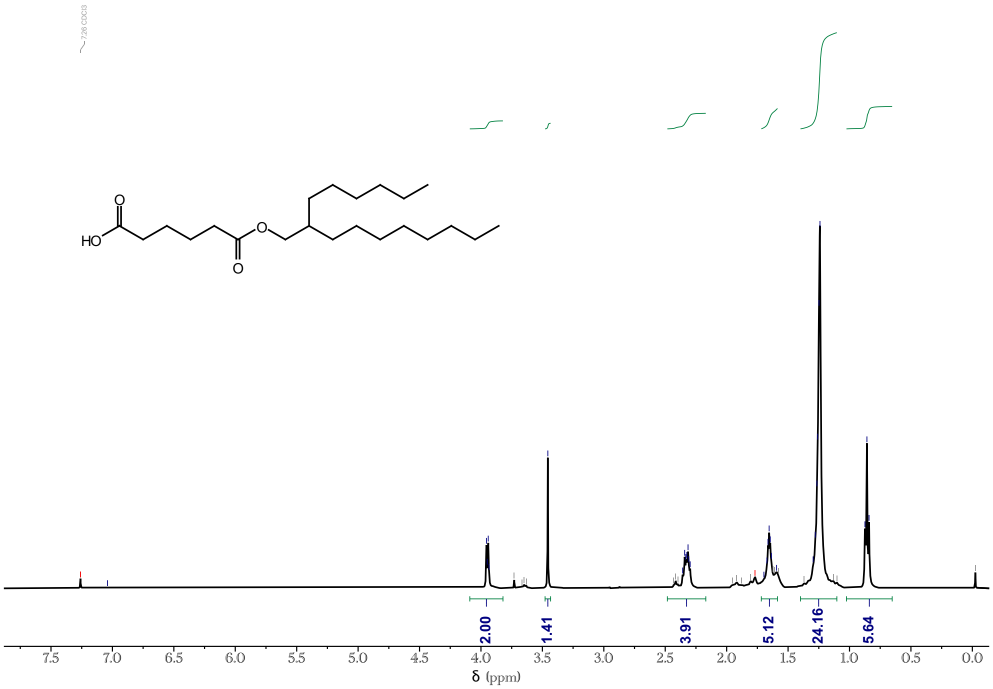

Supplementary Figure S22: $^{1}$H NMR for tail D28.

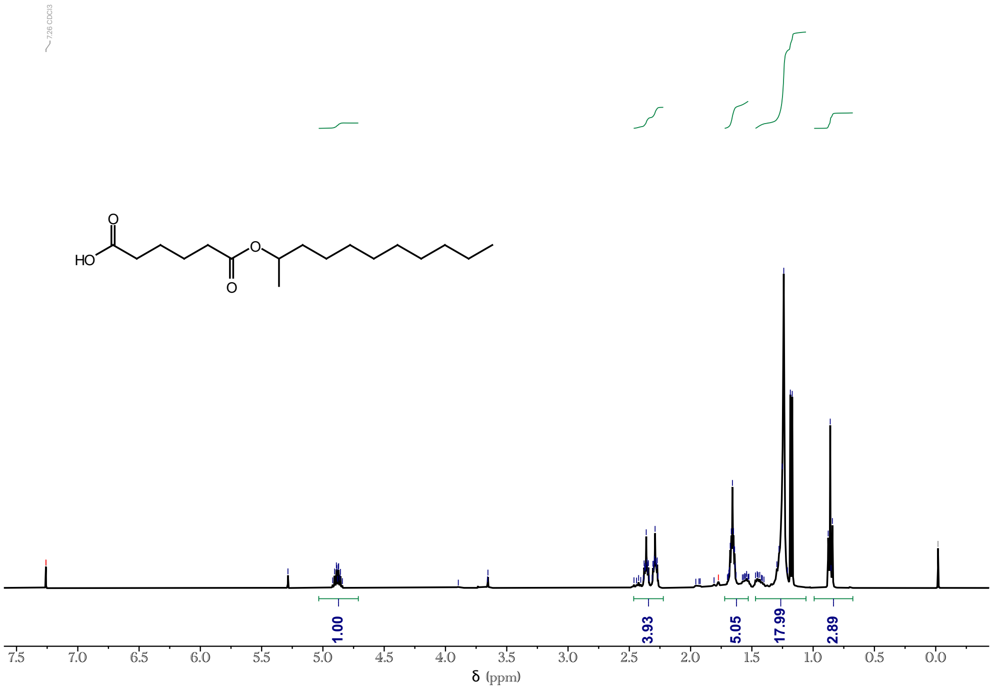

Supplementary Figure S23: $^{1}$H NMR for tail D29.

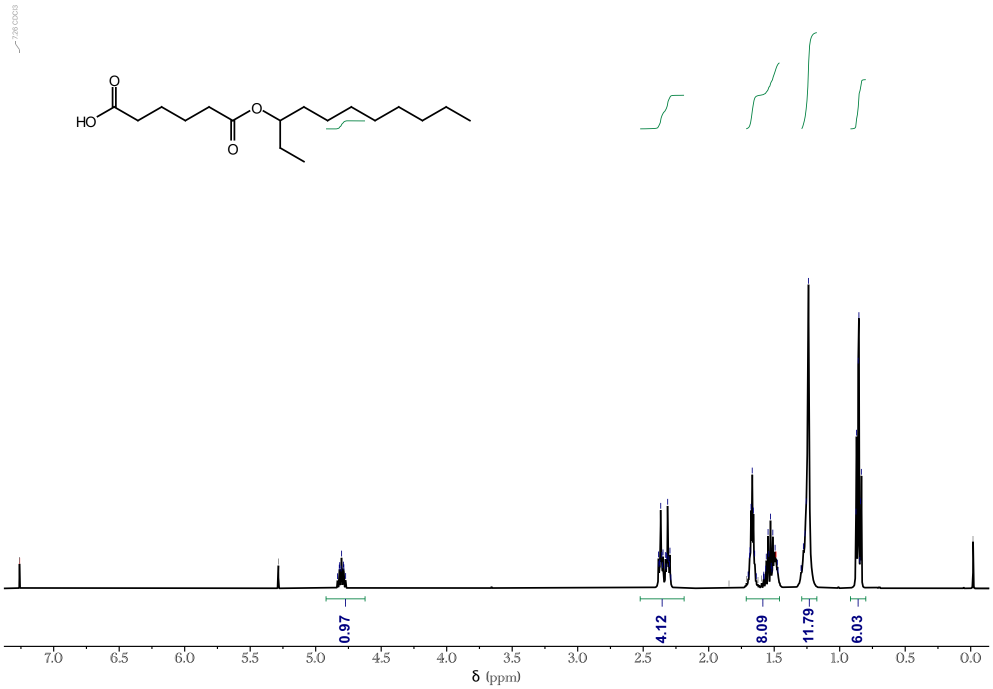

Supplementary Figure S24: $^1$H NMR for tail D30.

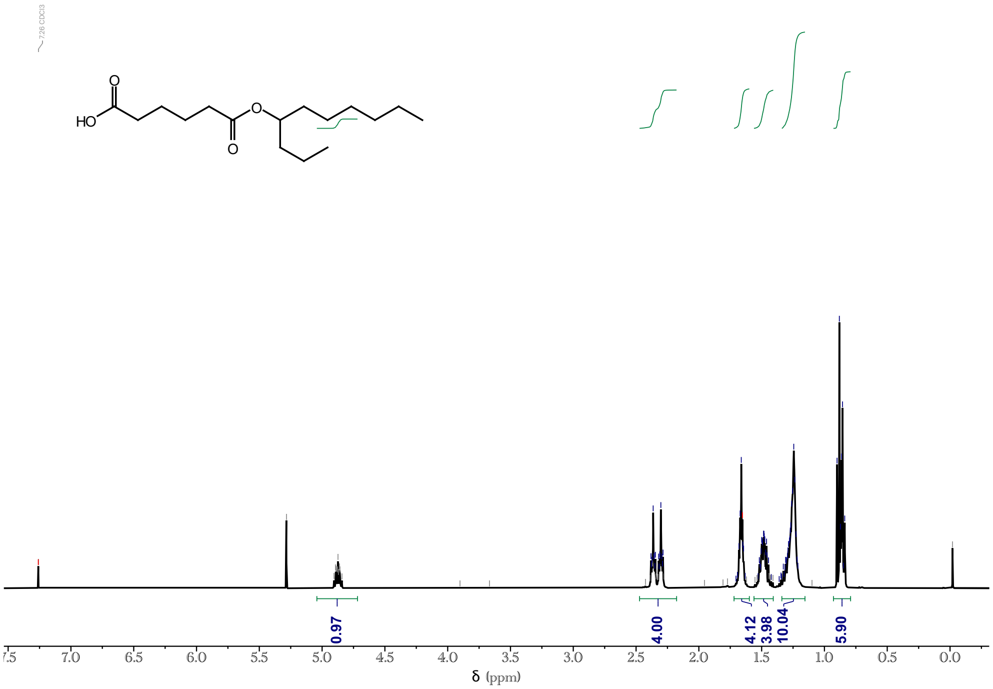

Supplementary Figure S25: $^1$H NMR for tail D31.

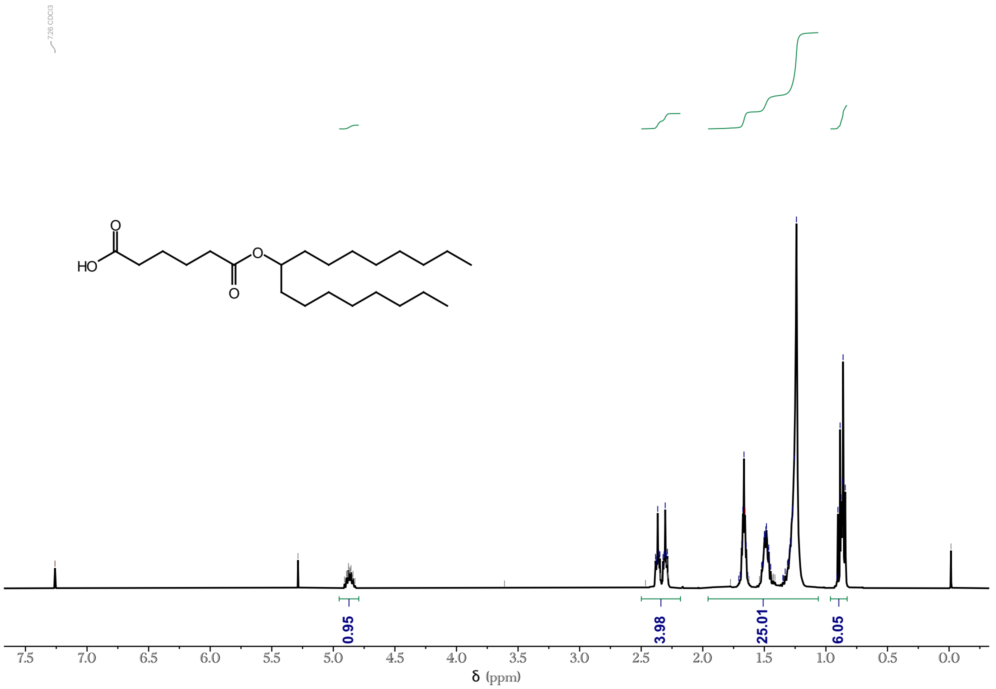

Supplementary Figure S26: $^{1}$H NMR for tail D32.

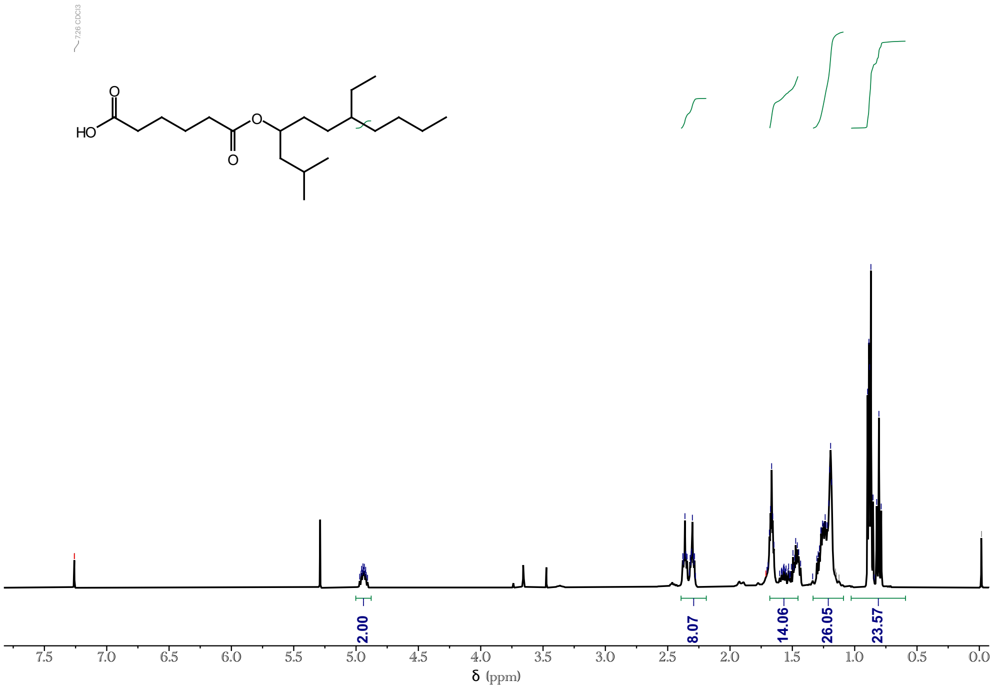

Supplementary Figure S27: $^{1}$H NMR for tail D33.

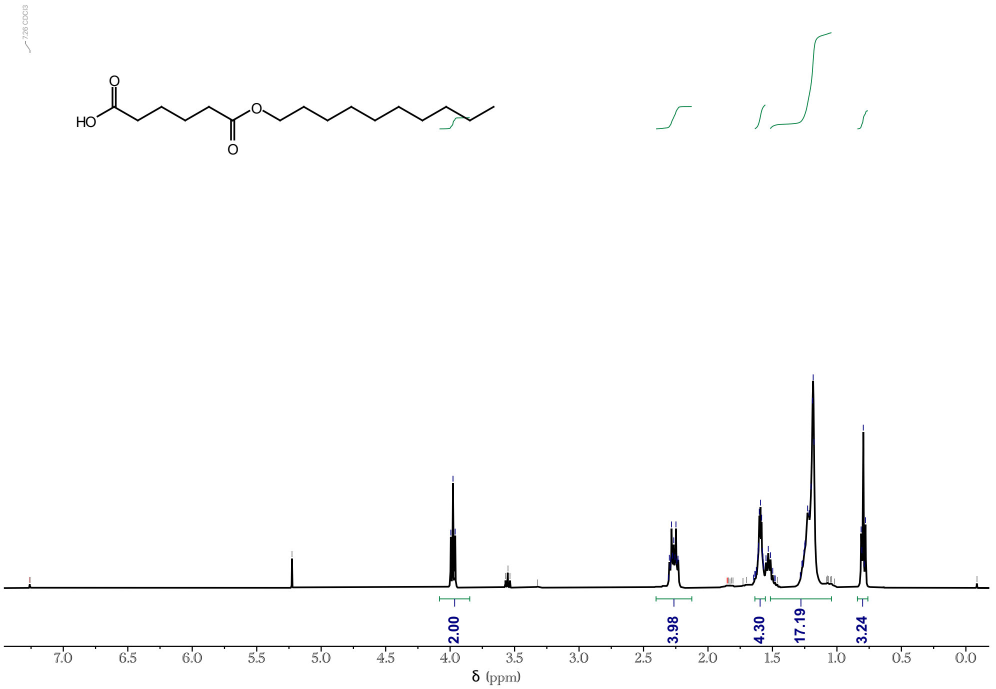

Supplementary Figure S28: $^{1}$H NMR for tail D34.

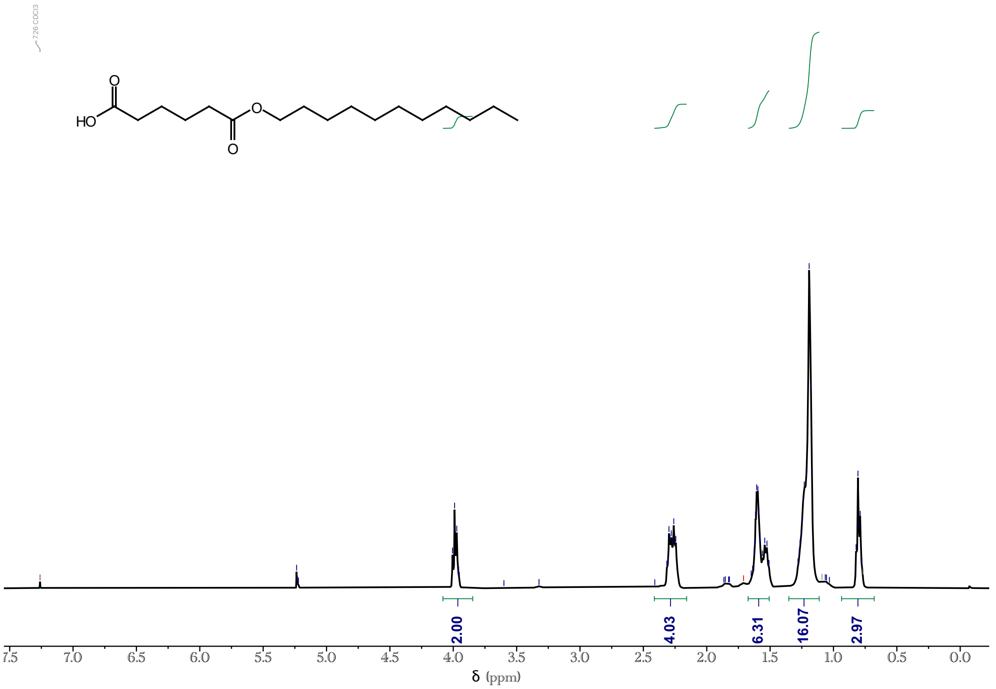

Supplementary Figure S29: $^{1}$H NMR for tail D35.

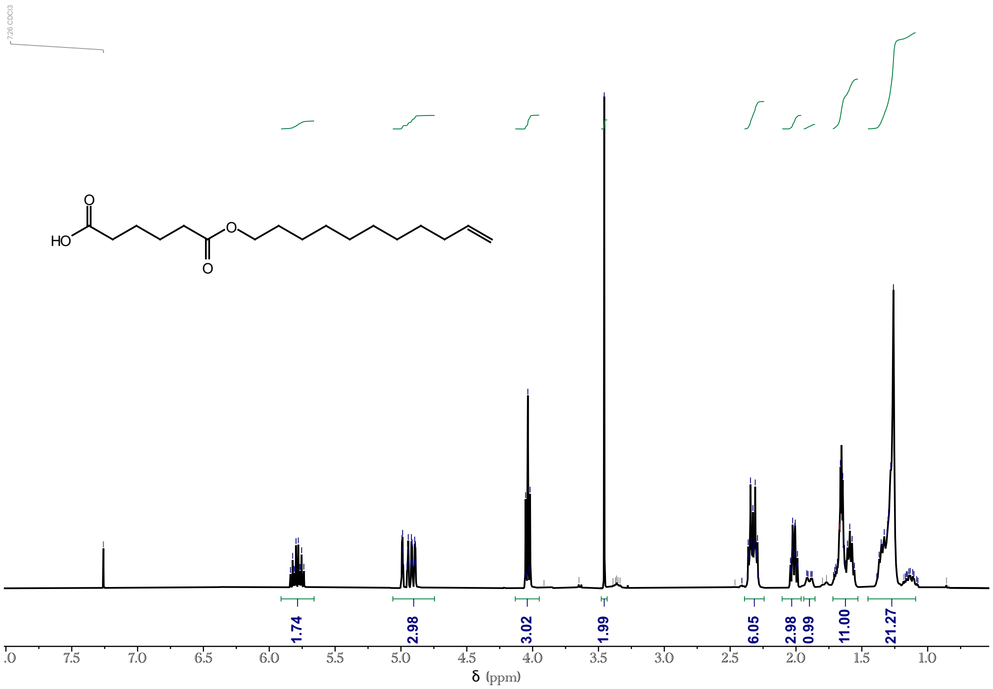

Supplementary Figure S30: $^1$H NMR for tail D36.

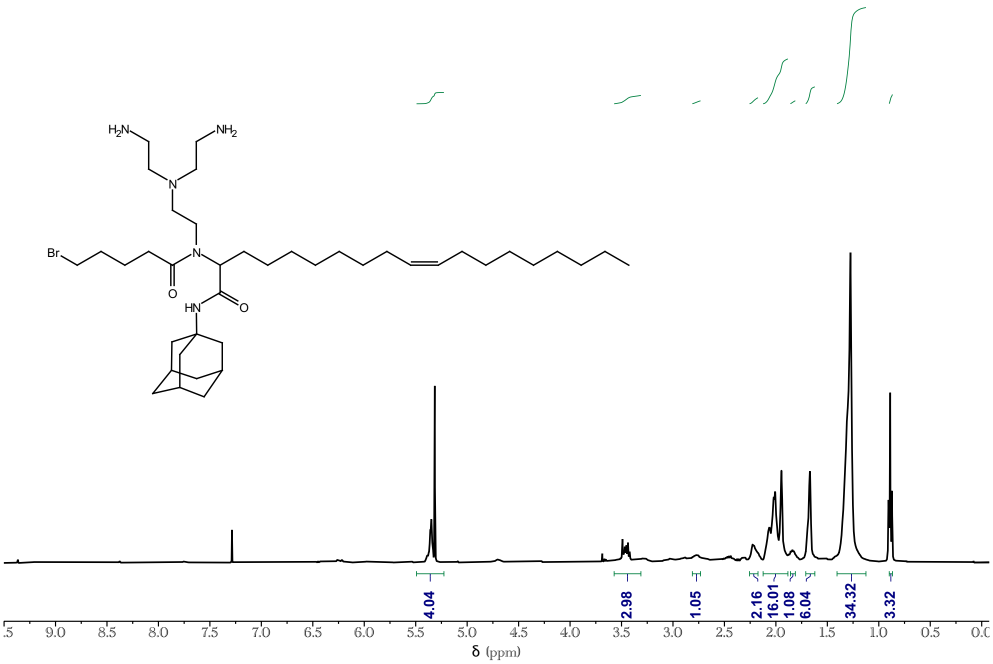

Supplementary Figure S31: $^1$H NMR for lipid LUMI-1.

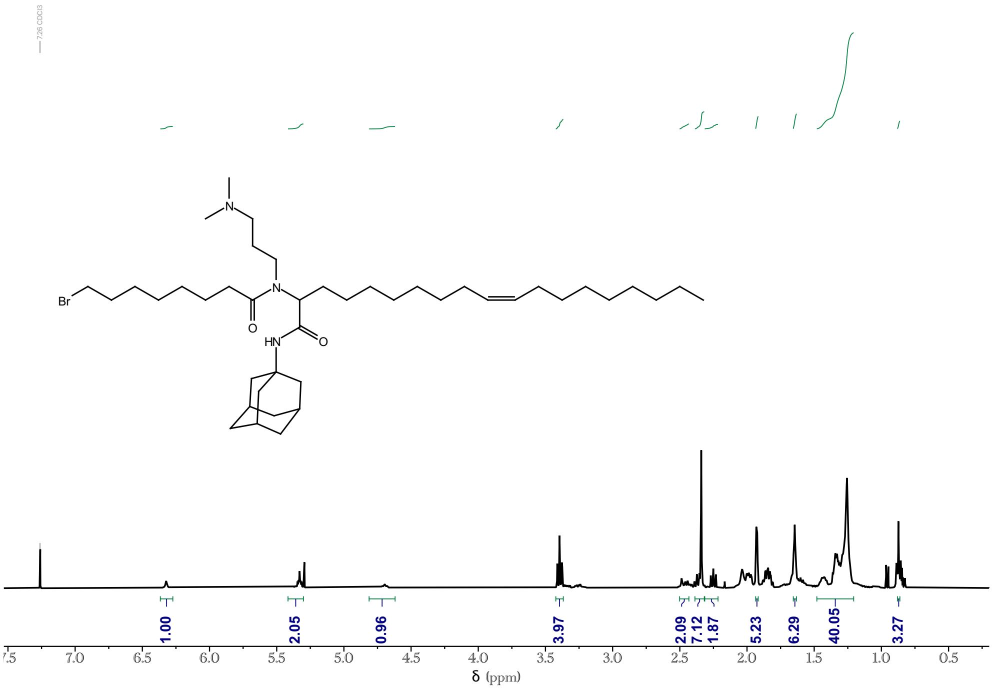

Supplementary Figure S32: $^{1}$H NMR for lipid LUMI-2.

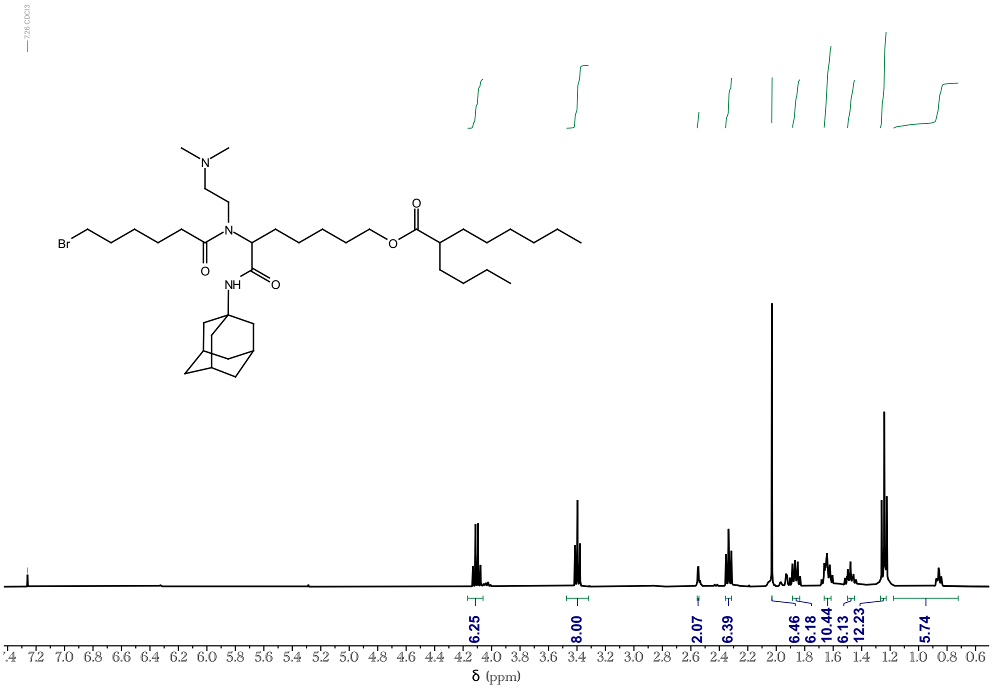

Supplementary Figure S33: $^{1}$H NMR for lipid LUMI-3.

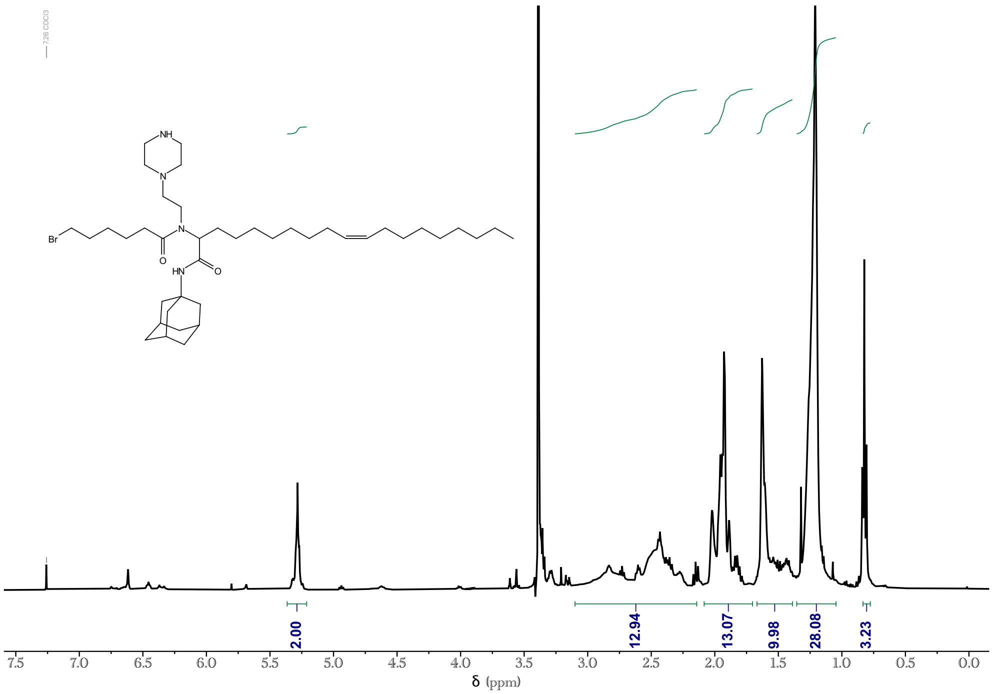

Supplementary Figure S34: $^1$H NMR for lipid LUMI-4.

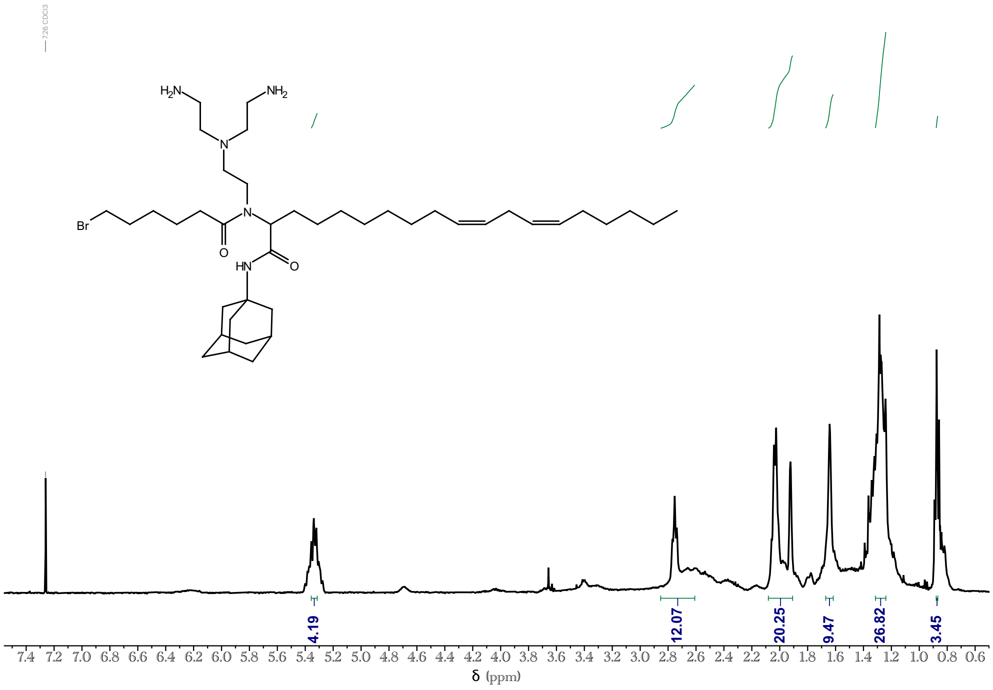

Supplementary Figure S35: $^1$H NMR for lipid LUMI-5.

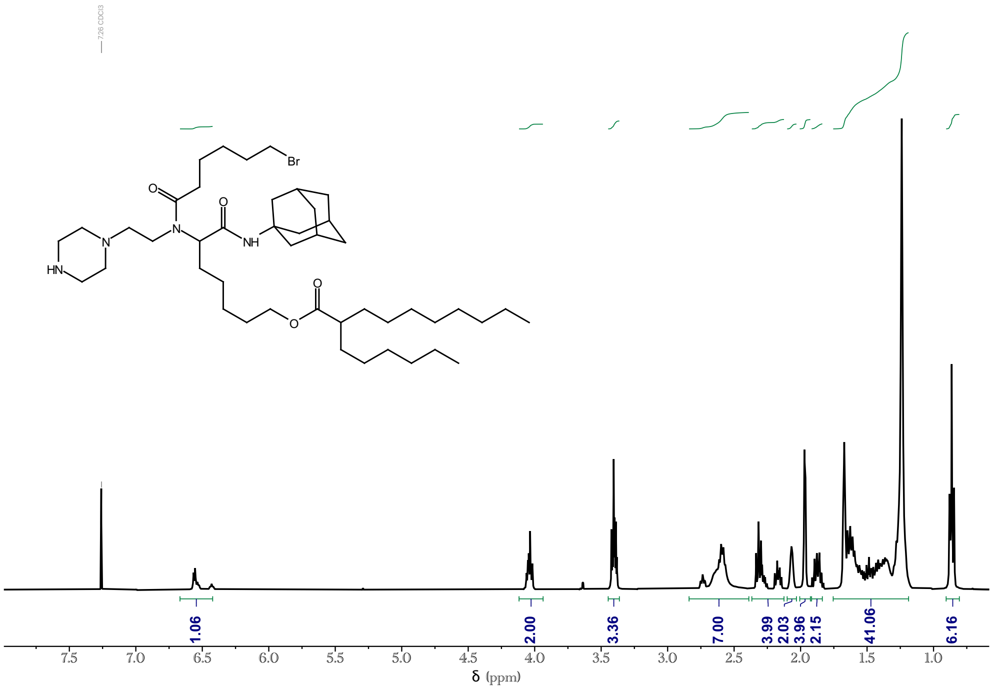

Supplementary Figure S36: $^{1}$H NMR for lipid LUMI-6.

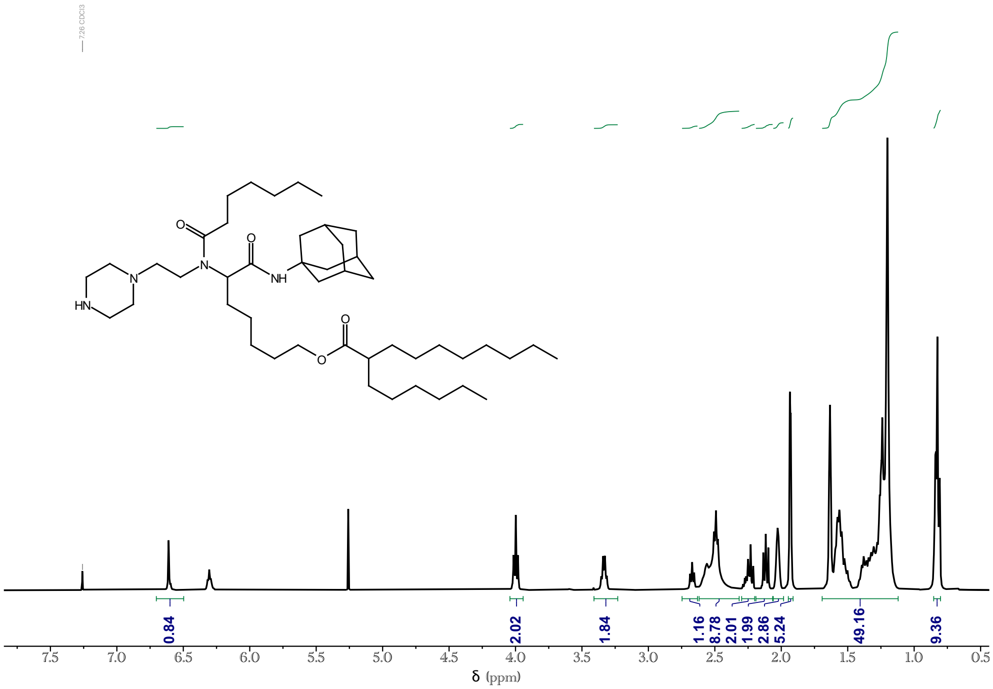

Supplementary Figure S37: $^{1}$H NMR for lipid LUMI-6D.
